# Cystathionine Beta-Synthase Promotes Anoikis Resistance and Transcoelomic Metastasis in Ovarian Cancer via SP1- ITGB1 regulation

**DOI:** 10.64898/2025.12.12.694000

**Authors:** Pallab Shaw, Arpan Dey Bhowmik, Akrit Pran Jaswal, Rameswari Velayutham, Srikanth Chiliveru, Samantha Ricketts, Chao Xu, Danny N. Dhanasekaran, Zhizhaung Joe Zhao, Resham Bhattacharya, Priyabrata Mukherjee, Shailendra Kumar Dhar Dwivedi, Geeta Rao

**Author notes:** Correspondence: Geeta Rao, PhD., Assistant Professor, Department of Pathology. OU Health Stephenson Cancer Center, The University of Oklahoma Health Campus, 941 Stanton L Young Blvd, BSEB#335, Oklahoma City, OK 73104, Oklahoma City, OK 73104, USA. Extension: 33680. **Conflict of interest**: The authors declare no potential conflicts of interest.

## Abstract

Anoikis resistance is crucial for ovarian cancer (OvCa) transcoelomic metastasis, during which exfoliated OvCa cells survive as spheroids before invading the omentum. Here, we demonstrate that cystathionine β-synthase (CBS), an H_2_S-producing transsulfuration pathway enzyme, is a key determinant of OvCa spheroidal viability and metastatic potential. Analysis of publicly available patient datasets, as well as an in-house tissue microarray, revealed that high CBS expression positively correlates with both poor progression-free survival and clinically observed peritoneal/omental metastasis. Integrated functional and proteomic analyses indicated that CBS silencing induces apoptosis in 2D monolayers. Consistent with this, CBS silencing in spheroids caused apoptosis and disrupted spheroid architecture. Mechanistically, this phenotype was associated with downregulation of oncogenic stemness and epithelial-mesenchymal transition. Further, through proteomic and bioinformatic analyses, we identified ITGB1 to be the hub protein in OvCa spheroidogenesis. Interestingly, knockdown of CBS led to abrogation of the ITGB1-mediated downstream pathway. Moreover, by proteomic and network analyses, we identified SP1 as a key transcriptional regulator of CBS-induced pro-spheroidal transcriptional programs of stemness and invasiveness. Stabilization of SP1 through persulfidation by H_2_S supplementation restored spheroidal viability and underlying protein signaling. Further, our results reveal that loss of CBS, through ITGB1 repression, disrupts spheroid architecture, leading to reduced metastatic docking on the murine omental surface *in vivo*. Collectively, these findings identify CBS as a central regulator of anoikis resistance and OvCa transcoelomic metastasis, highlighting the therapeutic potential of targeting CBS-dependent SP1-ITGB1 regulation to control metastatic spread.

## Introduction

Ovarian cancer (OvCa) is one of the most common and lethal gynecologic malignancies [1–3], largely due to advanced stage diagnosis and metastatic progression [4]. Early-stage OvCa, when the disease is confined to the ovaries, has an approximately 90% cure rate [5]. OvCa metastasis is largely transcoelomic, i.e., intraperitoneal [6]; due to the absence of an anatomic barrier, malignant cells from the ovaries exfoliate in the peritoneal cavity as either single cells or multicellular aggregates (MCAs) called spheroids [7–9]. These spheroids exhibit an anchorage-independent survival mechanism termed anoikis resistance (AR); normally, when cells detach from the extracellular matrix (ECM), they undergo detachment-induced programmed cell death called anoikis. However, cancer cells evade anoikis during metastasis, i.e., they become anoikis resistant [7, 10]. Habyan et al. compared exfoliated single cells and spheroids and showed that the spheroids exhibit greater AR [8]. Additionally, other reports suggest that single cells aggregate in the abdominal cavity to acquire AR, making spheroidogenesis a critical neoplastic attribute that facilitates OvCa metastasis [11]. These spheroids are enriched in cancer stem cells (CSCs) [12], which exhibit increased tumorigenicity, apoptosis resistance and metastatic potential [13]. Thus, targeting AR and the spheroidogenic potential of OvCa cells could be a promising avenue for effectively combating OvCa metastasis.

The literature suggests that apart from adhesion and stemness pathways, cellular metabolism and redox homeostasis play significant roles in regulating spheroid survival and AR [10]. Especially, transsulfuration pathway enzymes that facilitate cysteine, glutathione (GSH), and hydrogen sulfide (H_2_S) generation have recently been identified as important regulators of mitochondrial activity, survival signaling, and redox balance in cancer cells [14–17]. This pathway is catalyzed by cystathionine-γ-lyase (CSE) and cystathionine-β-synthase (CBS), with the latter found to exhibit profound pro-tumorigenic attributes [18]. High CBS expression in OvCa has been associated with neoplastic growth, chemoresistance and metastasis [19, 20]. While CBS-derived H_2_S aids mitochondrial respiration and ATP production, GSH, on the other hand, curbs oxidative stress, and thus the two act in consort to promote OvCa cell viability and tumor growth [19]. Loss of CBS was found to cause oxidative stress–mediated JNK activation and MFN2 degradation, leading to mitochondrial fragmentation and metabolic collapse. Therefore, CBS has been found to regulate mitochondrial morphogenesis [21]. In addition to this, CBS has also been found to exert control over transcriptional programs that mediate tumor progression. Chakraborty et al. (2015) reported that CBS regulates lipid metabolism in OvCa through a transcriptional axis involving SP1 and sterol regulatory element-binding proteins (SREBP1/2), thus aiding in lipogenesis, membrane biosynthesis, and invasive capacity [22]. This situates CBS as an upstream regulator of pro-oncogenic transcriptional networks in OvCa. Other reports regarding the role of CBS in OvCa demonstrate how it induces Nrf2-mediated oxidative stress response [23], xCT-mediated glutathione production [24] and ferroptosis resistance [25].

Cumulatively, this literature suggests that CBS plays a pleiotropic role in OvCa and is likely a candidate prognostic marker. Despite all this evidence linking CBS to chemoresistance and redox regulation, its direct role in regulating spheroid architecture and metastasis remains undefined.

Interestingly, our initial analysis of publicly available data suggested that high CBS expression correlates with reduced progression-free survival (PFS) of patients with advanced-stage OvCa characterized by transcoelomic metastasis through formation of spheroids that exhibit AR. In line with this, our analysis using an in-house OvCa patient tissue microarray (TMA) revealed a positive correlation between CBS expression in OvCa tissues and presence of omental metastasis. However, the mechanism by which CBS mediates trans-coelomic metastasis via spheroidogenesis and its role in AR remains unknown.

Considering that spheroid formation and AR play critical roles in OvCa metastasis, we investigated the role of CBS in promoting spheroidogenesis and AR. Our results clearly demonstrate that CBS enhances AR, stemness, and epithelial-mesenchymal transition (EMT) in 3D spheroids. These effects contribute to CD24⁺ cell enrichment within OvCa spheroids and are associated with the upregulation of ITGB1 and subsequent remodeling of spheroid histoarchitecture. Furthermore, CBS facilitates the ITGB1-FBN interaction, which is crucial for anchoring OvCa spheroids to the omental surface and initiating metastatic invasion. Specifically, these CBS-mediated effects are facilitated by H_2_S, which stabilizes the transcription factor, SP1, a central regulator of stemness-related genes and ITGB1. Cumulatively, our findings identify CBS as a promising therapeutic target for inhibiting AR and OvCa metastasis, enhancing treatment sensitivity, and ultimately improving patient outcomes.

## Materials and Methods

### Chemicals and reagents

Chemicals, reagents and antibodies used in this study are detailed in the Supplementary file S1 (Table S1 and Table S2).

### Animals

Six-week-old female Nu/J mice were procured from Jackson Laboratory and housed under controlled dark/light cycle and humidity/temperature in the OUHC-regulated animal facility. All animal studies were performed in accordance with guidelines and protocols approved by the OUHC Institutional Animal Care and Use Committee (IACUC) (Approval # 25-007-CH).

### Tumor microarray analysis

The TMA, comprising 109 OvCa patients along with 12 normal ovarian and 12 fallopian tube tissue sections, was accessed via the OU Health Stephenson Cancer Center (SCC) Tissue Pathology and Biospecimen Shared Resource. Immunohistochemical staining was done using CBS antibody (1:100) (Supplementary File S1; Table S1). H scores were obtained after analyzing the sections using the HALO: Quantitative Image Analysis for Pathology tool. H scores and sections to be considered were validated by a pathologist (R.V.). H score distribution by median with 95% CI among the cohort was generated using GraphPad Prism 10.0.1. A survival curve corresponding to gene expression of CBS in the TMA was generated using GraphPad Prism 10.0.1 as survival proportions vs. CBS H score. For metastasis vs. CBS H score correlation analysis, a clinician-provided dataset of patients with observed peritoneal/omental metastasis was analyzed (Supplementary File S1; Table S3). Analysis was performed using Logistic regression with a class-weighted method by assigning a binary score to denote presence or absence of metastasis, and was used to generate the AUC and odds ratio (probability/outcome) for CBS H score vs metastasis (binary) using GraphPad Prism 10.0.1. Further, to explore the association of CBS expression and tumor differentiation, the pathologist (RV) classified patients according to grade, where Grade 1 is well differentiated, Grade 2 moderately differentiated, and Grade 3 poorly differentiated tissue. The differences in CBS expression across tumor grades were evaluated using the Kruskal-Wallis test, followed by Dunn’s multiple comparisons test for pairwise analyses. Also, as a complementary analysis, pairwise comparisons were performed using Welch’s t-test, which does not assume equal variances. All tests were two-sided, and statistical significance was defined as p < 0.05.

### Bioinformatic analysis

Detailed protocols for the bioinformatics analyses conducted in this manuscript have been provided in Section 1 of the Methods provided in Supplementary File S1.

### Cell culture

FTE187, FTE188, OSE, HOSE, OVCAR8, OVCAR4, TYK-nu, TYK-nu.CP-r, A2780-CP20, COV362, COV318 and OVCAR3 were cultured as specified in Supplementary File S1; Table S4. All cell lines were authenticated and tested for the absence of Mycoplasma contamination at the SCC Cell & Tissue Analysis Core.

### Proteomics analysis of OvCa monolayers and spheroids

Raw data for this study were generated at the IDeA National Resource for Quantitative Proteomics, AR, USA. Proteomic profiling of both monolayers and spheroids (CBS-expressing and CBS-silenced COV318 cultures) was performed. The MetaboAnalyst, Enrichr, ExpressAnalyst, and Harmonizome platforms were used to analyze normalized datasets to identify transcription factor networks, enriched pathways, and differentially expressed proteins. The NCI Nature 2016 gene set, MSigDB Hallmark, and Cancer Hallmark gene sets were used for the over-representation analyses. See Methods section 2 of Supplementary file S1 for a detailed workflow.

### Phalloidin staining

4% paraformaldehyde-fixed 2D monolayers were stained with CytoPainter Phalloidin-iFluor 647 Reagent according to the manufacturer’s protocol (for details, refer to Section 3 of Methods in Supplementary file S1).

### Spheroid formation

For spheroid formation, cells grown on 2D monolayers were transferred to poly-HEMA-coated 60 mm plates and incubated for 48 h, after which they were dissolved into single cell suspensions that were in turn used for siRNA transfection for 96 h. For rescue experiments with GYY4137, spheroids were treated with 1 mM GYY4137 for the final 24 h, i.e., 72 h after knockdown. (for details, refer to Section 4 of Methods Supplementary file S1).

### 3D Cell viability assay

The Cell Counting Kit 8/ CCK-8 Assay Kit/ WST-8 Assay Kit, no-wash, mix-and-read, cell viability assay was used to assess the viability of spheroids according to the manufacturer’s protocol (for details, refer to Section 5 of Methods in Supplementary file S1).

### Anoikis assay

Spheroids prepared as described above were assessed to determine live/dead cell ratio. 96 h after siRNA transfection, the spheroids were stained with 2 mM Calcein-AM and 4 mM ethidium-homodimer for 30 min before imaging with a Nikon Eclipse Ti2E inverted fluorescence microscope. Composite images were constructed using Z-stack images. The quantification of fluorescence intensity was conducted utilizing ImageJ Fiji software. The corrected spheroidal fluorescence intensity was calculated separately for FITC (Live) and TRITC (Dead) filters as integrated intensity/total spheroid area.

### Western blot analysis

Spheroidal lysates corresponding to 15 µg proteins from each sample were subjected to Western blotting. Blots were incubated with appropriate primary antibodies, followed by corresponding secondary antibodies. The bands were developed with Clarity™ Western ECL Substrate or Super Signal West Atto and visualized using the Bio-Rad Chemidoc™ Imaging System (Refer to Section 6 of Methods in Supplementary file S1).

### Histology and Immunofluorescence

Spheroids were collected in 1.5 mL microfuge tubes, pelleted at 500 rpm for 2 min, washed in PBS, and fixed in 4% paraformaldehyde for 20 min. After a further PBS wash, spheroids were treated with chilled 70% ethanol. Following dehydration through graded alcohol, the spheroids were embedded in paraffin. 5 µm thick tissue sections were stained with hematoxylin-eosin (HE) following standard protocols and viewed under a light microscope. Briefly, slides with sections of 5 µm thickness were deparaffinized using xylene for 5 min, passed through graded alcohol concentrations, washed in running tap water for 10 min and subjected to antigen retrieval before blocking with 2.5% horse serum. The sections were microscopically examined after incubation with appropriate primary antibodies overnight, followed by incubation with the secondary antibody and mounting (for details, refer to Section 8 of Methods in Supplementary file S1).

The protocol for immunofluorescence from patient-derived ascites samples is provided in detail in Section 9 of Methods provided in Supplementary file S1.

### qRT-PCR analysis

cDNA synthesized using total RNA from spheroids was subjected to qRT-PCR. The 2^-ΔΔCT^ method was used to calculate relative mRNA expression after normalizing it to 36B4. Supplementary file S1 Methods section 7 contains comprehensive primer sequences and reaction conditions.

### CD24 staining and flow cytometry

APC-conjugated CD24 antibody and Ghost Dye™ Violet 510 were used to stain single-cell suspensions made from dissociated spheroids for viability and flow cytometry analysis performed using a Stratedigm-4 cytometer. Details of the staining procedure and gating strategy are provided in Methods section 10 and Fig. S24 of Supplementary file S1.

### Preclinical model of OvCa transcoelomic metastasis

A fluorescently labeled (Invitrogen molecular Probes CellTracker Red CMTPX Dye) single cell suspension derived from spheroids was injected into 6-week-old Nu/J mice, 4h after which the omentum was dissected out and examined under the microscope (Fig. 6A). For a detailed protocol, see Methods section 11 of the Supplementary file S1.

### Fibronectin attachment assay

CBS-expressing and silenced spheroids were generated with anoikis-resistant enriched OvCa cells as described above under Spheroid Formation. After 72 h, siCBS spheroids for rescue experiments were treated with 1mM GYY4137. After 94 h of siRNA treatment, spheroids were collected in tubes, dissolved into single cell suspensions by mild trypsinization (2 min) and counted for cell viability. For each group, equal numbers of viable cells, labelled with Invitrogen molecular Probes CellTracker Red CMTPX Dye, were plated on FBN-coated 96-well plates (2000 cells/well). After 1 or 2h incubation, media were aspirated, and wells were washed with PBS before visualization by microscopy. For quantification of cells attached to FBN-coated wells, 16-bit images of whole wells were procured by the ‘acquire stitched image’ option in NIS-Elements Advanced Research software using the Nikon Eclipse Ti2E inverted fluorescence microscope. These images were analyzed using ImageJ Fiji software to count the number of cells.

### In silico identification of SP1 binding sites in the ITGB1 promoter

We used the Eukaryotic Promoter Database (EPD) (https://epd.expasy.org/cgi-bin/epd/get_doc?db=hgEpdNew&format=genome&entry=ITGB1_1) to obtain the human ITGB1 promoter sequence (−2000 to +100 bp relative to the transcription start site, TSS). Next, to predict putative SP1 binding sites, we used the JASPAR database (https://jaspar.elixir.no/). The detailed protocol is provided in Methods section 12 of Supplementary file S1.

### CBS overexpression and SP1 inhibition assay

OVCAR8 cells were used to assess SP1-dependent regulation of ITGB1. Cells were transiently transfected with either empty vector or CBS expression plasmid (FLAG-CBS-pcDNA3 plasmid) for 24 h, following which they were subjected to mithramycin treatment (25 nM) for another 24 h. Whole-cell lysates were subjected to Western blot analysis.

### Chromatin immunoprecipitation (ChIP) and qPCR

Based on the spatial positioning of predicted binding sites, six primer sets were made to amplify areas that are rich in SP1 binding motifs (Fig. 7I). Briefly, chromatin was sheared to ⁓200-500 bp fragments in microTUBE-130 tubes using Covaris E220 ultra-sonicator (Conditions: peak power: 105.0, duty factor: 5.0, cycles/burst 200, and duration 120 sec). ChIP assays were performed using the Magna ChIP™ A/G Chromatin Immunoprecipitation Kit (Millipore, 17-10085) following the manufacturer’s protocol. The detailed protocol is provided in Methods section 13 of Supplementary file S1.

### Modified biotin-switch assay

To assess SP1 persulfidation in OvCa cells, OVCAR8 cells were transfected with siCon or siCBS and cultured for 96 h. For rescue experiments, CBS-silenced cells were supplemented with GYY4137 (1 mM) for the final 24 h (i.e., after 72 h of knockdown). Cell lysates were prepared and subjected to the modified biotin-switch protocol as described in Supplementary File S1, Method section 14. Briefly, free thiols were blocked, and persulfidated cysteine residues were selectively labeled with Biotin HDDP for subsequent pull-down. For negative control, a parallel set of samples was treated with DTT to eliminate persulfidation signals. Biotinylated proteins were pulled down and analyzed by western blotting for SP1 in the pull-down fraction. Input lysates were immunoblotted for SP1, GAPDH, and α-tubulin, with GAPDH serving as a positive control for persulfidation.

### Statistical Analysis

All analyses were performed in triplicate for three independent experiments, except for the Mass spectrometry-based proteomics, where three biological replicates were used. The results were expressed as mean ± SEM. Significance (P < 0.05) was determined by one-way ANOVA, two-way ANOVA, and two-tailed paired t tests, unless otherwise noted. Class-weighted logistic regression was used for AUC and ROC analysis. The Mann-Whitney test (P < 0.05) was used for TNM plot analysis. Mantel-Cox log-rank and Gehan-Breslow-Wilcoxon tests were used for survival curve analysis with error bars indicating 95% confidence interval (CI).

## Results

### High CBS expression is positively correlated with OvCa metastasis

We analyzed a combined cohort of 764 patients (1,364 samples) from four major studies [Ovarian Cancer (CPTAC, GDC), Ovarian Cancer (Gray Foundation, *Cancer Discovery* 2024), Ovarian Cancer - MSK SPECTRUM (MSK, *Nature* 2022), and High-Grade Serous Ovarian Cancer (TCGA, GDC)] on the cBioPortal platform, and observed a progressive decline in the alive-to-dead patient ratio with advancing FIGO stage (Fig. S1A). Differential expression analysis of CBS in OvCa using the ‘TN plot: compare tumor and normal’ analysis tool revealed CBS to be significantly upregulated in OvCa (P<0.0001) (Fig. 1A). We also analyzed the expression of CBS across OvCa FIGO stages from a publicly available dataset (CPTAC, GDC) on cBioPortal, which indicated that the median expression of CBS is significantly higher in IIIB, IIIC and IV than in earlier stages (Fig. 1B). Next, we generated Kaplan-Meier survival plots, synthesized from the microarray dataset, for CBS (Affymetrix ID: 212816_s_at) in a total of 1232 high-grade serous OvCa (HGSOC) patients at different FIGO stages (Stage I: 107; Stage II: 72; Stage III: 1079 and Stage IV: 189 patients) revealing significant correlation between CBS expression and PFS of HGSOC patients. Interestingly, the difference in median PFS between the low expression cohort and high expression cohort increased with FIGO stage (Fig. 1C). These data suggest that elevated CBS expression associates with advanced disease stage, poor PFS, and metastatic burden in OvCa. Additionally, we also investigated the overall survival (OS) in these patients as a function of CBS expression. This stage-stratified survival analysis revealed that while high CBS expression was significantly associated with reduced PFS in Stage 1 (HR = 2.93, p = 0.037), Stage 3 (HR = 1.21, p = 0.02), and Stage 4 (HR = 1.49, p = 0.045) tumors (Fig. 1C), OS was significantly affected by high CBS only in Stage 4 disease (HR = 1.46, p = 0.04), whereas earlier stages showed no statistically significant impact on OS (Supplementary File S1; Fig. S1C). These findings suggest that CBS predominantly contributes to disease progression, with its effect on OS becoming evident in advanced-stage OvCa. To validate these findings, we evaluated the expression of CBS in our in-house OvCa TMA (Fig.1D) comprising 109 HGSOC patients by immunohistochemical analysis. A pathologist (RV) assigned H scores to all 109 patients (Fig. 1E); upon analyzing the survival proportion of these 109 patients, we found that the survival probability was inversely proportional to CBS H score levels (Fig. 1F). To further corroborate the role of CBS in transcoelomic metastasis, we analyzed our TMA cohort with a CBS H-score and observed metastasis into the peritoneum/omentum. The data revealed significantly higher CBS in terms of H scores in the metastatic cohort in comparison to the non-metastatic cohort (P value <0.0001) (Fig. 1G). Further analysis using class-weighted logistic regression revealed that there is a positive correlation between CBS expression and presence of metastasis in OvCa patients in our TMA cohort (AUC=0.675) (Fig. 1H and Fig.1I). These results suggest that CBS plays a critical role in the metastatic process of OvCa. In addition to this, we found no statistically significant association between the expression of CBS and tumor grades by non-parametric methods (Kruskal-Wallis test), implying that the expression distributions of Grade 1, Grade 2 and Grade 3 tumors were not consistently different. However, Welch’s t-test showed a significant difference in the means of Grade 1 vs. Grade 2 and Grade 1 vs. Grade 3 (Supplementary File S1; Fig. S1B). Notably, the reliability of mean-based inference is limited by the small sample size for Grade 1 (n = 3), therefore these results should be interpreted cautiously. Nevertheless, there was a trend of increased CBS expression in higher grade tumors, suggesting a potential association with tumor aggressiveness and dedifferentiation. This pattern might become more evident with more balanced cohort(s) and is consistent with our findings linking CBS with stemness, EMT and metastatic potential.

**Fig. 1.**
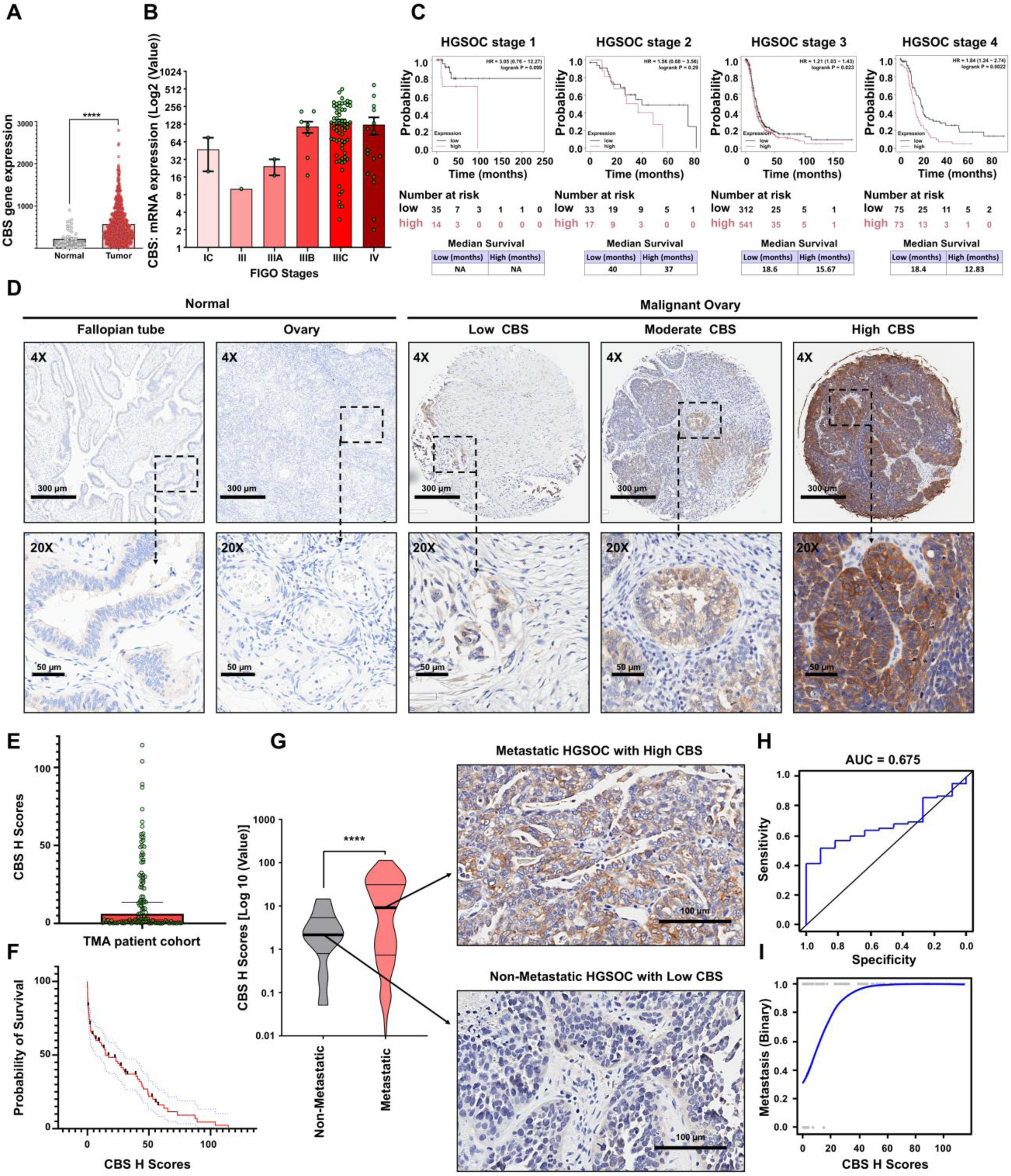
High CBS expression in OvCa correlates with advanced disease stage, poor prognosis, and metastasis in patient cohorts. (A) Differential expression analysis of CBS using the TNM plot tool (tumor vs. normal) shows significant upregulation of CBS in OvCa. (B) CBS expression across FIGO stages (CPTAC cohort) demonstrates higher median expression in advanced-stage (IIIB-IV) disease. (C) Kaplan-Meier survival analysis of HGSOC patients (N = 1232; Stage I: 107, Stage II: 72, Stage III: 1079, Stage IV: 189) shows that higher CBS expression is associated with significantly reduced PFS. (D) Representative IHC images of CBS expression in normal fallopian tube, ovary, and OvCa tissues from the in-house TMA cohort (n = 109), showing low, moderate, and high expression. (E) Distribution of CBS H-scores across the OvCa TMA cohort (N = 109). (F) Survival analysis of the TMA cohort (N = 109; death events = 73, censored = 36) indicating an inverse relationship between CBS H-score and patient survival probability. (G) Comparison of CBS H-scores between non-metastatic (n = 12) and peritoneal/omental metastatic tumors (n = 97) within the TMA cohort, reflecting the predominance of advanced-stage cases typical of OvCa presentation. (H) Receiver operating characteristic (ROC) curve for CBS H-score versus observed metastasis in the TMA cohort (N = 109, AUC = 0.675). (I) Class-weighted logistic regression analysis depicting the association between CBS H-score and observed metastasis in OvCa patients from the TMA cohort (N = 109, AUC = 0.675).

### CBS expression in OvCa cell lines is positively correlated with expression of stemness factors

Since spheroidogenesis is a prerequisite for transcoelomic dissemination in OvCa, we next evaluated whether CBS expression is correlated with the transcription factors that drive this phenomenon. Robinson et al. have shown that Sox2, Oct4 and Nanog are consistently upregulated in OvCa cells grown in anchorage-independent conditions and associate with tumor-initiating potential [26]. The keyword “ovarian cancer spheroids” yielded 3,947 genes in a GeneCards search [27]. Analysis of this gene set using the transcription factor analysis module

(CHEA 2022) in the Enrichr tool revealed Oct4-, Sox2- and Nanog-regulated transcriptional programs among the top 10 enriched programs (Supplementary File S1; Fig. S2A). Immunoblotting of these three transcription factors in both cancer and normal cells revealed a positive correlation with CBS protein levels (Fig. 2A). High CBS level in OvCa was validated since we found high CBS expression in human OvCa cell lines (OVCAR8, OVCAR4, A2780-CP20, COV318, Tyknu CPR, COV362 and OVCAR3) compared to normal human fallopian tube epithelium and ovarian surface epithelium cell lines (i.e., FTE187 and FTE188, and OSE and HOSE respectively) (Fig. 2A). We also determined the status of CBS expression in 2D monolayers and 3D spheroids of COV318, a representative ascites-derived HGSOC cell line, and found that CBS is upregulated in the 3D spheroids compared to 2D culture (Supplementary File 1; Fig. S2 B). These findings implicate CBS as a driver of spheroidogenic capacity, anchorage-independent survival and intraperitoneal metastasis in OvCa.

**Fig. 2.**
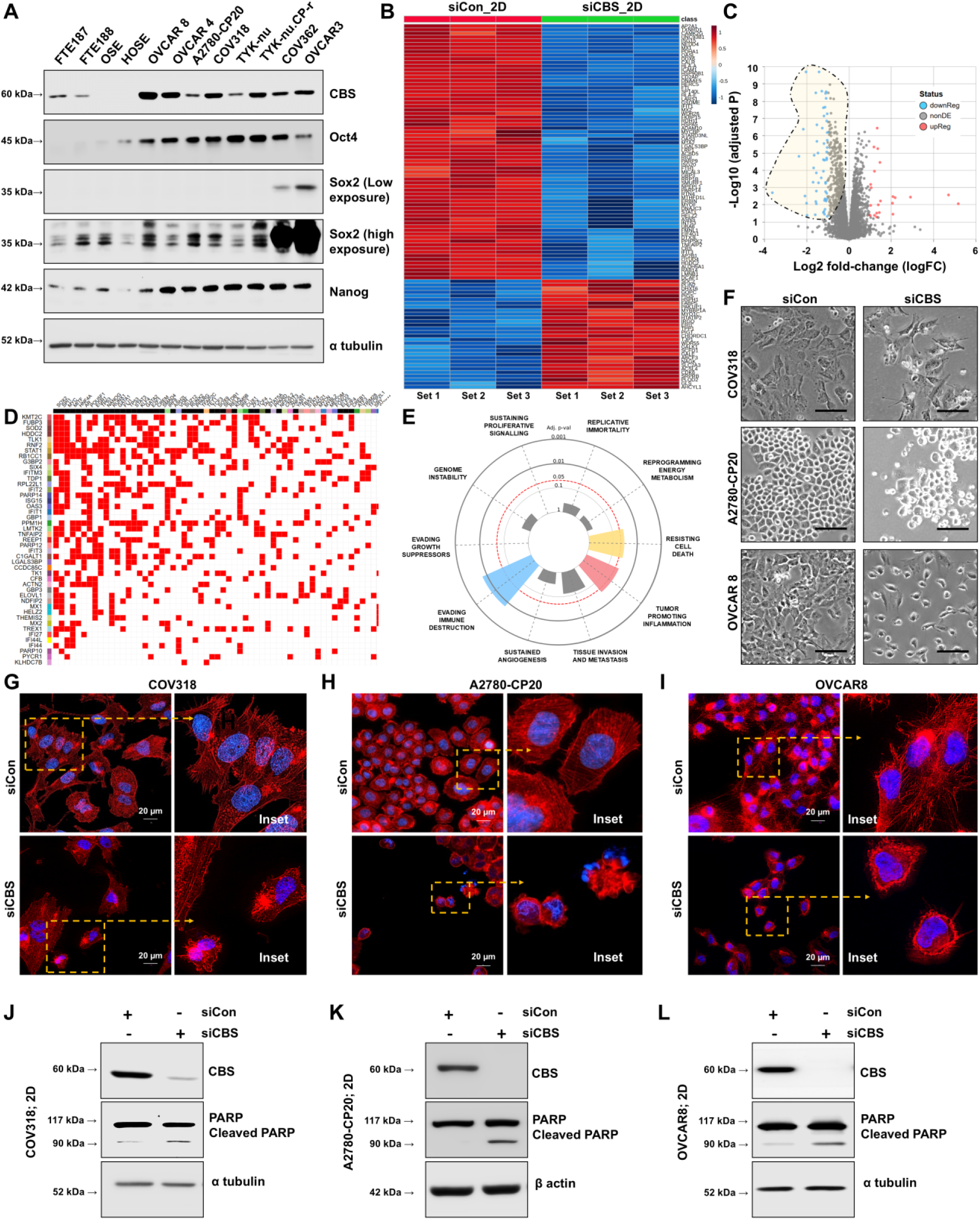
CBS expression correlates with pro-spheroidogenic transcription factors and promotes OvCa survival. (A) Immunoblot analysis of CBS and stemness factors (Oct4, Sox2 and Nanog) in human OvCa (OVCAR8, OVCAR4, A2780-CP20, COV318, Tyknu CPR, COV362, OVCAR3), normal human fallopian tube epithelium (FTE187, FTE188), and ovarian surface epithelium (OSE, HOSE) cell lines. (B) Heat map representing the top 100 DEGs upon proteomic analysis of siCon vs siCBS 2D monolayers. (C) Volcano plot showing significantly upregulated and downregulated DEGs (log2FC=1.05, p=0.05). (D) Transcription factor enrichment analysis of CBS-dependent DEPs identifies Oct4, Sox2, and Nanog among the top regulators. Clustergram shows Sox2, Oct4 and Nanog among the top 50 enriched transcription factors. (E) ORA with Cancer Hallmark gene set using significantly downregulated DEGs. (F) Photomicrographs showing morphological alteration upon CBS knockdown in COV318, A2780-CP20, and OVCAR8 monolayers. (G, H and I) Phalloidin red staining of COV318, A2780-CP20, and OVCAR8 monolayers, respectively, demonstrating hallmarks of apoptosis, including membrane blebbing, cell shrinkage, and disorientation of F-actin. (J, K and L) Immunoblots confirming apoptosis via detection of cleavage of PARP1 into the 89 kDa fragment in CBS-silenced cells. All experiments were performed in three independent biological replicates (n = 3).

For mechanistic studies, we used COV318, A2780-CP20 and OVCAR8, each representing distinct disease features. COV318 cells, derived from the ascites of a HGSOC patient, represent an *in vitro* model of AR. A2780-CP20 cells are a cisplatin-resistant subline of A2780, itself an endometrioid OvCa cell line, and represent a chemoresistant epithelial OvCa variant. OVCAR8 cells have a unique ability to form ascites *in vivo* and thus represent HGSOC [28].

For proteomic analysis to demonstrate the role of CBS in OvCa cell monolayers (2D), we used COV318, i.e., the ascites-derived HGSOC cell line. The data showing significant differentially expressed genes (DEGs), is represented as a heat map (Fig. 2B) and volcano plot (Fig. 2C). CBS-expressing and -silenced COV318 monolayers exhibited distinct proteomic profiles as evident from PCA (PLSDA) analysis (PC1=46.21% and PC2=33.8%) (Supplementary File S1; Fig. S3). Transcription factor enrichment analysis of the downregulated DEGs revealed Oct4-, Sox2-, and Nanog-associated transcriptional programs (Fig. 2D), suggesting that stemness-associated regulatory networks contribute to CBS-dependent gene expression in OvCa monolayers.

### CBS knockdown promotes apoptosis in 2D culture of OvCa cells

To further delineate the role of CBS in OvCa progression, we analyzed the proteomic profile of CBS-expressing and CBS-silenced OvCa monolayers. Overrepresentation analysis (ORA) with the CBS-knockdown associated downregulated DEGs (in 2D siCBS) using the Cancer Hallmark gene set revealed ‘resisting cell death’, ‘evading immune destruction’ and ‘genome instability’ as the most enriched pathways, implying that CBS regulates these oncogenic hallmarks (Fig.2E). siRNA-mediated knockdown of CBS in COV318, A2780-CP20 and OVCAR8 monolayers showed reduction in cell density accompanied by increased cell rounding, a detached-from-substratum morphology (Fig. 2F, Supplementary File S1; Fig. S4) and indicator of cell death. Further, F-actin staining using phalloidin red revealed a fine meshwork of short actin filaments in the cytoplasm or clusters of actin aggregates in the central part of the control cells, while CBS-silenced cells exhibited thick bundles of actin filaments, i.e., stress fibers near the plasma membrane (Fig. 2G, H, I, Supplementary File S1; Fig. S5, S6, S7). Other morphological hallmarks, including membrane blebbing and cell shrinkage, were evident in cells with CBS knockdown. To further confirm apoptosis by CBS knockdown, we performed immunoblotting, which showed robust induction of cleaved PARP1, a hallmark of apoptosis, in the CBS knockdown condition as compared to the control group (Fig. 2J-L, Supplementary Files S1; Fig. S8). These results suggest that CBS promotes the survival of OvCa cells and is associated with OvCa progression.

### CBS is crucial for anchorage-independent survival and growth of OvCa cells

Next, to evaluate the potential role of CBS in promoting anchorage-independent survival (spheroid growth) and AR in OvCa, we grew OvCa cells in low-attachment plates for 48h (to enrich anoikis-resistant cells), followed by CBS knockdown (Fig. 3A). CBS-silenced spheroids showed a marked reduction in anchorage-independent cell viability in comparison to CBS-expressing spheroids, implying a role for CBS in endorsing AR (Fig. 3B, Supplementary File S1; Fig. S9). CBS silencing reduced viability to 43.03 ± 1.03% in COV318, 36.83 ± 1% in A2780-CP20, and 39.7 ± 0.59% in 3D spheroids compared to their respective control (Fig. 3C).

**Fig. 3.**
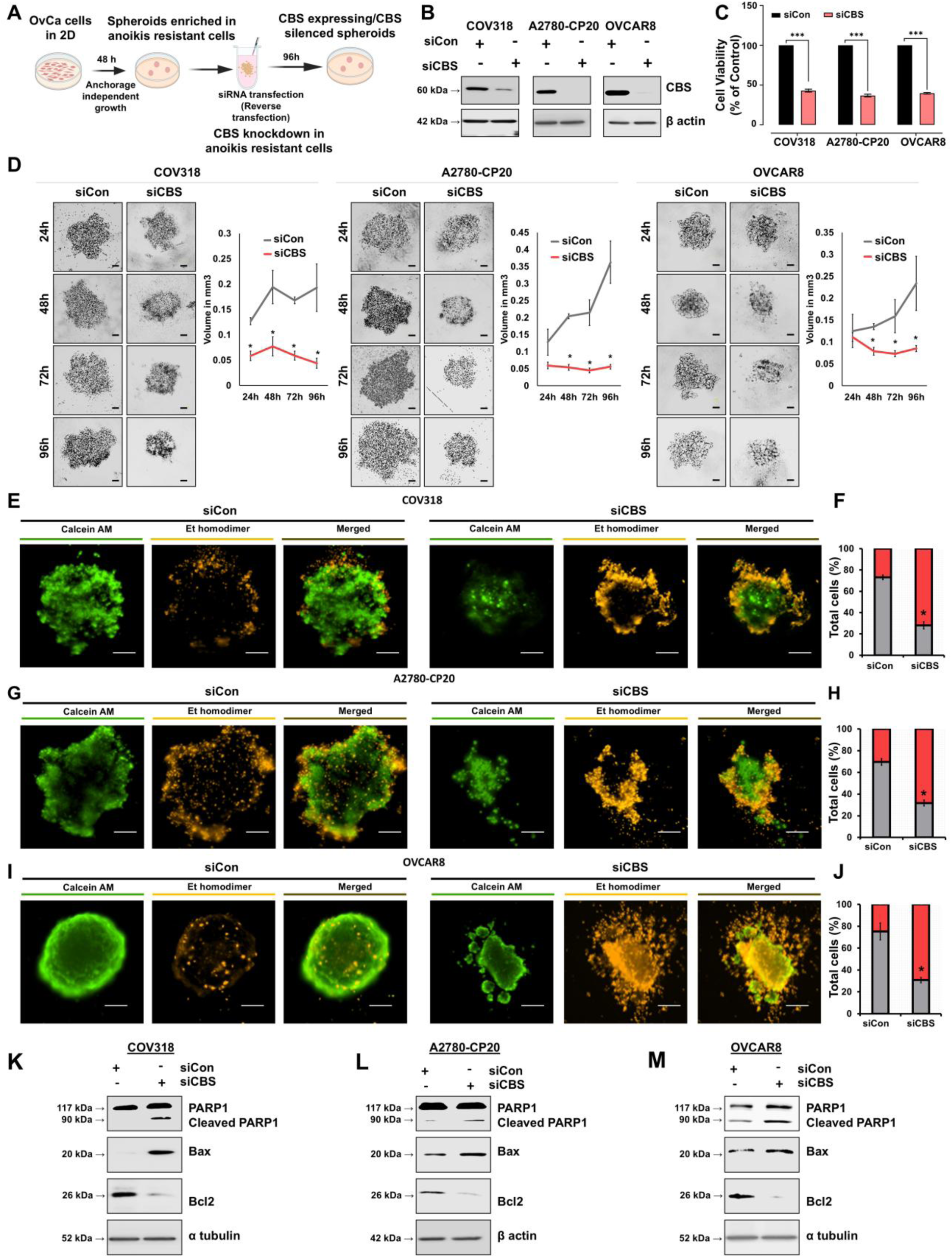
CBS supports anchorage-independent survival of OvCa spheroids and protects against anoikis. (A) Schematic representation of experimental workflow used to knockdown CBS in anoikis-resistant spheroidal cells. (B) Immunoblots confirming siRNA-mediated CBS knockdown in anchorage-independent spheroids of COV318, A2780-CP20 and OVCAR8. (C) Reduced cell viability after CBS silencing in COV318 (P value= 0.0004), A2780-CP20 (P value= 0.0003), and OVCAR8 spheroids (P value<0.0001). (D) Microscopic examination, from 24h after transfection through 96h of 3D culture, showing CBS-expressing spheroids increase in size, whereas CBS-depleted spheroids fail to grow. (E-J) Calcein AM-Ethidium homodimer dual staining of CBS-expressing and CBS-silenced OvCa spheroids, demonstrating loss of viability in CBS-silenced spheroids (Scale bars =200 µm) along with their quantification plots showing dead-to-live cell ratio. (K-M) Immunoblot analysis showed upregulation of Bax and cleaved PARP1 with concomitant downregulation of Bcl-2 in CBS-depleted spheroids. All experiments were performed in three independent biological replicates (n = 3).

Microscopic examination of spheroids starting immediately after transfection and continuing through 96h, showed that CBS-silenced spheroids exhibited reduced size compared to their respective controls (Fig. 3D) further establishing the role of CBS in anchorage independent growth. Immunofluorescence-based detection of CBS in CBS-silenced spheroids showed overall downregulation of CBS and reduced intracellular levels (Supplementary File S1; Fig. S10A, S10B, S10C). The assessment of dead cell-to-live cell ratio of these spheroids through Calcein AM-Ethidium homodimer dual staining indicated significant increases in the dead cell percentage (Fig. 3E-J) (Supplementary File S1; Fig. S11) within spheroids, further indicating triggering of anoikis. This was confirmed by the cleaved PARP1, increase in Bax along with reduced Bcl2 expression (Fig. 3K, 3L, 3M) (Supplementary File S1; Fig. S12). These findings suggest that CBS plays a key role in anchorage-independent survival and AR in OvCa.

### CBS promotes cancer stemness and epithelial-mesenchymal transition in OvCa spheroids

Spheroidogenesis, i.e., formation of MCAs, is critical for OvCa cells to exhibit anchorage-independent survival in the peritoneal cavity. As OvCa spheroidogenesis, AR and metastasis are associated with neoplastic stemness [29], we investigated the stemness circuitry of spheroids grown under control and CBS-silenced conditions. Immunoblotting-based expression profiling indicated that CBS knockdown resulted in the downregulation of CD44, along with the stemness transcription factors Oct4, KLF4, and Nanog in the spheroids of COV318 (Fig. 4A), A2780-CP20 (Fig. 4B) and OVCAR8 (Fig. 4C, Supplementary File S1; Fig. S13). Sox2 was significantly downregulated in COV318 (Fig. 4A) and A2780-CP20 (Fig. 4B) but not in OVCAR8 spheroids (Fig. 4C, Supplementary File S1; Fig. S13), suggesting cell line-based variation in stemness factor expression.

**Fig. 4.**
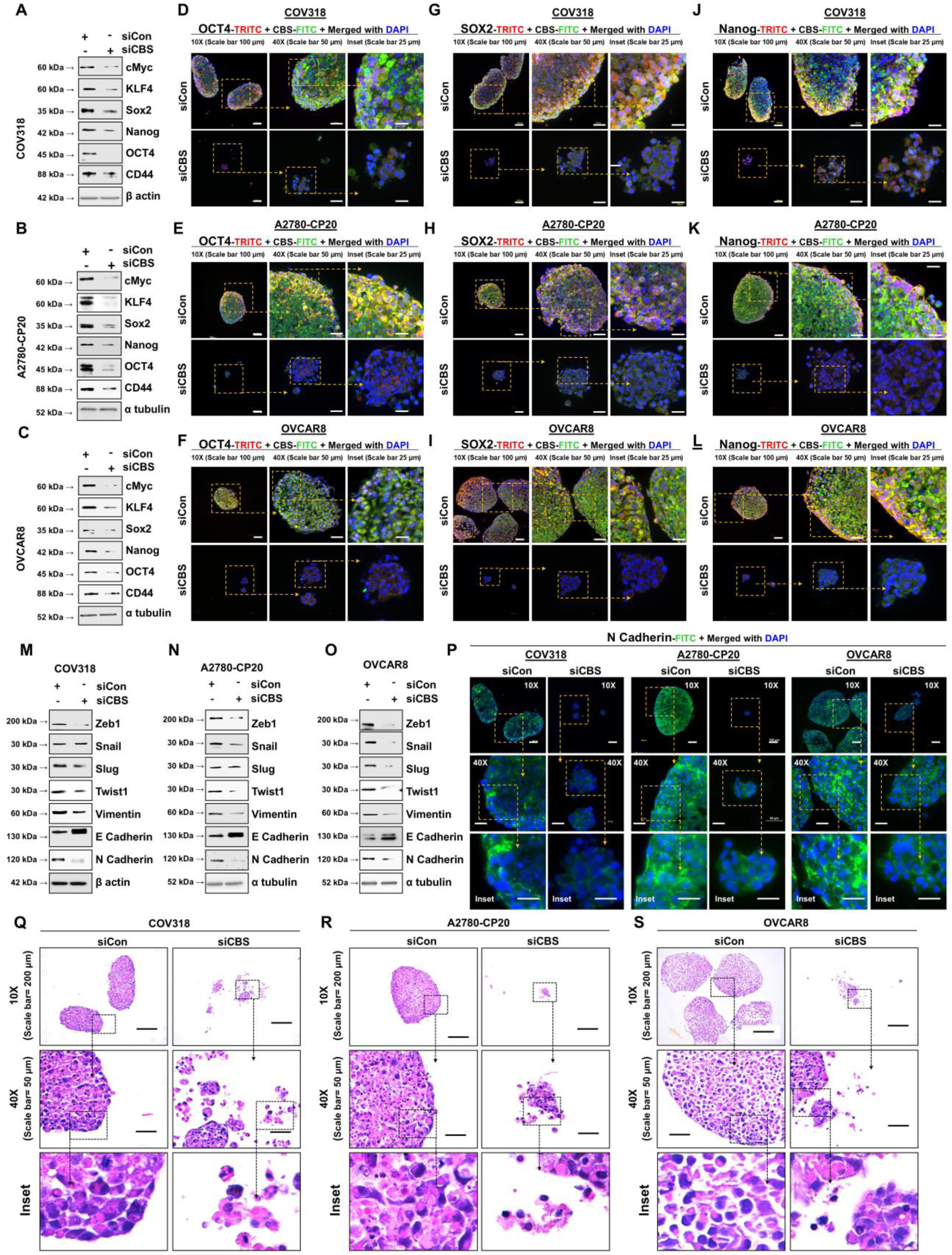
CBS promotes stemness circuitry and EMT programs in anoikis-resistant OvCa spheroids. Immunoblotting of COV318 (A), A2780-CP20 (B), and OVCAR8 (C) spheroid-lysates shows that CBS knockdown reduced CD44 along with the stemness-associated transcription factors Oct4, Sox2, KLF4, and Nanog. (D-L) Immunofluorescence analysis of spheroids showing co-expression of CBS (green) with stemness-associated transcription factors Oct4, Sox2, and Nanog (red) in COV318 (D, G, J), A2780-CP20 (E, H, K), and OVCAR8 (F, I, L) cells. (M, N and O) Western blots showing CBS depletion causing downregulation of EMT transcription factors, Zeb1, Snail, Slug, and Twist1, accompanied by upregulation of the epithelial marker E-cadherin and downregulation of the mesenchymal markers N-cadherin and vimentin in all three cell lines, indicating suppression of the mesenchymal phenotype. (P) N-cadherin localization by immunofluorescence analysis showing N-cadherin enrichment at spheroidal cell-cell junctions in control spheroids, and its decrease upon CBS knockdown. (Q, R and S) Photomicrographs of H & E-stained OvCa spheroids; FFPE section histoarchitectural analysis of OvCa spheroids following H & E staining demonstrates cohesive spheroids in CBS-expressing spheroids but loose spheroids with CBS knockdown. All experiments were performed in three independent biological replicates (n = 3).

To further determine whether CBS expression is associated with stemness-enriched cell populations within spheroids, we performed IF analysis in 3D spheroids generated from COV318, A2780-CP20, and OVCAR8 cells. In control spheroids, Oct4 (TRITC/red) (Fig. 4D, 4E, 4F, Supplementary File S1; Fig. S16), Sox2 (TRITC/red) (Fig. 4G, 4H, 4I, Supplementary File S1; Fig. S17), and Nanog (TRITC/red) (Fig. 4J, 4K, 4L, Supplementary File S1; Fig. S18) were observed with strong CBS expression (FITC/green). Merged images showed substantial overlap between CBS and stemness marker-positive cells, observed as a yellow/orange signal, suggesting that CBS-expressing cells are enriched within stemness-associated populations under anchorage-independent conditions. Notably, while Oct4 and Nanog showed significant nuclear localization, Sox2 showed a less defined nuclear pattern in CBS-expressing spheroids. CBS silencing resulted in decreased expression of Oct4, Sox2, and Nanog, with reduced structural integrity of spheroids. These findings suggest positive association between CBS and stemness transcriptional programs in OvCa spheroids.

Further protein-protein interaction (PPI) network analysis revealed that the stemness-associated proteins are significantly linked to EMT regulators that are crucial for metastatic onset (Supplementary File S1; Fig. S14). The expression of EMT transcription factors, such as Zeb1 and Twist1, was consistently reduced following CBS knockdown (Fig. 4M, N, O) in all three cell lines (Supplementary File S1; Fig. S15). Snail was found to be specifically downregulated in A2780-CP20 and OVCAR8 but not in COV318 (Supplementary File S1; Fig. S15), while Slug was repressed following CBS knockdown in COV318 and OVCAR8 but not across all biological replicates in A2780-CP20 (Supplementary File S1; Fig. S15).

Since the mesenchymal phenotype of cancer cells is influenced by factors that result in reduced E-cadherin levels and elevated N-cadherin and vimentin expression [30–32], we next evaluated this signature in spheroids grown under control and CBS-silenced conditions. Our data show that CBS knockdown induced a mesenchymal-to-epithelial transition marked by the up-regulation of E-cadherin and suppression of N-cadherin/ vimentin through repression of EMT transcription factors, viz. Zeb1, Snail, Slug and Twist1 (Fig. 4M, 4N, 4O). Furthermore, immunofluorescence analysis of the spheroids revealed that N-cadherin is localized at spheroidal cell junctions, with a reduction observed following CBS knockdown (Fig. 4P, Supplementary File S1; Fig. S19), indicating suppression of the mesenchymal phenotype. These results suggest that CBS promotes stemness circuitry and EMT programs in OvCa spheroids.

### CBS facilitates spheroidal architecture via Integrin β1 cell surface interactions and **downstream effector pathway**

Since spheroid histoarchitecture plays a key role in the survival and invasion potential of metastatic spheroids [8, 33], we evaluated the histoarchitecture of spheroids grown under control and CBS-silenced conditions. Cells with CBS knockdown did not form cohesive spheroids compared to the paired controls, which established spheroids of well-formed compact structure (Fig. 4Q, 4R and 4S). Specifically, CBS-deficient spheroids formed loose, poorly cohesive multicellular clusters with only a few cell-cell connections. These results indicate reduced spheroidogenesis and compactness in spheroids formed under CBS-silenced conditions. Thus, we next investigated factors that can influence spheroid architecture. Comparing the proteomic profiles of COV318 monolayers (2D) and spheroids (3D) that exhibited distinct signatures (PCA; Supplementary File S1; Fig. S20) provided us with multiple DEGs, the top 100 of which are depicted in Fig. 5A. The primary Log2 fold changes were visualized in a volcano plot to further identify the most significantly upregulated and downregulated DEGs (upregulated=67, downregulated =63) (Fig. 5B). Overrepresentation analysis with the Cancer Hallmark gene set revealed enrichment of pathways including resistance to cell death, tissue invasion and metastasis, and sustained angiogenesis which are critically involved in cancer progression and metastasis (Fig.5C). This was supported by the Enrichment analysis of upregulated proteins using MSigDB_Hallmark_2020 module which showed EMT to be the most enriched cancer hallmark upon anchorage independent (3D) growth of anoikis resistant cells (Supplementary File S1; Fig. S21E & S21F). Overrepresentation analysis and network analysis performed using the proteins most significantly upregulated in 3D compared to 2D culture identified β1 integrin (ITGB1) cell surface interactions (Fig. 5D, Supplementary File S1; Fig. S21A & S21B) and ITGB1-mediated cell signaling to be the top-enriched pathway, indicating a role for this pathway in OvCa spheroidogenesis and anchorage-independent survival. Interestingly, ITGB1 was also found to be the topmost interacting protein with the upregulated genes in the 3D spheroids using the PPI hub protein module in Enrichr (Supplementary File S1; Fig. S21C & S21D). Additionally, using the TNM plot functional analysis tool, Gene Ontology (GO) enrichment analysis of DEGs upregulated in 3D (as opposed to 2D) showed a significant enrichment for the cellular component term “Collagen-containing extracellular matrix” (GO:0062023) (Fig. 5F). This suggests that spheroids may actively remodel or interact with collagen-rich extracellular matrices. Additionally, it has been reported that ITGB1 mediates cancer cell anchoring, invasion, and metastasis by its binding to collagens [34]. Importantly, correlation analysis between CBS and ITGB1 in OvCa using the GEPIA tool showed their expression to be positively correlated (R=0.55, P-value=0) (Fig. 5G). To further validate our results from proteomic analysis, we next investigated the status of ITGB1 as a function of CBS knockdown in OvCa cell lines. CBS knockdown in spheroids resulted in downregulation of ITGB1 expression and repression of the downstream effector pathway, i.e., FAK signaling (Supplementary File S1; Fig. S22). In COV318 (Fig. 5H) and A2780-CP20 (Fig. 5I) spheroids, the inhibitory effect was evident with suppression of ITGB1-dependent focal adhesion kinase (FAK) signaling, while in OVCAR8, it was the level of phosphorylated active FAK that was reduced (Fig. 5J, Supplementary File S1; Fig. S22). The inhibitory effect of CBS knockdown on ITGB1 was found to be at the transcriptional level, as we observed *ITGB1* mRNA expression to be reduced at a relative fold change level of 0.72±0.75 in COV318, 0.63±0.03 in A2780-CP20 and 0.76±0.03 in OVCAR8 (Fig. 5K, 5L & 5M). Further, immunofluorescent analysis of ITGB1 in OvCa spheroids revealed its localization at spheroidal cell junctions, indicating its criticality in forming compact spheroids. However, spheroids with downregulated CBS had reduced ITGB1 expression, corroborating the observed loose architecture of the spheroids (Fig. 5N, 5O & 5P, Supplementary File S1; Fig. S23). These findings indicate that CBS plays a role in promoting spheroidal architecture via regulation of ITGB1 and associated downstream pathways.

**Fig. 5.**
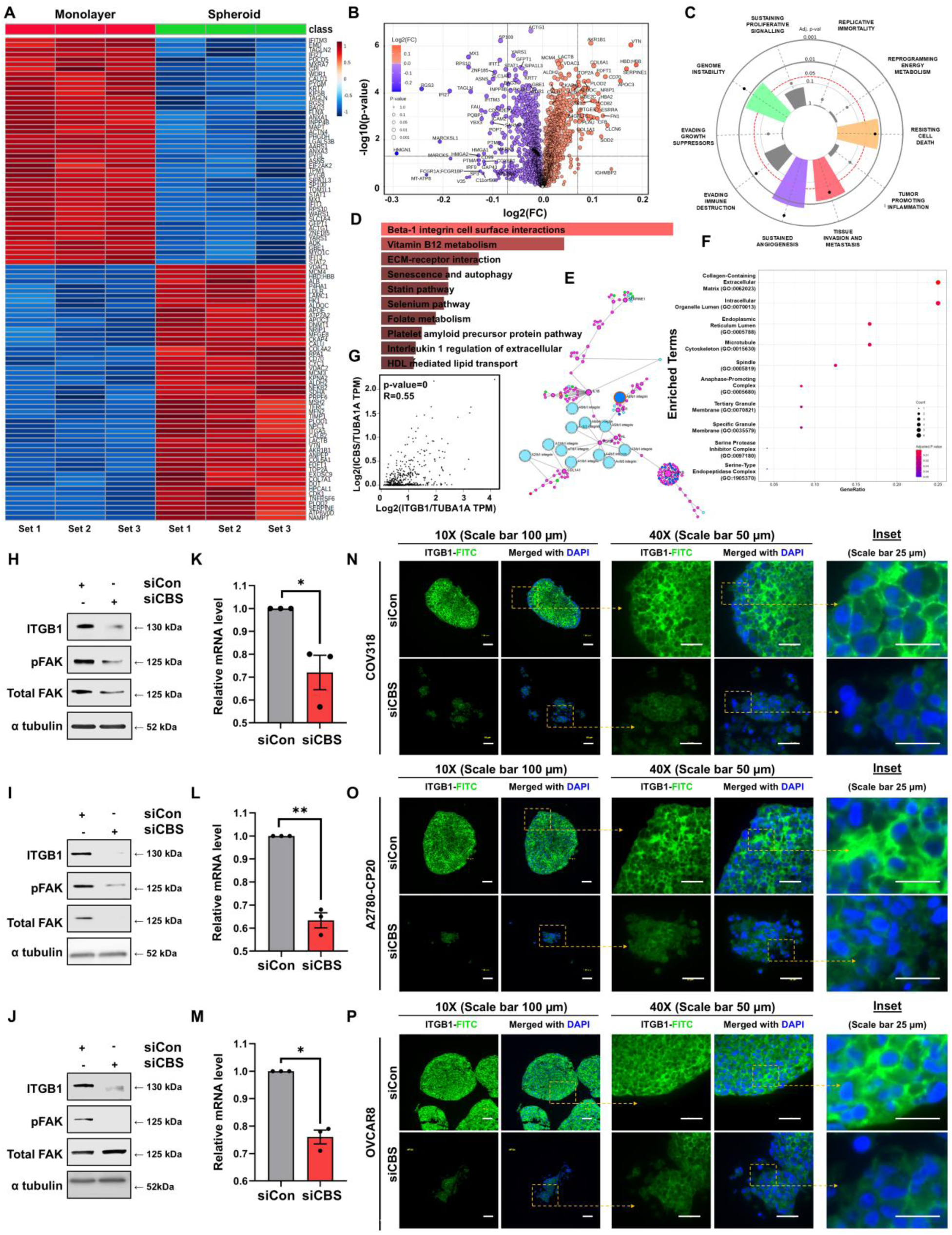
CBS promotes ITGB1-mediated signaling in OvCa spheroidogenesis. (A) Heat map of differentially expressed proteins (DEPs) between monolayer (2D) and spheroid (3D) cultures, representing top 100 differentially expressed genes (DEGs) upon proteomic analysis. (B) Volcano plot showing significantly upregulated and downregulated DEGs (log2FC=1.05, p=0.05). (C). ORA with Cancer Hallmark gene set using most significantly upregulated DEGs. (D &E) ORA using most significant upregulated genes from Enrichr database in 3D compared to 2D culture and network using ExpressAnalyst. (F) TNM plot enrichment, GO process analysis of ‘upregulated in 3D’ DEGs. Proteomic analysis was performed using three independent biological replicates (n=3) per condition (siCon and siCBS) (G) CBS-ITGB1 correlation: GEPIA analysis revealed a positive correlation between CBS and ITGB1 expression in OvCa (R = 0.55, p = 0). (H, I & J) Immunoblotting of COV318, A2780-CP20, and OVCAR8 spheroids showed that CBS depletion downregulates ITGB1 in all lines and reduced total FAK (COV318, A2780-CP20) or phosphorylated FAK (OVCAR8). (K, L & M) qPCR analysis of ITGB1 demonstrates significant transcriptional downregulation of *ITGB1* mRNA for COV318 (P value=0.02), A2780-CP20 (P value=0.007) and OVCAR8 (P value=0.01) following CBS silencing. (N, O & P) Immunofluorescence revealed ITGB1 accumulation at cell-cell junctions in CBS-expressing spheroids, whereas CBS knockdown caused reduced ITGB1, thereby reducing the spheroid compactness. All experiments were performed in three independent biological replicates (n = 3), and representative images are shown.

### CBS knockdown attenuates intraperitoneal metastasis of OvCa

Since our findings till now collectively suggest a role for CBS in promoting anchorage-independent survival of OvCa cells, i.e., a necessary event for transcoelomic metastasis, we sought to validate this in a murine model. We observed a significant reduction in omental homing and attachment of CBS-deficient spheroids (Fig. 6E & 6F, Supplementary File S1; Fig. S25). Additionally, since CD24+ve floating OvCa cells are more competent in terms of intraperitoneal dissemination [35], and there exists a significant positive association between CBS and CD24 mRNA expression levels (Spearman: 0.31 (p = 2.31e-12) in the OvCa patient cohort (Fig. 6B), we evaluated the CD24 composition of spheroids used in the *in vivo* assay. The flow cytometric analysis of spheroid-derived single cell suspensions revealed reduced CD24^+^/CD24^-^ cell ratio in the CBS-silenced group compared to the control (Fig. 6C& 6D) (Supplementary File S1; Fig. S24). These results suggest that CBS regulates the intraperitoneal dissemination of OvCa cells by modulating CD24 levels.

**Fig. 6.**
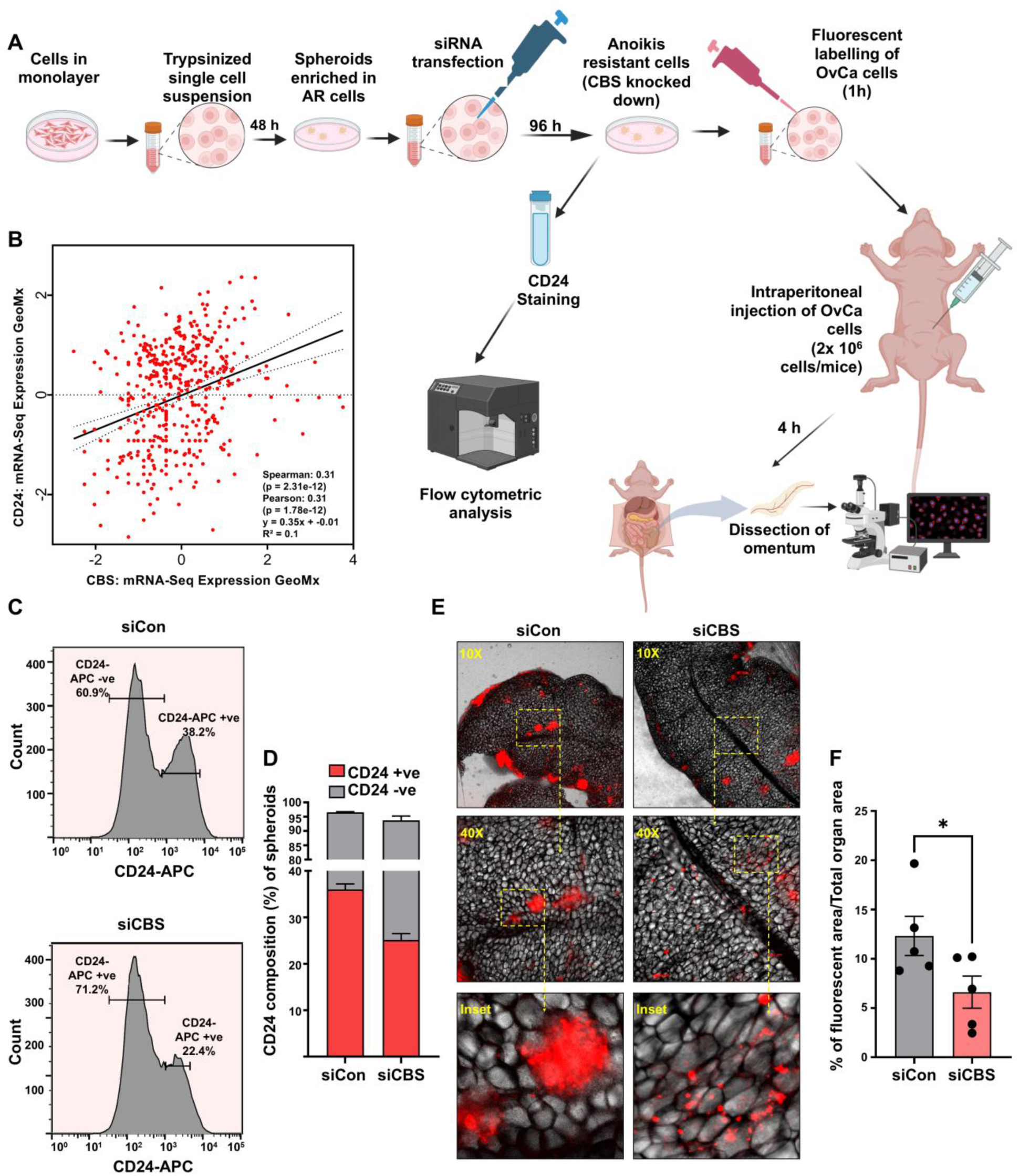
CBS promotes omental homing of OvCa cells in mice. (A) Workflow illustrating preclinical assessment of transcoelomic metastasis by CBS-expressing and CBS-silenced OvCa spheroids. (B) Correlation between CBS and CD24 mRNA expression in OvCa patients (Gray Foundation, Cancer Discovery 2024 cohort; data obtained from cBioPortal) (C) Representative histogram obtained through flow cytometry analysis showing CBS-knockdown mediated reduction in CD24+ve sub-population in anoikis-resistant spheroids, along with (D) bar diagram demonstrating statistical significance in depletion of CD24 composition of CBS-silenced spheroids (n=3; P value = 0.0073). (E) Representative photomicrographs of mouse omentum displaying adhered fluorescently labeled anoikis-resistant spheroidal cells, with corresponding quantification. (F) Data represent mean ± SEM* indicates that the Means are significantly different (n=5; P value ≤ 0.05).

### CBS-H_2_S-SP1 axis is crucial in mediating the pro-metastatic effects of CBS

The observed downregulation of *ITGB1* at the transcriptional level following CBS knockdown (Fig. 5K, 5L & 5M) prompted us to investigate potential transcription factors that may mediate the phenotypes associated with CBS knockdown. We aimed to identify a transcription factor that could serve as a molecular link between CBS knockdown and the downregulation of key proteins observed in our study. Transcription factor enrichment analysis using the top 30 ‘upregulated in 3D in comparison to 2D DEGs’ using TRRUST transcription factor 2019 and ChEA 2022 datasets in the Enrichr-KG web-server application identified SP1 as the most important pro-spheroidal transcription factor [36] (Fig. 7A).

**Fig. 7.**
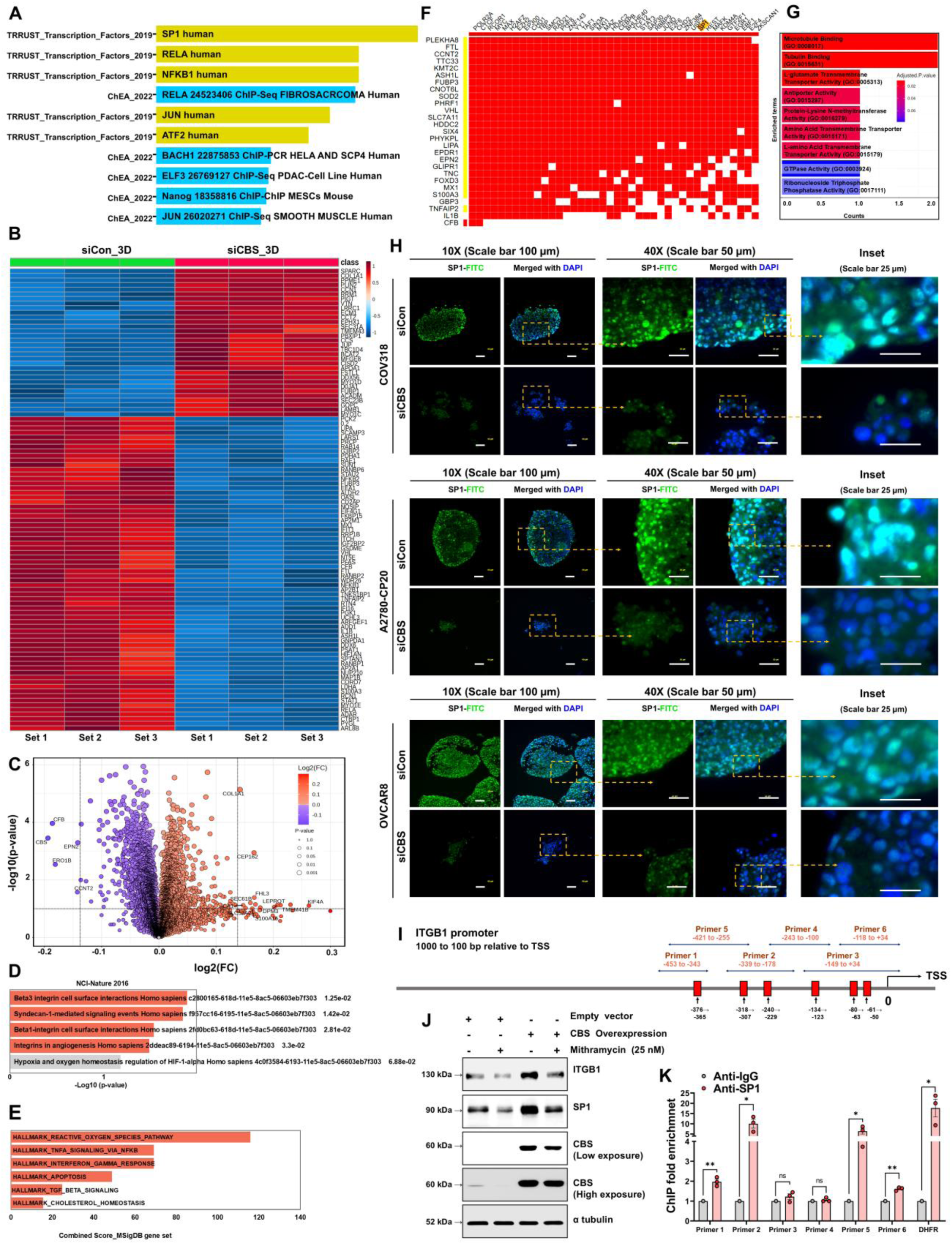
CBS knockdown-mediated repression of ITGB1 transcription and induction of apoptosis is associated with SP1. (A) Transcription factor enrichment analysis of 3D-upregulated genes (TRRUST 2019; ChEA 2022) identifies SP1 as the dominant regulator of pro-spheroidal gene expression. (B) Heat map representing top 100 DEGs upon proteomic analysis of 3D siCon vs 3D siCBS. (C) Volcano plot showing significantly upregulated and downregulated DEGs (log2FC=1.05, p=0.05). Pathway Enrichment analysis with the significant DEGs against NCI Nature 2016 database demonstrating Beta 1 integrin cell surface interaction among significantly enriched terms (D) and against MSigDB Hallmark gene set enrichment ranked by combined score, with apoptosis as the top hallmark (E). (F) Clustergram from Harmonizome showing the top 40 transcription factors, including SP1, predicted to regulate CBS silencing, associated-downregulated proteins. Proteomic analysis was conducted using n=3 independent biological replicates for siCon and siCBS each. (G) Gene Ontology (GO) enrichment analysis (Biological Process/Molecular Function) of CBS-dependent-SP1-regulated genes using OvCa datasets, revealing significant enrichment in microtubule binding, tubulin binding, amino acid transport, antiporter activity, and protein methyltransferase activity, indicating roles in cytoskeletal organization, metabolic transport, and epigenetic regulation. (H) Immunofluorescence showing reduced nuclear SP1 in CBS-silenced spheroids of COV318, A2780-CP20 and OVCAR8 compared to controls. (I) Schematic diagram representing putative SP1 binding sites on ITGB1 promoter (−1000 to +100 bp), where Primers 1, 2, 3, 4, 5 and 6 denote the primer sets used for ChIP-qPCR. (J) CBS overexpression causes ITGB1 upregulation, which is again reduced by mithramycin (25 nM), indicating involvement of SP1 in the regulation of CBS-mediated ITGB1 expression. (K) ChIP-qPCR analysis showing enrichment of SP1 binding at the ITGB1 promoter. Chromatin immunoprecipitated with anti-SP1 antibody shows increased enrichment across the Primer 1, Primer 2, and Primer 5 compared to IgG control, confirming SP1 occupancy at predicted binding sites. DHFR is included as a positive control for SP1 binding. Data are presented as mean ± SEM.

Next, we performed proteomic analysis comparing CBS-expressing to CBS-silenced spheroids, revealing multiple DEGs, among which the top 100 are shown in Fig. 7B. The PCA indicated that the siCon 3D and siCBS 3D samples had distinct proteomic profiles (Supplementary File S1; Fig. S26). The volcano plot (Fig. 7C) depicts the primary Log2 fold changes and led us to identify 38 significantly downregulated and 44 significantly upregulated DEGs. Pathway Enrichment analysis with these DEGs against the NCI Nature 2016 [37] database showed ITGB1 cell surface interaction as one of the most enriched terms (Fig. 7D). Additionally, the same approach with the MSigDB Hallmark gene set revealed apoptosis as a significantly enriched term (Fig. 7E).

Transcription factor enrichment analysis of the top 30 downregulated DEGs in CBS-silenced spheroids against Harmonizome ENCODE Transcription Factor Targets dataset [38] resulted in a clustergram providing multiple candidate transcription factors, among which SP1 ranked at position 32 among the top predicted transcription factors (Fig. 7F). Removing the basal transcription factors, we were left with CTCF, MYC, MAX, ZNF143, YY1, MXI1, MAZ, CEBPB, BHLHE40, TCF12, STAT3, JUND, E2F6, ZNF384, USF2, SP1, REST, MAFK, EGR1, EBF1, E2F4 and ZKSCAN1 (list presented according to rank). This analysis provides high relevance to the study as it identifies these transcription factors behind spheroidogenesis, some of which, such as Myc [39] and STAT3 [40], are well known to be highly expressed in OvCa and support disease progression. Given that this analysis identified SP1 as the only transcription factor listed among the topmost enriched ones, behind the pro-spheroidogenic DEGs found in our study (Fig. 7A), and since SP1 has been previously reported to mediate CBS-driven functions in OvCa [22], we hypothesised SP1 to be a downstream target of CBS mediating its pro-spheroidogenic functions. We found that among the top CBS-dependent genes, 75% of DEGs were under transcriptional control of SP1 (Supplementary File S1; Fig. S28A). These genes (list presented in Supplementary File S1; Fig. S28B), when subjected to functional enrichment using GO: biological process against OvCa dataset, showed enrichment in microtubule binding, tubulin binding, amino acid transport, antiporter activity, and protein methyltransferase activity, suggesting roles in cytoskeletal organization, metabolic transport, and epigenetic regulation (Fig. 7G, Supplementary File S1; Fig. S28C). Further pathway analysis using Enrichr (Supplementary File S1; Fig. S28D, S28E, S28F) showed enrichment of pathways associated with transcriptional regulation, cellular stress response, cell proliferation, and apoptotic signaling. Especially, pathways associated with oncogenic signaling, ferroptosis, and extracellular matrix regulation were significantly enriched, in line with a role for CBS-SP1 signaling in metabolic adaptation and tumor progression.

Additionally, our in-silico analysis using the JASPAR database revealed six high-confidence SP1 binding motifs (relative score ≥0.90) enriched within the proximal ITGB1 promoter (Site 1: −50 to −61, Site 2: −63 to −80, Site 3: −123 to −134, Site 4: −229 to −240, Site 5: −307 to −318, Site 6: − 365 to −376 relative to TSS) (Fig 7I). To experimentally confirm if SP1 is downstream of CBS and if it regulates ITGB1, we ectopically overexpressed CBS in OVCAR8 cells and measured downstream protein levels with or without the SP1-inhibitor mithramycin [41] to ascertain whether CBS controls ITGB1 expression via SP1. When compared to empty vector controls, immunoblot analysis demonstrated that CBS overexpression resulted in a significant increase in ITGB1 protein levels (Fig. 7J, Supplementary File S1; Fig. S30). Interestingly, mithramycin treatment (25 nM) substantially reduced this induction, implying that transcriptional activity of SP1 is required for CBS-mediated ITGB1 upregulation. Moreover, CBS overexpression also induced SP1 protein expression, indicating that CBS positively regulates SP1 stability and/or abundance. These results collectively show that CBS causes ITGB1 upregulation via a SP1-dependent mechanism, suggesting mechanistic support for a CBS-mediated SP1-dependent ITGB1 regulation that induces the metastatic potential of OvCa cells.

Next, we proceeded to experimentally validate whether SP1 directly binds the ITGB1 promoter. Since we found six high-confidence SP1 binding motifs through our *in-silico* analysis, we used ChIP-qPCR with six regions (covered by Primer 1, 2, 3, 4, 5 and 6) harboring one or more SP1 binding motifs (Fig 7I). The results revealed region-specific SP1 occupancy. Compared with IgG control, SP1 enrichment was found to be significant at Primer 1, Primer 2, and Primer 5 regions. The strongest enrichment was observed at Primer 2 (−318 to −178 bp) and Primer 5 (−421 to −255 bp), with a lower but significant signal at Primer 1 (−453 to −343 bp). These results suggest that binding sites “-376 to −365”, “-318 to −307” and “-240 to −229” at the ITGB1 promoter are crucial for SP1 binding in OvCa cells (Fig 7K). The DHFR promoter, used as a positive control, showed robust SP1 enrichment, validating assay specificity. These findings demonstrate that SP1 directly associates with multiple regions of the ITGB1 promoter.

Here, it should be mentioned that while CBS-generated H_2_S has earlier been reported to promote SP1 stability via its S-sulfhydration in endothelial cells [17], to our knowledge, this represents the first demonstration of CBS-dependent SP1 regulation in OvCa. In line with this, our investigation regarding potential persulfidation sites of SP1 using the amino acid sequences in pCysMod, a deep learning-based platform for predicting multiple cysteine modifications, indicated the same. We found that the three isoforms of SP1 had identifiable persulfidation sites; for NP_003100.1: Cys621, Cys626 (Supplementary File S1; Fig. S27A), for NP_001238754.1: Cys 580, Cys 585 (Supplementary File S1; Fig. S27B) and for NP_612482.2: Cys 628, Cys 633) (Supplementary File S1; Fig. S27C). In our study, reduced overall expression of SP1, leading to its attenuated nuclear presence in CBS-silenced spheroids (Fig. 7H, Supplementary File S1; Fig. S29) and its amelioration in CBS-silenced spheroids following H_2_S supplementation (discussed in the next segment), provides the first report of CBS-mediated regulation of SP1 in OvCa that ultimately drives the pro-spheroidal and anoikis-evasive effects. To directly determine whether SP1 undergoes persulfidation in a CBS-dependent manner, we conducted modified biotin-switch assays in OVCAR8 cells with CBS-expressing, CBS-silenced and GYY4137-supplemented

CBS-silenced cells. SP1 was consistently detected in the biotin-switch fraction under control conditions, indicating the presence of basal SP1 persulfidation (Fig. 8A, 8B, Supplementary File S1; Fig. S31). Notably, despite comparable SP1 levels in the input samples, CBS silencing (siCBS) resulted in a substantial reduction in SP1 persulfidation. Treatment with GYY4137 restored SP1 persulfidation in CBS-silenced cells, supporting the role of CBS-derived H_2_S in facilitating this modification. GAPDH, a recognized persulfidation target of CBS, showed expected patterns across conditions and therefore served as a positive control, suggesting assay specificity. Therefore, these data provide direct biochemical evidence that SP1 undergoes persulfidation in a CBS-dependent and H_2_S-mediated manner.

**Fig. 8.**
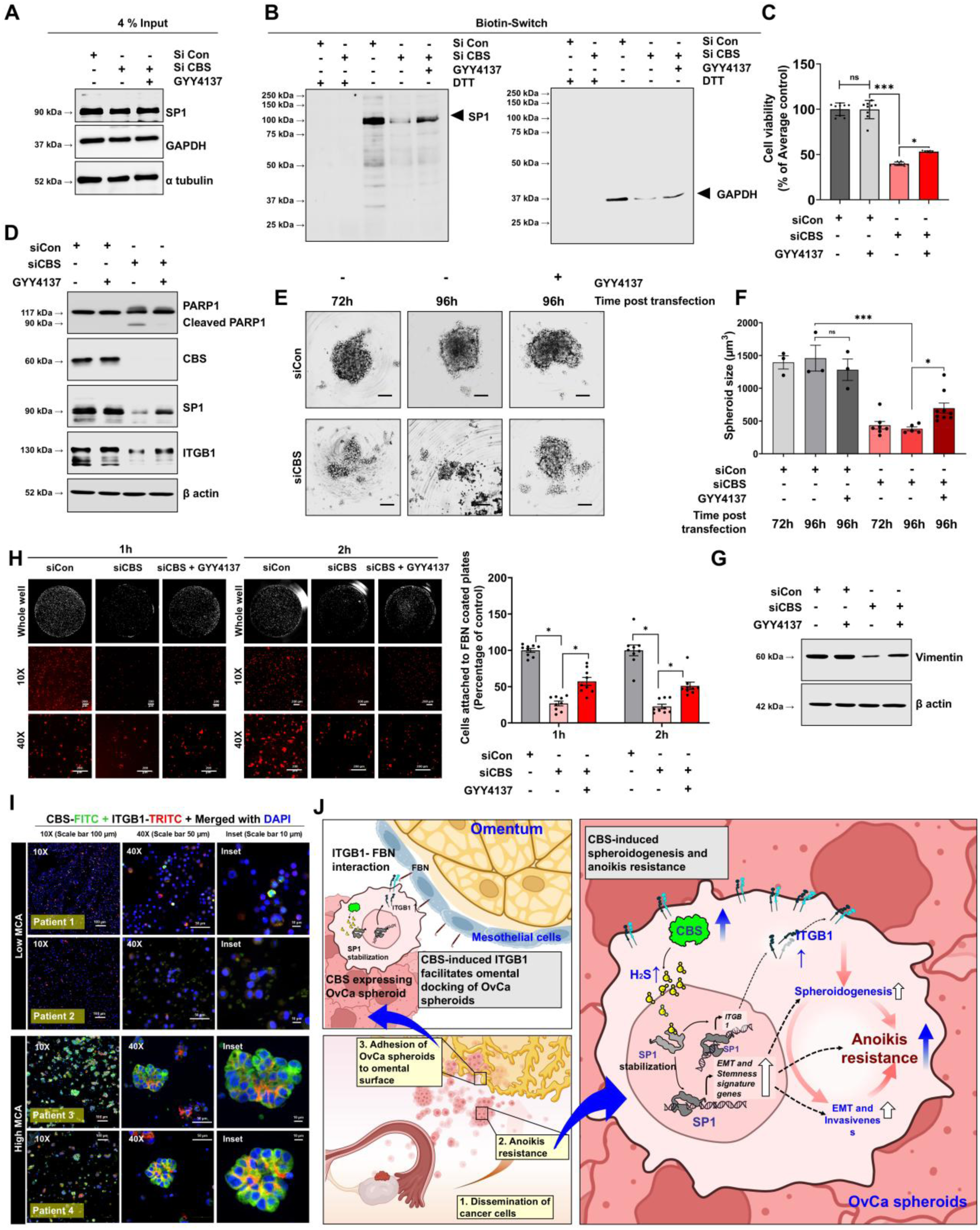
CBS-derived H_2_S persulfidates and stabilizes SP1 to promote SP1-mediated ITGB1 regulation and spheroidal function in OvCa. (A) Immunoblots showing SP1 persulfidation assessed using a modified biotin-switch assay with siCon or siCBS-transfected OVCAR8 cells. For rescue, cells were treated with 1 mM GYY4137 for the final 24 h, i.e., 72 h after knockdown. Dithiothreitol (DTT) treatment was used as a negative control to reduce persulfide bonds. Left panels (4% input): Total protein lysates showing SP1 expression levels across conditions and indicating comparable protein loading. GAPDH and α-tubulin serve as loading controls. (B) Enrichment of persulfidated SP1 following biotin labeling and pull-down. GAPDH is shown as a known persulfidation target of CBS, serving as a positive experimental control. (C) Rescued anchorage-independent survival upon GYY4137 treatment of siCBS COV318 spheroids. (D) Immunoblot of PARP1, CBS, SP1 and ITGB1 indicating restoration of spheroidal viability and function in CBS-silenced COV318 spheroids following H_2_S supplementation. (E) Microscopy images showing more compact spheroids in GYY4137-treated CBS-knockdown cells. (Scale bar = 200 µm). (F) Bar diagram representing difference in spheroid sizes of each group. While siCBS exhibited significantly reduced size after 96h (P value=0.0004), GYY4137 treatment ameliorated this (P=0.01). (G) Immunoblot panel demonstrating restoration of vimentin as a representative of OvCa spheroidal invasiveness, in CBS-silenced spheroids following GYY4137 treatment. (H) Photomicrographs of fibronectin-coated wells (96 well plate) plated with labelled CBS-expressing and CBS-silenced spheroidal cells after 1 and 2h of transfer onto FBN-coated wells, with bar diagram indicating the number of cells attached to an FBN-coated well of 96-well plate in siCon, siCBS, and siCBS supplemented with GYY4137 treatment. All experiments were conducted in three independent biological replicates (n=3). (I) Representative immunofluorescence images of patient-derived ascites samples classified based on low and high multicellular aggregate (MCA) content. The upper panel shows low MCA ascites samples (Patients 1 and 2), characterized by non-spheroid single tumor cells with low CBS and ITGB1 signal. The lower panel shows high MCA ascites samples (Patients 3 and 4), exhibiting compact multicellular aggregates (spheroids) with marked enrichment of CBS (Green) and ITGB1 expression. The scale bars are 100 µm (10×), 50 µm (40×), and 10 µm (insets). (J) Schematic diagram depicting the mechanism by which CBS drives OvCa transcoelomic metastasis, where cancer cells disseminate, resist anoikis through spheroidogenesis, and adhere to secondary sites. CBS promotes H_2_S-mediated SP1 stabilization and hence spheroid stemness, mesenchymal traits, along with upregulating ITGB1 to maintain spheroid compactness and omental adhesion.

### H_2_S donor restores the SP1-dependent ITGB1 regulation and spheroidal function in CBS-deficient OvCa spheroids

Since SP1 persulfidation is mediated by H_2_S produced by CBS via the transsulfuration pathway, we next investigated whether H_2_S supplementation could rescue the loss of spheroidal compactness and size observed in CBS-silenced COV318 cells. Our data indicate reduced anchorage-independent survival in CBS-deficient 3D cultures, which was significantly restored upon treatment with GYY4137 (1 mM), a slow-releasing H_2_S donor. Restoration of spheroidal growth (Fig. 8E, Supplementary File S1; Fig. S32), along with immunoblotting data showing decreased levels of cleaved PARP1 (Fig. 8D, Supplementary File S1; Fig. S33), strongly suggests that CBS exerts its pro-AR and pro-spheroidal effects through H_2_S signaling. Further evidence showing the restoration of SP1 and ITGB1 protein levels previously downregulated by CBS knockdown confirmed the role of the CBS-H_2_S-SP1 axis in regulating ITGB1 and driving OvCa spheroidogenesis (Fig. 8D, Supplementary File S1; Fig. S34).

Given that ITGB1 is a receptor for fibronectin (FBN), and previous studies [42] have shown that the interaction between ITGA5/ITGB1 on OvCa spheroids and FBN expressed by omental mesothelial cells is critical for driving omental metastasis [43], we next performed FBN adhesion assays using single-cell suspensions derived from spheroids. CBS-knockdown cells showed significantly reduced adhesion to FBN (percentage of control: 26.82 ± 4.91 at 1 h and 21.26 ± 8.88 at 2 h), which was partially restored upon treatment with GYY4137 (percentage of control: 57.19 ± 6.39 at 1 h and 46.15 ± 3.55 at 2 h) (Fig. 8H), further supporting a critical role for ITGB1 in omental metastasis (Supplementary File S1; Fig. S35). Additionally, H_2_S-mediated restoration of Vimentin indicated renewal of spheroid invasiveness in CBS-silenced cells (Fig. 8G, Supplementary File S1; Fig. S33). These results suggest that CBS is upstream to SP1-mediated regulation of ITGB1 and spheroidal function in OvCa spheroids through H_2_S signaling.

### CBS expression is enriched in multicellular aggregates and associates with ITGB1 in patient-derived ascites

To assess the clinical relevance of our mechanistic findings, we examined patient-derived OvCa ascites samples classified by MCA content. Immunofluorescence staining for CBS, ITGB1, and DAPI demonstrated distinct expression patterns in low- and high-MCA samples (Fig. 8I, Supplementary File S1; Fig. S36-S39). Cells in ascites samples with low MCA content (patients 1 and 2) primarily demonstrated non-spheroid single cell organization exhibiting low CBS and ITGB1 expression. In contrast, high MCA samples (patients 3 and 4) had well-defined spheroidal aggregates exhibiting high CBS and ITGB1 expression. These findings collectively indicate that CBS-ITGB1 regulation is associated with spheroid-rich ascites, suggesting its involvement in AR.

## Discussion

OvCa metastasis differs from most solid tumors as the predominant route is transcoelomic rather than hematogenous dissemination [44]. During this process, cancer cells detach from the ovarian surface epithelium, survive in suspension in the peritoneal fluid, and subsequently adhere and invade secondary sites. To survive this anchorage-independent phase, these cells must evade anoikis, a form of programmed cell death triggered by detachment. A key mechanism facilitating AR is the formation of MCAs or spheroids. Studies have demonstrated that OvCa cells that do not form spheroids are more vulnerable to anoikis and are therefore less metastatic [8]. Moreover, spheroids with higher structural integrity show increased drug resistance and AR and are therefore more aggressive in driving intraperitoneal metastasis [33, 44]. Together, these observations identify spheroidogenesis as a key event in OvCa transcoelomic metastasis.

Recent studies have suggested the role of metabolic enzymes as regulators of spheroid survival [45, 46]. CBS is a key transsulfuration pathway enzyme that catalyses the conversion of methionine-derived homocysteine into cysteine [47]. This pathway supports the biosynthesis of GSH and generates H_2_S, a gas transmitter which modulates protein function via persulfidation [48]. Previous studies have confirmed a role for CBS in promoting OvCa progression, redox adaptation, and chemoresistance [19, 21–23, 49], but its function in OvCa metastasis remains unexplored, especially in the context of spheroidogenesis and AR. Our in-house TMA analysis demonstrated that CBS overexpression correlates significantly with poor patient survival and increased prevalence of omental metastasis, highlighting the clinical relevance of CBS in OvCa metastasis. Interestingly, our comparative analysis of 2D monolayer and 3D spheroid cultures revealed that CBS is upregulated in COV318 spheroids, implying its role in OvCa anchorage-independent survival.

Given the established role of spheroidogenesis and cancer stemness in transcoelomic dissemination of OvCa, we next sought to examine the relationship between CBS expression and stemness-associated factors. We observed positive correlation between CBS expression and core stemness regulators Nanog, Oct4, and Sox2, suggesting that CBS actively contributes to regulating the stem-like state of OvCa cells. Moreover, immunofluorescence-based co-localization analyses revealed that CBS-expressing cells in spheroids spatially coincide with Oct4, Sox2, and Nanog-positive populations, thereby underpinning the association between CBS expression and OvCa stemness. Corroborating this, transcription factor analysis of CBS-influenced DEGs, downregulated in CBS-silenced COV318 monolayer cells (as identified by proteomic profiling), revealed enrichment of Sox2, Oct4, and Nanog among the top transcriptional regulators, mediating CBS-induced oncogenic stemness. Further, over-representation analysis (ORA, using integrated cancer hallmark gene set (n=6763) (https://cancerhallmarks.com/) revealed that CBS-regulated DEGs are enriched in cancer hallmark gene sets associated with resistance to cell death. Corroborating this, our *in vitro* studies suggested that CBS silencing in 2D monolayer resulted in cell death. In control cells, F-actin was organized either as central clusters of actin aggregates or as a dispersed meshwork of short filaments throughout the cytoplasm. In contrast, CBS-silenced cells exhibited prominent thick bundles of actin filaments, stress fibers, located near the plasma membrane, cytoskeletal rearrangements, which were previously associated with apoptosis in embryonal carcinoma cells [50]. Additional apoptotic features, such as membrane blebbing and cell shrinkage, were also observed in CBS-knockdown cells. The induction of cleaved PARP1, corroborated by these morphological observations, confirmed the occurrence of apoptosis following CBS silencing. Together, these results confirm CBS as a key regulator of OvCa cell survival.

To further examine the role of CBS in transcoelomic dissemination, we utilized spheroids (3D cultures), which closely mimic MCAs, the functional units of OvCa metastasis in the peritoneal cavity. CBS silencing caused significant reduction in spheroid viability and caused time-dependent inhibition of spheroid growth, suggesting its key role in anchorage-independent survival and proliferation in OvCa. These results are very pertinent as spheroids in OvCa are metastatic units responsible for transcoelomic dissemination, with more spheroidogenesis associated with anoikis evasion and peritoneal docking [8]. This loss of viability was mechanistically linked to anoikis, as evident by increased cleaved PARP1, Bax induction, and Bcl-2 suppression [51]. This underscores that CBS is an upstream regulator of the cascade that enables OvCa spheroids to escape anoikis through activation of cell survival pathways via upregulation of anti-apoptotic proteins, including Bcl-2 [10, 52].

Since cancer stemness plays a critical role in MCA/spheroid formation in OvCa [29] and AR [53] [54], we speculated that the observed CBS-depletion-associated anoikis in our study might be associated with the disruption of stemness-related transcription factors. Our data showed that CBS knockdown resulted in coordinated inhibition of the pluripotency network, including Oct4, Sox2, KLF4, and Nanog, as well as the CSC marker CD44, and key EMT markers (Zeb1, Snail, Slug, Twist), indicating a loss of stem-like and mesenchymal attributes. Notably, the mesenchymal signature, marked by increased N-cadherin and vimentin along with decreased E-cadherin, is known to support AR [55] and has been associated with Bcl-2 upregulation in prostate [56] and breast cancers [57]. Moreover, N-cadherin⁺ OvCa spheroids are reported to be more cohesive, adhesive to mesothelium, and metastatic than E-cadherin⁺ or hybrid phenotypes [58]. Additionally, low E-cadherin and high ITGB1 levels are linked to compact, invasive spheroids, while vimentin induction also marks increased invasiveness [59]. Similarly, CBS-silenced spheroids in our study showed reduced N-cadherin and vimentin, increased E-cadherin, and disrupted histoarchitecture, indicating attenuated metastatic potential.

Like cadherins, integrins are heterodimeric transmembrane proteins, comprising α and β chains, that facilitate spheroidal cell-adhesion through their interaction with connective tissue components like FBN or ECM glycoproteins, including laminins and collagens. Among integrins, α5β1, αvβ3 and α2β1 have been found to play a key role in the poor prognosis of OvCa [60]. Integrin α5β1 has been confirmed to significantly contribute to MCA-associated peritoneal metastasis [61]. Moreover, ITGB1 overexpression is associated with OvCa spheroid compactness [59], a phenotype associated with promotion of AR, ultimately inducing transcoelomic metastasis [33]. In order to uncover molecular drivers underlying spheroidogenesis, our comparison of the proteomic profiles of monolayer (2D) and anchorage-independent (3D spheroid) HGSOC ascites-derived cells (COV318) identified ITGB1-mediated signaling as central to OvCa anchorage-independent survival and spheroid formation. Additionally, this analysis also revealed key differentially expressed proteins between monolayer and spheroid cultures, highlighting distinct pathways involved in spheroid-mediated OvCa dissemination, many of which have not been previously explored in the context of spheroidal morphogenesis. Therefore, we examined the expression of ITGB1 and its downstream effector FAK [62], and found that CBS knockdown reduced ITGB1 expression in spheroids from all three examined cell lines, FAK expression in COV318 and A2780-CP20, and activated FAK levels in OVCAR8. Transcript-level analysis confirmed reduced *ITGB1* mRNA upon CBS depletion in OvCa spheroids. Immunofluorescence analysis revealed ITGB1 to be enriched at spheroid cell junctions in controls, similar to N-cadherin, supporting spheroid cohesion, whereas CBS-deficient spheroids showed ITGB1 loss and disrupted architecture, ultimately indicating probable attenuation of their metastatic potential. To test this *in vivo*, we conducted homing experiments in female Nu/J mice and observed reduced omental metastasis from CBS-depleted anoikis-resistant cells. Moreover, since CD24⁺ OvCa spheroidal cells are known to have higher metastatic potential [35], the depletion of the CD24⁺ subpopulation in these spheroids upon CBS-knockdown further suggested reduced metastatic potential from CBS silencing. It is important to note here that CD24 has been identified as a context-dependent marker for stem-like and tumor-initiating populations, differing from the CD44⁺CD24⁻ paradigm as robustly observed in breast cancer [63, 64]. These findings corroborate the study by Li et al., reporting that CD24 plays an important role in the development of OvCa AR [65].

The transcriptional downregulation of ITGB1 following CBS silencing suggested the involvement of a transcription factor linking CBS to the observed changes in proteomic profile. Transcription factor enrichment of DEGs upregulated in 3D identified SP1 as a key regulator of pro-spheroidal proteins. Interestingly, SP1 also emerged among the top predicted regulators of CBS-dependent DEGs downregulated in spheroids. Furthermore, functional enrichment analysis of genes that are both CBS-dependent and SP1-regulated in OvCa spheroids demonstrated significant enrichment of pathways related to survival, cell adhesion and metastatic progression, thus supporting its role in facilitating pro-spheroidal phenotypes. Therefore, SP1 could be considered a putative effector of spheroidogenesis downstream of CBS. Our ‘CBS gain-of-function coupled with pharmacological inhibition of SP1’s studies further supported this functional association. In these studies, CBS overexpression raised the levels of SP1 and ITGB1, while treatment with the SP1 inhibitor, mithramycin A [41], lowered ITGB1 expression even though CBS was overexpressed. This showed that SP1 is responsible for regulating ITGB1. In line with this, we found direct evidence of SP1 binding at the ITGB1 promoter via chromatin immunoprecipitation (ChIP)-qPCR analysis, confirming that ITGB1 is a direct transcriptional target of SP1 in OvCa cells. Although the detailed mechanism regarding this has not been reported in OvCa, CBS-derived H_2_S has been shown to stabilize SP1 through S-sulfhydration in endothelial cells [17]. Corroborating this, our computational prediction identified persulfidation-prone cysteine residues across all SP1 isoforms, supporting the probability of H_2_S-mediated regulation. Considering previous findings about SP1 mediating the CBS-driven pro-tumorigenic effects of OvCa [22], we hypothesized SP1 to bridge CBS to its pro-spheroidal and pro-metastatic programs evident in this study. Importantly, we offer direct biochemical evidence substantiating this mechanism, evidenced by biotin-switch assays indicating CBS-dependent persulfidation of SP1, which diminished upon CBS silencing and was restored following H_2_S supplementation. This situates SP1 as a downstream target of CBS-derived H_2_S in OvCa cells.

Interestingly, while ITGB1 downregulation was not detected in the proteomics experiment, pathway enrichment of CBS-dependent DEGs demonstrated ITGB1-mediated signaling, consistent with our immunoblotting, transcriptional, and immunofluorescence data. Similar discrepancies between proteomics and other detection methods for membrane proteins have been widely reported and are often attributed to limitations of mass spectrometry, especially with large, hydrophobic, and glycosylated proteins like ITGB1 [66–70]. Glycosylation and membrane association can hinder peptide ionization and digestion, resulting in underrepresentation in proteomic data. For instance, Liu et al. [71] found ITGB4 upregulation by Western blot following PRA1 silencing, despite its absence in proteomic profiles, which was attributed to poor solubilization and ionization of glycosylated, membrane-bound proteins [66, 67, 71]. Nevertheless, CBS-dependent DEGs pointed us toward SP1 as a key regulatory link. Confirming this, we found that CBS knockdown reduced the nuclear presence of SP1 in spheroidal cells while H_2_S supplementation restored its expression in these spheroids. Mechanistically, we demonstrate here that H_2_S stabilizes SP1 [17], which transcriptionally upregulates ITGB1 to promote spheroidal cell adhesion to FBN-coated surfaces that mimic the FBN-expressing microenvironment of mesothelial cells on the omental surface. These findings are highly relevant since ITGB1-FBN interactions drive OvCa omental metastasis, with cancer cells inducing mesothelial cell FBN release, which in turn binds OvCa cell integrins α5/β1 to promote adhesion[43]. At the phenotypic level, H_2_S supplementation in CBS-silenced spheroids supported anchorage-independent growth and mitigated anoikis. Reduced PARP1 cleavage, along with restoration of SP1 and ITGB1 expression, further confirmed the role of the CBS-H_2_S-SP1 axis in sustaining these pro-metastatic traits in OvCa. Additionally, vimentin, a known mediator of OvCa spheroid invasiveness during mesothelial clearance [55], was also downregulated following CBS knockdown and rescued after H_2_S supplementation, highlighting CBS as a key upstream regulator of transcoelomic metastasis.

A limitation of our study is the variability observed in the perturbation of some markers (e.g., Sox2, Snail, and Slug) across biological replicates following CBS silencing in 3D. This likely reflects the dynamic and context-dependent features of cancer stemness and EMT programs, affected by tumor heterogeneity and cellular plasticity [72, 73]. Consequently, these markers were analyzed qualitatively, and our conclusions are based on consistent trends observed across various cell lines and supplementary assays. These shortcomings, however, do not impede the overall conclusions, which are supported by integrating multiple complementary experimental approaches.

Notably, the analysis of patient-derived ascites that indicated high CBS in MCAs/spheroids was consistent with our TMA analysis that demonstrated similar findings on OvCa tissue of patients with omental metastasis. Additionally, high ITGB1 in the ascites of high MCA patients indicated the role of CBS-ITGB1 regulation in OvCa MCA formation and AR. While consistent expression patterns were observed across multiple microscopic fields within each sample, our study is limited by the small number of patient-derived ascites samples analyzed (N=4). Future studies with larger patient cohorts and organoid-based systems will be highly helpful to further confirm the clinical relevance of the CBS-SP1-ITGB1 axis in actuating OvCa AR. Nevertheless, the multilayered approaches used in this study situate CBS as a potential therapeutic target to mitigate OvCa metastasis. We identified key molecular differences between 2D and 3D cultures, offering novel insights into metastatic mechanisms. Mechanistically, the CBS-H_2_S-SP1 axis promotes transcription of stemness and EMT factors, along with ITGB1, supporting AR, omental homing and colonization during transcoelomic metastasis (Fig. 8J). These findings pave the way for future translational research into targeted strategies to mitigate OvCa metastasis, where we can utilize CBS inhibitors in patients with ascites that will inhibit metastasis and disease progression to give us more time to treat the patients with a possibility of better therapeutic outcomes.

## Acknowledgements

We acknowledge the MTCRO-COBRE Cell & Tissue Analysis Core and the OU Health Stephenson Cancer Center Tissue Pathology and Biospecimen Shared Resource for cell authentication and histology services, respectively.

## Authors contribution

G.R., S.K.D.D., and P.S. conceived and designed the study. P.S. performed data acquisition, conducted data analysis, and wrote the original manuscript draft. A.D.B. and A.P.J. contributed to data analysis, interpretation, and manuscript revision. S.C. contributed to acquisition and analysis of flow cytometry data. R.V. and S.R. generated and analyzed the tissue microarray data. C.X. performed statistical validation. D.D., Z.J.Z, P.M., and R.B. provided essential resources and critical scientific input. G.R. and S.K.D.D. accessed and verified the underlying data, supervised the overall study, and contributed to manuscript revision. All authors read and approved the final manuscript.

## Data availability

All data needed to evaluate the conclusions in the article are present in the article and/or the Supplementary Materials (including uncropped blots in Supplementary File S2). The data and materials used in the current study are available from the corresponding authors upon reasonable request.

## Declarations

### Consent for publication

Consent to Publish declaration: not applicable

### Consent to Participate

Consent to Participate declaration: not applicable.

### Ethics Declaration

Ethics declaration: All animal experiments were performed under IACUC-approved protocols (Approval # 25-007-CH). No additional ethical approvals were required.

Regarding the use of patient-derived ascites samples, de-identified ascites samples were collected from OvCa patients during paracentesis under an active IRB protocol (#2044) approved by the Institutional Review Board at the University of Oklahoma Health Campus.

