## Supplementary File 1 for "Cystathionine Beta-Synthase Promotes Anoikis Resistance and Transcoelomic Metastasis in Ovarian Cancer via SP1- ITGB1 regulation"

***Correspondence:**

**Supplementary file 1**

### Methods

#### 1. Bioinformatic analysis

- To evaluate patient vital status across FIGO stages, we integrated publicly available OvCa data sets from CPTAC, GDC, Gray Foundation, Cancer Discovery 2024 [1], MSK SPECTRUM (MSK, Nature 2022) [2], and TCGA, GDC to construct a combined dataset comprising 764 patients (1,364 tumor samples) (https://www.cbioportal.org/, last accessed on 07.08.2025) via cBioPortal [3]. Bar diagram was plotted in GraphPad Prism 10.8.1 using data on each patient's vital status vs FIGO stages, extracted from the constructed dataset. The data is presented in Fig. S1.

- To evaluate the expression of CBS mRNA in normal and OvCa tumor tissues, the dataset was extracted for differential gene expression analysis in tumor, node, and metastatic tissues (TNM plot; https://www.tnmplot.com, last accessed June 6, 2025) [4]. The data is presented in Fig. 1A.

- To assess the distribution of CBS mRNA expression across OvCa FIGO stages, data was extracted from the Ovarian Cancer (CPTAC GDC, 2025) dataset in cBioPortal and visualized as scatter plots showing individual data by GraphPad Prism 10.8.1. The data is presented in Fig. 1B.

- To evaluate the prognostic significance of CBS expression at different disease stages in OvCa patients, we employed KM-plotter tool (https://kmplot.com/) for survival analysis. The data is presented in Fig. 1C.

- In line with our finding that CBS promotes ITGB1 expression in OvCa, we performed pairwise gene correlation analysis for CBS vs. ITGB1 using the Pearson correlation statistics function in GEPIA using the OvCa TCGA dataset (http://gepia.cancer-pku.cn/index.html) [5]. The respective expressions of both genes were normalized to TUBA1A. The Pearson Correlation Scatter Plot is presented in Fig. 5G.

- To determine the transcription factors regulating OvCa spheroidogenesis, we first conducted a keyword-based search in the GeneCards database (https://www.genecards.org/) [6] using the term "ovarian cancer spheroids", which yielded 3,947 associated genes, and then subjected this gene set for transcription factor enrichment analysis using the CHEA 2022 module at the Enrichr platform (https://maayanlab.cloud/Enrichr/) [7]. Data visualization through a clustered heatmap of the top 10 enriched transcription factor programs was generated in Enrichr (Fig. S2).

- To assess the functional connectivity between stemness-associated proteins and EMT regulators implicated in metastatic initiation, protein–protein interaction (PPI) network analysis was constructed using the GeneMANIA website (http://genemania.org) [8]. We used this tool with the official names of genes encoding EMT and stemness proteins as input to construct the PPI network represented in Fig. S14.

- To identify transcription factors regulating CBS-dependent pro-spheroidal proteins, the set of DEGs down regulated in CBS-silenced spheroids were used following proteomic analysis. The top 15 down-regulated proteins were used as input in Harmonizome (https://maayanlab.cloud/Harmonizome/visualize/heat_map/input_genes) [9]. We performed transcription factor enrichment analysis using the ENCODE Transcription Factor Targets dataset to create a heatmap for visualization in Harmonizome of the top 20 enriched transcription factors regulating the gene set [9]. Data represented in Fig. 7F. To predict Putative S-persulfidation sites in SP1, we used pCysMod (http://pcysmod.omicsbio.info/) [11], a deep learning-based platform for cysteine modification analysis. The three human SP1 isoforms (NP_003100.1, NP_001238754.1, and NP_612482.2) each had their full-length amino acid sequences submitted to the pCysMod server. Results revealed that Cys621 and Cys626 for NP_003100.1 (Supplementary Fig. S27A), Cys580 and Cys585 for NP_001238754.1 (Supplementary Fig. S27B), and Cys628 and Cys633 for NP_612482.2 (Supplementary Fig. S27C) were candidate persulfidation sites.

### 2. Proteomics analysis of OvCa monolayers and spheroids

In this manuscript, three types of proteomic datasets were generated and compared as follows:

(i) 2D siControl vs. 2D siCBS,

(ii) 3D spheroids vs. 2D monolayer, and

(iii) 3D siControl vs. 3D siCBS.

The workflow established is presented below.

For both monolayers (2D culture) and spheroids (3D culture), CBS-expressing and CBS-silenced COV318 were trypsinized, pelleted, washed in PBS, repelleted and the cell pellets were submitted for proteomic analysis at the IDeA National Resource for Quantitative Proteomics. Data acquisition and primary analysis were performed by the vendor. We used the normalised and fully annotated dataset provided by the vendor to identify differentially expressed target proteins and genes.

#### (i) Analysis of 2D siControl vs. 2D siCBS

- For heatmap, PCA and Volcano plot visualisation of datasets, we processed and uploaded the normalised data to the MetaboAnalyst platform in the specified format with auto scaling and with log2FC=1.05 and log10p (adj.) <0.05 for 2D siCBS vs 2D siControl (Fig. 2B). The vendor provided volcano plot is presented in Fig. 2C.
- Downregulated DEGs (log2FC=-1.05, log10p (adj.) <0.05) from the same data set were input to transcription factor enrichment analysis using the Harmonizome 'ChEA Transcription factor Targets' dataset (Generated result in Fig. 2D).
- For Enrichment analysis in the 2D siCon vs. 2D siCBS data set, over-representation analysis (ORA) with the integrated cancer hallmark gene set (n=6763) (https://cancerhallmarks.com/) [12] was carried out with significantly downregulated DEGs up to $log_2$(FC) of <-1.05 to identify pathways using 'enrichment plot with distribution' relevant to cancer progression, with p<0.05. The results is presented in 2E.

#### (ii) Analysis of 3D spheroids vs. 2D monolayer,

- For heatmap visualisation, PCA (PLSDA) and Volcano plot, we processed and uploaded the normalised data to the MetaboAnalyst platform in the specified format with auto scaling and with log2FC=1.05 and log10p (adj.) <0.05 for 3D spheroids vs 2D monolayer (Presented in Fig. 5A, 5B).

- ORA with the integrated Cancer hallmark gene set [12] was carried out to identify pathways using 'enrichment plot with distribution' relevant to cancer progression, with $p<0.05$. Pathway analysis and interaction network visualisation were performed using upregulated genes in 3D spheroids vs 2D monolayer using ExpressAnalyst (https://www.expressanalyst.ca/) [13] (Fig. 5C).
- To perform Enrichment analysis, for 2D vs 3D proteomic analysis, we used the vendor-provided volcano plot to extract the upregulated gene/protein list in the 3D spheroids dataset, which was uploaded to the Enrichr database. The generated enrichment plots using different datasets in Enrichr is presented in Fig. 5D and Supplementary figure S18.
- Further, to identify overrepresented Gene Ontology (GO) terms for cellular components, functional enrichment of genes upregulated in 3D spheroids was carried out using the TNM plot functional analysis tool. The top 30 genes from the above-mentioned dataset were used for transcription factor enrichment analysis (Generated enrichment plot presented in Fig. 5F).
- Key transcription factors that may regulate spheroid-associated gene expression programs were identified using the TRRUST Transcription Factor 2019 and ChEA 2022 datasets, accessible on the Enrichr-KG web server (https://maayanlab.cloud/Enrichr/) [14] (Generated enrichment plot presented in Fig. 6A).

**(iii) Analysis of 3D siControl vs. 3D siCBS.**

- For heatmap, PCA and Volcano plot visualisation, we processed and uploaded the normalised data to the MetaboAnalyst platform in the specified format with auto scaling, log2FC=1.05 and log10p (adj.) <0.05 for 3D siControl vs. 3D siCBS**.** The results are depicted in Fig. 7B and 7C.
- To perform pathway enrichment analysis on significantly altered proteins in proteomics dataset from 3D siCon vs 3D siCBS analysis, the NCI Nature 2016 and MSigDB Hallmark databases, on the Enrichr platform were used, and the top 30 CBS-dependent downregulated proteins were subjected to transcription factor enrichment analysis using the ENCODE Transcription Factor Targets dataset. The results are presented as Fig. 7C and Fig. 7D.
- Top 30 downregulated proteins ($log_2FC < -1.05$; log10p (adj.) $< 0.05$) in 3D siCBS (compared to 3D siCon) were used to perform transcription factor enrichment analysis against the ENCODE Transcription Factor Targets dataset, accessible at Harmonizome website (Fig. 7F).

- To identify CBS-dependent proteins potentially regulated by SP1, we performed an integrative analysis combining proteomic and transcription factor target datasets. CBS-dependent proteins were obtained from the 3D spheroid proteomic dataset (siCBS vs siControl). A curated list of SP1 transcriptional targets was retrieved from the Harmonizome ENCODE Transcription Factor Targets (https://maayanlab.cloud/Harmonizome/gene_set/SP1/ENCODE+Transcription+Factor+Targets). The CBS-dependent protein list was intersected against SP1 target gene set to identify overlapping genes representing CBS-dependent SP1-regulated candidates. Functional enrichment analysis of the overlapping gene set was performed using two complementary approaches: TNMplot functional analysis (Supplementary Fig. S29C): Gene Ontology (GO) enrichment was performed using TNMplot by selecting: Tissue: ovarian cancer;

Ontology: Biological Process. Enrichr-based pathway analysis (Fig. D–F): The overlapping gene set was further analysed using Enrichr across multiple libraries, including GO Biological Process 2025 WikiPathways 2024 Human BioPlanet 2019.

**3. Phalloidin staining**

To evaluate the morphological effects of CBS silencing on OvCa monolayers, we stained them with CytoPainter Phalloidin-iFluor 647 Reagent to visualize the actin cytoskeleton. Cells were seeded in 60 mm plates (0.5M cells/plate) with coverslips. The next day, cells were transfected with control scrambled siRNA or siRNA targeting CBS. 96h later, the media was aspirated, and the cover slips were gently collected, rinsed with PBS, and fixed with 4% paraformaldehyde for 10–30 min at room temperature. Cells on the coverslips were incubated for 3–5 min with 0.1% Triton X-100 in PBS for permeabilization and washed again with PBS. 1× phalloidin conjugate solution was applied to the coverslips for 30 min at room temperature, protected from light. The cells were washed twice with PBS, and the coverslips were mounted with VECTASHIELD PLUS Antifade Mounting Medium with DAPI and visualized by fluorescence microscopy using an Ex/Em ~650/665 nm optical filter.

**4. Spheroid formation**

For spheroid formation, cells from 2D monolayers were transferred to poly-HEMA-coated 60 mm plates and incubated them for 48 h. For poly-HEMA coating, 1 mL of poly-HEMA solution in 95% Ethanol (2 mg/ml) was left to dry on the plates overnight. Cells were seeded the next day after a PBS wash. We then collected the cells in spheroids that had survived this anchorage-free culture; spheroids were pelleted at 500 rpm for 2 min, washed in PBS, and counted following Trypan Blue staining. Equal numbers of cells were then transfected with siRNA, either Mission® siRNA Universal Negative Control or siCBS (100 nM), using Lipofectamine™ RNAiMAX Transfection Reagent. Cells in transfection media were distributed to Ultra-low attachment U-bottom 96-well plates (4000 cells/well). Spheroid formation was monitored at specified time-points through 96 h using a bright-field microscope. For rescue experiments with GYY4137, spheroids were treated with 1 mM GYY4137 for the final 24 h, i.e., 72 h after knockdown.

**5. 3D cell viability assay**

10 µL of ready-to-use WST-8 solution was added to each well containing spheroids with 100 µL media and incubated for a 4h. The absorbance, as readout proportional to the number of living cells, was recorded spectrophotometrically at 460 nm using an ELISA plate reader.

**6. Western blot analysis**

Harvested spheroids from suspension cultures were collected, pelleted, washed in PBS and lysed in ice-cold radioimmunoprecipitation assay (RIPA) buffer with freshly added 1% protease-phosphatase inhibitor cocktail by incubation on ice for 30 min with intermittent agitation. Following centrifugation at 13,000 rpm for 10 min at 4˚C, supernatants were collected as spheroid lysates. Volumes corresponding to equal amounts of protein (15 µg) were subjected to Western blotting. Blots were incubated with appropriate primary antibodies, followed by corresponding secondary antibodies. The bands were developed with Clarity™ Western ECL Substrate or Super Signal West Atto and visualized using the Bio-Rad Chemidoc™ Imaging System.

**7. qRT-PCR analysis**

Total RNA was isolated from the spheroids using Trizol™ Reagent according to the manufacturer's protocol. The total RNA concentration and purity was assessed using the Thermo Scientific™ NanoDrop™ OneC Microvolume UV-Vis Spectrophotometer. 1 µg of total RNA was used to synthesize cDNA using the Iscript™ cDNA Synthesis Kit following the manufacturer's protocol. The 10 µL reaction mixture contained 5 µL of Itaq™ Universal SYBR, forward and reverse primers (250 nM each), 1 µL of cDNA template, and nuclease-free water. The relative abundance of mRNA, using 36B4 transcript as an internal reference gene, was expressed by using the comparative cycle threshold method ($2^{-\Delta\Delta CT}$). Primer sequences used in this study were as follows:

| | |
|---|---|
| ITGB1 (Forward): | 5′-GGATTCTCCAGAAGGTGGTTTCG-3′ |
| ITGB1 (Reverse): | 5′-TGCCACCAAGTTTCCCATCTCC-3′ |
| 36B4 (Forward): | 5′-ATGCAGCAGATCCGCATGT-3′ |
| 36B4 (Reverse): | 5′-ATCATGGTGTTCTTGCCCATC-3′ |

**8. Immunofluorescence of spheroids**

To detect CBS, N Cadherin, ITGB1 and SP1 in spheroids, we performed immunofluorescence staining on 5-µm FFPE sections of COV318/A2780-CP20/OVCAR8 spheroids. Briefly, slides with sections from both CBS-expressing and CBS-silenced spheroids obtained after routine microtomy, were used for immunofluorescence studies. Briefly, the slides were deparaffinized using xylene for 5 min, passed through graded alcohol concentrations (100%, 95%, 70% and 50%), washed in running tap water for 10 min and subjected to antigen retrieval by treating with boiling Antigen Retrieval Buffer (1X Tris-EDTA Buffer, pH 9.0) for 20 min in a pressure cooker. Subsequently, the sections were blocked using 2.5% horse serum after washing with TBS twice. The sections were incubated with appropriate primary antibodies overnight. Subsequent steps were performed using VectaFluor™ Excel Amplified Kit (Anti-Rabbit IgG, DyLight® 488) according to the manufacturer's protocol.

**9. Immunofluorescence from Patient-derived ascites samples**

Samples were collected from OvCa patients during paracentesis under an IRB-approved protocol (#2044) at the University of Oklahoma Health Campus. Ascitic cells were pelleted by centrifugation at 400 × g, and red blood cells were removed using an ammonium chloride–based RBC lysis buffer. The remaining cells were washed with PBS and re-pelleted at 400 × g. Cell pellets were fixed in 10% neutral buffered formalin (equivalent to 3.7% formaldehyde in PBS) at 4°C overnight, followed by storage in 70% ethanol at 4°C. For paraffin embedding, fixed cells were resuspended in HistoGel, processed, and embedded as cell blocks for subsequent histological and immunofluorescence analyses.

**10. CD24 staining and flow cytometry**

We conducted flow cytometric detection of CD24+ve cells in OvCa spheroids. Briefly, the spheroids were washed with 1X PBS twice and dissociated with 0.05% trypsin for 5 min. The trypsin was neutralized by addition of serum-supplemented media and centrifuged for 5 min at 1500 rpm. The cell pellet was washed, centrifuged and resuspended in 1X PBS to prepare a single-cell suspension and adjusted to a density of $1 \times 10^6$ cells/100 µL. The cells were stained for viability with 1µL of Ghost Dye™ Violet 510 (Cat# SKU 13-0870-T100; Cytek Biosciences, CA, USA) and incubated for 30 minutes at 2-8°C protected from light. The cells were washed, centrifuged and resuspended in Cell Staining buffer (Cat#420201; BioLegend, CA, USA). The cells were stained with 5µL of Allophycocyanin (APC) conjugated CD24 (Cat#311118; BioLegend, CA, USA) and incubated for 15 minutes in the dark at room temperature. The cells were washed with 3ml of Cell Staining buffer and centrifuged for 5 min at 1500 rpm. The cell pellet was resuspended in 500 µL of Cell Staining buffer for further analysis. Flow cytometry was performed using the Stratedigm-4 cytometer (Stratedigm, Inc., CA, USA), and data analysis was done using FlowJo 10.10 software.

**11. Preclinical model of OvCa transcoelomic metastasis**

To probe the omental metastatic potential of OvCa spheroidal cells, we used murine omental homing assay. For this, we first enriched COV318 spheroids for anoikis-resistant cells by anchorage-independent culture for 48 h and then transfected with siRNA for 92 h as described above (section 4). We then made single-cell suspensions of both CBS-expressing and CBS-silenced spheroids, counted the cells, labelled them with Invitrogen molecular Probes CellTracker Red CMTPX Dye, and injected $2 \times 10^6$ cells intraperitoneally in 6-week-old Nu/J mice (IACUC # 25-007-CH). After 4h, omentum was dissected and examined under a fluorescent microscope. The number of OvCa cells homing to the omentum was quantified as fluorescent area/total omental area.

**12. In silico identification of SP1 binding sites in the ITGB1 promoter**

The human ITGB1 promoter sequence (-2000 to +100 bp relative to the transcription start site, TSS) was obtained from the Eukaryotic Promoter Database (EPD) (https://epd.expasy.org/cgi-bin/epd/get_doc?db=hgEpdNew&format=genome&entry=ITGB1_1). Using the JASPAR database (https://jaspar.elixir.no/), we found possible SP1 binding sites by scanning the promoter sequence with the SP1 position weight matrix (MA0079 variants) at a relative score threshold of 90%. This analysis produced several high-confidence GC-rich SP1 motifs primarily situated within the proximal promoter region. Six primer sets were designed to amplify areas that are rich in SP1 motifs based on the spatial clustering of predicted binding sites. Standard parameters were used to design the primers.

**13. Chromatin immunoprecipitation (ChIP) and qPCR**

ChIP assays were performed using the Magna ChIP™ A/G Chromatin Immunoprecipitation Kit (Millipore, 17-10085) following the manufacturer's protocol. Briefly, cells were crosslinked with formaldehyde and chromatin was sheared to ~200–500 bp fragments using Covaris E220 ultra-sonicator. Sonication was done in microTUBE-130 tubes under the following conditions: peak power: 105.0, duty factor: 5.0, cycles/burst 200, and duration 120 sec. All samples underwent identical sonication conditions to ensure uniformity among biological replicates. Immunoprecipitation was performed using Anti-SP1 antibody (ChIPAb+ Sp1 kit, Millipore, 17-601) and Normal IgG (negative control). Protein-DNA complexes were captured using Protein A/G magnetic beads followed by reverse crosslinking. DNA was purified for downstream analysis. qPCR was performed using SYBR Green with the following primer sets targeting the ITGB1 promoter.

| **Primer Set** | **Target Region (relative to TSS)** | **Forward Primer (5′→3′)** | **Reverse Primer (5′→3′)** |
|---|---|---|---|
| **Primer 1** | **-453 to -343** | GAGGAAGAGGAACTTAAAGAAC | TGCCTCAGTTTCCTCCTCTG |
| **Primer 2** | **-339 to -178** | GACTGTCTCAACGCAGTCCC | CTCCGGAAACGCATTCCTCT |
| **Primer 3** | **-149 to +34** | GATCAGACGCGCAGAGGA | CCTGCTGTTCCGCGACTC |
| **Primer 4** | **-243 to -100** | CATCCTCCGCCTCCTCCTA | CCCCCGGCAGGAAACCA |
| **Primer 5** | **-421 to -255** | CAGGCTACACCAGGAACGC | CGACCGGGACAAAGGAACC |
| **Primer 6** | **-118 to +34** | CTGGTTTCCTGCCGGGG | CCTGCTGTTCCGCGACT |
| **DHFR (Positive control)** | **Known SP1 target** | TCGCCTGCACAAATAGGGAC | AGAACGCGCGGTCAAGTTT |

A validated DHFR promoter primer set (provided with ChIPAb+ Sp1 kit, 17-601) was used as a positive control for SP1 binding.

ChIP enrichment was calculated relative to input DNA and normalized against IgG controls.

**14. Modified Biotin Switch Assay to detect CBS-dependent SP1 persulfidation**

For Modified Biotin Switch Assay, OVCAR8 cells were washed with ice-cold PBS and lysed in HENS buffer supplemented with deferoxamine (100 µM) and protease inhibitors. Lysates were clarified by centrifugation (12,500 rpm, 30 min, 4°C), and protein concentration was determined. Equal amounts of protein (1000 µg) were adjusted in homogenization buffer and treated with methyl methanethiosulfonate (MMTS; 10 M stock) and SDS (final concentration 2%) to block free thiols, followed by incubation at 50°C for 20 min in the dark with intermittent mixing. Proteins were precipitated using chilled acetone, washed with 70% acetone, and resuspended in HENS buffer. An aliquot was reserved as input control. Persulfidated cysteine residues were then labeled by incubation with biotin-HPDP (4 mM stock) for 3 h at room temperature in the dark. Following a second acetone precipitation and resuspension, biotinylated proteins were captured using NeutrAvidin-agarose beads by overnight incubation at 4°C with gentle rotation. Beads were washed twice with HENS buffer, and bound proteins were eluted in Laemmli sample buffer containing β-mercaptoethanol, boiled at 95°C for 5 min, and subjected to immunoblot analysis. For negative control, selected samples were treated with dithiothreitol (DTT, 10 mM) prior to pull-down to abolish persulfidation signals. Input lysates were immunoblotted for SP1, GAPDH, and α-tubulin, with GAPDH serving as an experimental positive control for persulfidation. All experiments were performed using three independent biological replicates (n = 3).

**Table S1. Details of all antibodies used in the study**

| Sl. No. | Antibody name | Company | Catalogue Number | RRIDs | Ab. Dilution (For western blot) | Ab. Dilution (For IF) | Ab. Dilution (For Flow cytometry) |
|---|---|---|---|---|---|---|---|
| 1 | CBS (D8F2P) Rabbit mAb | Cell Signaling Technology | 14782S | AB_2798609 | 1:1000 | 1:400 | NA |
| 2. | CBS Mouse McAb | Proteintech | 67861-1-IG-150UL | AB_2918619 | NA | 1:400 | NA |
| 2 | Anti-OCT4 Rabbit Antibody | Cell Signaling Technology | 2750S | AB_823583 | 1:1000 | NA | NA |
| 3 | Sox2 (D6D9) XP® Rabbit mAb | Cell Signaling Technology | 3579S | AB_2195767 | 1:1000 | NA | NA |
| 4 | Nanog (D73G4) XP® Rabbit mAb | Cell Signaling Technology | 4903S | AB_10559205 | 1:1000 | NA | NA |
| 5 | PARP Rabbit Antibody | Cell Signaling Technology | 9542S | AB_2160739 | 1:1000 | NA | NA |
| 6 | Cystathionine γ-Lyase (D1N1D) Rabbit mAb #19689 (CSE) | Cell Signaling Technology | 19689S | AB_2798824 | 1:1000 | NA | NA |
| 7 | Anti-MPST antibody produced in rabbit, Prestige Antibodies® | Sigma-Aldrich | HPA001240-100UL | AB_1079408 | 1:1000 | NA | NA |
| 8 | Bax Antibody | Cell Signaling Technology | 2772S | AB_10695870 | 1:500 | NA | NA |
| 9 | Bcl 2 (D17C4) Rabbit mAb | Cell Signaling Technology | 3498S | AB_1903907 | 1:1000 | NA | NA |
| 10 | Klf4 (D1F2) Rabbit mAb | Cell Signaling Technology | 12173S | AB_2797840 | 1:1000 | NA | NA |
| 11 | cMyc (D3N8F) Rabbit mAb | Cell Signaling Technology | 13987S | AB_2631168 | 1:1000 | NA | NA |
| 12 | CD44 (E7K2Y) XP® Rabbit mAb | Cell Signaling Technology | 37259S | AB_2750879 | 1:1000 | NA | NA |
| 13 | ZEB1 (E2G6Y) XP® Rabbit mAb | Cell Signaling Technology | 70512S | AB_2935802 | 1:1000 | NA | NA |
| 14 | Anti-SNAI1 Rabbit Polyclonal Antibody | Proteintech | 13099-1-AP | AB_2191756 | 1:1000 | NA | NA |
| 15 | Slug (C19G7) Rabbit mAb | Cell Signaling Technology | 9585S | AB_2239535 | 1:1000 | NA | NA |
| 16 | TWIST1 (E5G9Y) Rabbit mAb | Cell Signaling | 90445S | AB_3064916 | 1:1000 | NA | NA |

| | | | | | | | |
|---|---|---|---|---|---|---|---|
| | | Technology | | | | | |
| 17 | Vimentin (D21H3) XP® Rabbit mAb | Cell Signaling Technology | 5741S | AB_10695459 | 1:1000 | NA | NA |
| 18 | E-cadherin Polyclonal antibody | Proteintech | 20874-1-AP | AB_10697811 | 1:10000 | NA | NA |
| 19 | N-Cadherin (D4R1H) XP® Rabbit mAb | Cell Signaling Technology | 13116S | AB_2687616 | 1:1000 | 1:400 | NA |
| 20 | Anti-Integrin beta-1 Rabbit Polyclonal Antibody | Proteintech | 12594-1-AP | AB_2130085 | 1:5000 | 1:400 | NA |
| 21 | Phospho FAK (Tyr397) (D20B1) Rabbit mAb | Cell Signaling Technology | 8556S | AB_10891442 | 1:1000 | NA | NA |
| 22 | Fak Antibody | Cell Signaling Technology | 3285S | AB_2269034 | 1:1000 | NA | NA |
| 23 | Sp1 (D4C3) Rabbit mAb | Cell Signaling Technology | 9389S | AB_11220235 | 1:1000 | 1:400 | NA |
| 24 | Anti-TUBA1 Mouse Monoclonal Antibody [clone: 1E4C11] | Proteintech | 66031-1-IG | AB_11042766 | 1:20000 | NA | NA |
| 25 | Anti-ACTB Mouse Monoclonal Antibody [clone: 2D4H5] | Proteintech | 66009-1-IG | AB_2687938 | 1:20000 | NA | NA |
| 26 | Anti-CD24 Mouse Monoclonal Antibody (APC (Allophycocyanin)) [clone: ML5] | BioLegend | 311118 | AB_2072735 | NA | NA | 1:20 |

**Table S2. Reagents used in this study**

| | Reagents | Company | Catalogue Number |
|---|---|---|---|
| 1 | Paraformaldehyde solution, 4%, pH 7.0 – 7.6, in 1X PBS | Santa Cruz Technologies | sc-281692 |
| 2 | VECTASHIELD PLUS Antifade Mounting Medium with DAPI | Vector Laboratories | H-2000-10 |
| 3 | RIPA Buffer | Boston BioProducts | BP-115 |
| 4 | Halt™ Protease and Phosphatase Inhibitor Cocktails, (100X) | Thermo Fisher Scientific | PI78446 |
| 5 | Super Signal West Atto | Thermo Fisher Scientific | PIA38555 |
| 6 | CellTracker Red CMTPX Dye | Thermo Fisher Scientific | C34552 |
| 7 | CellTracker Green CMFDA Dye | Thermo Fisher Scientific | C7025 |
| 9 | Lipofectamine™ RNAiMAX Transfection Reagent | Thermo Fisher Scientific | 13778150 |
| 9 | Clarity™ Western ECL Substrate | Bio-Rad | 105061 |
| 10 | iScript™ cDNA Synthesis Kit | Bio-Rad | 1708891 |
| 11 | iTaq™ Universal SYBR | Bio-Rad | 1725122 |
| 12 | WST-8 Assay Kit | Abcam | ab228554 |
| 13 | Antigen Retrieval Buffer (100X Tris-EDTA Buffer, pH 9.0) | Abcam | ab93684 |
| 14 | siCBS: MISSION PRED siRNA 10 OD Desalted (Sequence: CCAUUGACUUGCUGAACUUAAGUUCAGCAAGUCAAUGG) | Sigma-Aldrich | SASI_Hs01_00214623 |
| 15 | Mission® siRNA Universal Negative Control | Sigma-Aldrich | SIC001-10NMOL |
| 16 | Calcein-AM | Sigma-Aldrich | 206700-1MG |
| 17 | Ethidium homodimer | Sigma-Aldrich | 46043-1MG-F |
| 18 | Poly (2-hydroxyethyl methacrylate) (PolyHEMA) | Sigma-Aldrich | P3932-25G |
| 19 | Ghost Dye™ Violet 510 | Cytek Biosciences | SKU 13-0870-T100 |
| 20 | Cell Staining buffer | BioLegend, CA, USA | 420201 |
| 21 | CytoPainter Phalloidin-iFluor 647 Reagent | Abcam | ab176759 |
| 22 | HENS Buffer (HEPES, EDTA, Neocuproine, SDS) | Thermo Fisher Scientific | 90106 |
| 23 | Deferoxamine Mesylate | Sigma-Aldrich | D9533 |
| 24 | Methyl Methanethiosulfonate (MMTS) | Sigma-Aldrich | 208795 |
| 25 | Sodium dodecyl sulfate,BioReagent, suitable for electrophoresis, for molecular biology, ≥98.5% (GC) | Sigma-Aldrich | L3771-100G |
| 26 | Acetone,laboratory Reagent, ≥99.5% | Sigma-Aldrich | 179973-1L |

| | | | |
|---|---|---|---|
| 27 | Biotin-HPDP | Cayman Chemical Company | 16459 |
| 28 | Pierce™ NeutrAvidin™ Agarose, Thermo Scientific, NeutrAvidin™ agarose, standard capacity, Pack Size=5 mL | Thermo Scientific | 29200 |
| 29 | Halt Protease Inhibitor | Fisher Scientific | NC2066967 |
| 30 | Dithiothreitol (DTT) | Goldbio | DTT10 |
| 31 | ChIPAb+ Sp1 - ChIP Validated Antibody and Primer Set, ChIPAb+ Sp1 - ChIP Validated Antibody and Primer Set | Sigma-Aldrich | 17-601 |
| 32 | Magna ChIP™ A/G Chromatin Immunoprecipitation Kit, Magna ChIP™ A/G Chromatin Immunoprecipitation Kit | Sigma-Aldrich | 17-10085 |
| 33 | FLAG-CBS-pcDNA3 plasmid | GenScript (Piscataway, NJ, USA) | OHu26151 |

**Table S3. Clinico-pathological data from in house TMA of 109 OvCa patients**

| Patient ID | H Scores | SURVIVAL EVENT Death=1, Censored=0 | CBS H Score | METASTASIS, Y=1, N=0 |
|---|---|---|---|---|
| OV1 | 0.732382 | 1 | 0.732382 | 1 |
| OV3 | 52.42176 | 0 | 52.42176 | 1 |
| OV4 | 40.79137 | 1 | 40.79137 | 1 |
| OV5 | 27.30815 | 0 | 27.30815 | 1 |
| OV6 | 30.95523 | 1 | 30.95523 | 1 |
| OV7 | 13.05478 | 1 | 13.05478 | 1 |
| OV8 | 0.048317 | 1 | 0.048317 | 1 |
| OV9 | 103.9028 | 1 | 103.9028 | 1 |
| OV10 | 14.64353 | 1 | 14.64353 | 0 |
| OV11 | 0.619215 | 0 | 0.619215 | 1 |
| OV12 | 62.19788 | 1 | 62.19788 | 1 |
| OV13 | 30.1619 | 0 | 30.1619 | 1 |
| OV14 | 3.032678 | 0 | 3.032678 | 1 |
| OV16 | 2.606635 | 0 | 2.606635 | 0 |
| OV17 | 0.068027 | 1 | 0.068027 | 1 |
| OV18 | 5.105526 | 1 | 5.105526 | 1 |
| OV19 | 1.386258 | 1 | 1.386258 | 1 |
| OV20 | 44.18121 | 1 | 44.18121 | 1 |
| OV21 | 0.70559 | 1 | 0.70559 | 1 |
| OV22 | 48.96868 | 1 | 48.96868 | 1 |
| OV23 | 9.160813 | 0 | 9.160813 | 1 |
| OV24 | 0.01471 | 1 | 0.01471 | 1 |
| OV25 | 0.076972 | 1 | 0.076972 | 1 |
| OV26 | 17.1087 | 1 | 17.1087 | 1 |
| OV27 | 9.743237 | 1 | 9.743237 | 1 |
| OV28 | 11.59091 | 1 | 11.59091 | 1 |
| OV29 | 2.601961 | 1 | 2.601961 | 1 |
| OV30 | 0.157134 | 1 | 0.157134 | 1 |
| OV31 | 53.11391 | 1 | 53.11391 | 1 |
| OV34 | 0.235941 | 1 | 0.235941 | 0 |
| OV35 | 14.73961 | 1 | 14.73961 | 1 |
| OV36 | 0.374468 | 1 | 0.374468 | 1 |
| OV38 | 6.109243 | 1 | 6.109243 | 1 |
| OV40 | 23.25066 | 1 | 23.25066 | 1 |
| OV44 | 2.31388 | 1 | 2.31388 | 1 |
| OV45 | 1.35982 | 0 | 1.35982 | 0 |
| OV46 | 3.671785 | 1 | 3.671785 | 0 |
| OV47 | 49.21508 | 1 | 49.21508 | 1 |
| OV48 | 114.3696 | 1 | 114.3696 | 1 |
| OV49 | 1.010536 | 1 | 1.010536 | 1 |
| OV50 | 39.3441 | 1 | 39.3441 | 1 |
| OV51 | 7.062174 | 1 | 7.062174 | 0 |
| OV53 | 13.32529 | 0 | 13.32529 | 1 |
| OV54 | 14.80305 | 1 | 14.80305 | 1 |
| OV55 | 2.629776 | 1 | 2.629776 | 1 |
| OV56 | 1.451562 | 1 | 1.451562 | 1 |
| OV57 | 2.152178 | 0 | 2.152178 | 0 |
| OV58 | 57.54411 | 0 | 57.54411 | 1 |
| OV59 | 0.356404 | 0 | 0.356404 | 1 |
| OV60 | 0 | 1 | 0 | 1 |
| OV61 | 1.090404 | 1 | 1.090404 | 1 |

| | | | | |
|---|---|---|---|---|
| OV62 | 32.49139 | 0 | 32.49139 | 1 |
| OV63 | 0.029603 | 1 | 0.029603 | 1 |
| OV64 | 55.11294 | 1 | 55.11294 | 1 |
| OV65 | 1.834182 | 1 | 1.834182 | 1 |
| OV67 | 1.113606 | 1 | 1.113606 | 1 |
| OV69 | 27.45889 | 1 | 27.45889 | 1 |
| OV70 | 24.78675 | 1 | 24.78675 | 1 |
| OV71 | 39.21162 | 0 | 39.21162 | 1 |
| OV72 | 0.05117 | 1 | 0.05117 | 0 |
| OV73 | 65.22137 | 1 | 65.22137 | 1 |
| OV74 | 0.082546 | 1 | 0.082546 | 1 |
| OV75 | 30.03518 | 1 | 30.03518 | 1 |
| OV76 | 57.17244 | 1 | 57.17244 | 1 |
| OV77 | 1.843111 | 1 | 1.843111 | 0 |
| OV78 | 55.49451 | 0 | 55.49451 | 1 |
| OV79 | 73.17779 | 1 | 73.17779 | 1 |
| OV80 | 13.92182 | 1 | 13.92182 | 1 |
| OV81 | 9.582689 | 0 | 9.582689 | 1 |
| OV82 | 0.306123 | 0 | 0.306123 | 0 |
| OV83 | 13.58475 | 0 | 13.58475 | 1 |
| OV84 | 26.10141 | 1 | 26.10141 | 1 |
| OV85 | 6.95721 | 0 | 6.95721 | 1 |
| OV86 | 0.372373 | 0 | 0.372373 | 1 |
| OV87 | 42.24902 | 1 | 42.24902 | 1 |
| OV88 | 1.358762 | 1 | 1.358762 | 1 |
| OV91 | 0.080677 | 1 | 0.080677 | 1 |
| OV92 | 23.12474 | 1 | 23.12474 | 1 |
| OV93 | 43.79212 | 1 | 43.79212 | 1 |
| OV94 | 89.21015 | 1 | 89.21015 | 1 |
| OV95 | 1.233893 | 1 | 1.233893 | 0 |
| OV96 | 87.29064 | 1 | 87.29064 | 1 |
| OV97 | 30.90284 | 0 | 30.90284 | 1 |
| OV98 | 2.819889 | 1 | 2.819889 | 1 |
| OV99 | 0 | 0 | 0 | 1 |
| OV100 | 0.076191 | 0 | 0.076191 | 1 |
| OV101 | 0.451528 | 1 | 0.451528 | 1 |
| OV102 | 0.1499 | 1 | 0.1499 | 1 |
| OV103 | 0.418656 | 1 | 0.418656 | 1 |
| OV104 | 1.010117 | 0 | 1.010117 | 1 |
| OV105 | 0.430657 | 1 | 0.430657 | 1 |
| OV106 | 29.07972 | 1 | 29.07972 | 1 |
| OV107 | 0.819904 | 0 | 0.819904 | 1 |
| OV108 | 4.335813 | 0 | 4.335813 | 1 |
| OV112 | 0.088387 | 0 | 0.088387 | 1 |
| OV113 | 1.914626 | 1 | 1.914626 | 1 |
| OV114 | 0.968324 | 0 | 0.968324 | 1 |
| OV115 | 11.1067 | 1 | 11.1067 | 1 |
| OV116 | 17.10135 | 0 | 17.10135 | 1 |
| OV117 | 45.41485 | 1 | 45.41485 | 1 |
| OV118 | 13.63064 | 0 | 13.63064 | 1 |
| OV119 | 49.8002 | 1 | 49.8002 | 1 |
| OV120 | 2.333841 | 1 | 2.333841 | 1 |
| OV121 | 22.40777 | 0 | 22.40777 | 1 |
| OV122 | 6.275235 | 1 | 6.275235 | 1 |

| | | | | |
|---|---|---|---|---|
| OV123 | 0.501728 | 1 | 0.501728 | 1 |
| OV125 | 0.253618 | 1 | 0.253618 | 1 |
| OV126 | 2.269478 | 0 | 2.269478 | 1 |

**Table S4. Details of cell lines and their respective cell culture conditions**

| Sl. No. | Cell lines | Media | Supplement | Source |
|---|---|---|---|---|
| 1. | FTE187 | 1:1 Medium 199 and MCDB105 medium with | 10% fetal bovine serum (FBS) + 1% Corning® Penicillin-Streptomycin | Gift from Dr. Jinsong Liu (MD Anderson Cancer Center, Houston, TX) |
| 2. | FTE188 | 1:1 Medium 199 and MCDB105 medium with | 10% fetal bovine serum (FBS) + 1% Corning® Penicillin-Streptomycin | Gift from Dr. Jinsong Liu (MD Anderson Cancer Center, Houston, TX |
| 3. | OSE | 1:1 Medium 199 and MCDB105 medium with | 10% fetal bovine serum (FBS) + 1% Corning® Penicillin-Streptomycin | Gift from Dr. V. Shridhar (Mayo Clinic, Rochester, MN). |
| 4. | HOSE | 1:1 Medium 199 and MCDB105 medium with | 10% fetal bovine serum (FBS) + 1% Corning® Penicillin-Streptomycin | Gift from Dr. V. Shridhar (Mayo Clinic, Rochester, MN). |
| 5. | OVCAR8 | RPMI 1640 Medium with HEPES, L-glutamine | 10% fetal bovine serum (FBS) + 1% Corning® Penicillin-Streptomycin | Purchased from the DCTD Tumor Repository, National Cancer Institute at Frederick, Maryland. |
| 6. | OVCAR4 | RPMI 1640 Medium with HEPES, L-glutamine | 10% fetal bovine serum (FBS) + 1% Corning® Penicillin-Streptomycin | Gift from Ronny I. Drapkin, formerly at Dana-Farber Cancer Institute, Boston, MA, USA |
| 7. | A2780-CP20 | RPMI 1640 Medium with HEPES, L-glutamine | 10% fetal bovine serum (FBS) + 1% Corning® Penicillin-Streptomycin | Gift from Dr. Anil K. Sood, MD Anderson Cancer Center, Houston, TX |
| 8. | COV318 | RPMI 1640 Medium with HEPES, L-glutamine | 10% fetal bovine serum (FBS) + 1% Corning® Penicillin-Streptomycin | Purchased from Sigma-Aldrich (St. Louis. MO) |
| 9. | TYK-nu | MEM (Minimum Essential Medium) | 10% fetal bovine serum (FBS) + 1% Corning® Penicillin-Streptomycin | Purchased from JCRB Cell Bank (Japanese Collection of Research Bioresources Cell Bank) |
| 10. | TYK-nu.CP-r | MEM (Minimum Essential Medium) | 10% fetal bovine serum (FBS) + 1% Corning® Penicillin-Streptomycin | purchased from JCRB Cell Bank (Japanese Collection of Research Bioresources Cell Bank) |
| 11. | COV362 | DMEM (Dulbecco's Modified Eagle's Medium) | 10% fetal bovine serum (FBS) + 1% Corning® Penicillin-Streptomycin | Purchased from Sigma-Aldrich (St. Louis. MO) |
| 12. | OVCAR3 | RPMI 1640 | 20% fetal bovine serum (FBS) + 1% Corning® Penicillin- | Gifted by Susan K. Murphy (Duke University, Durham, |

| | | | | |
|---|---|---|---|---|
| | | | Streptomycin | NC) |

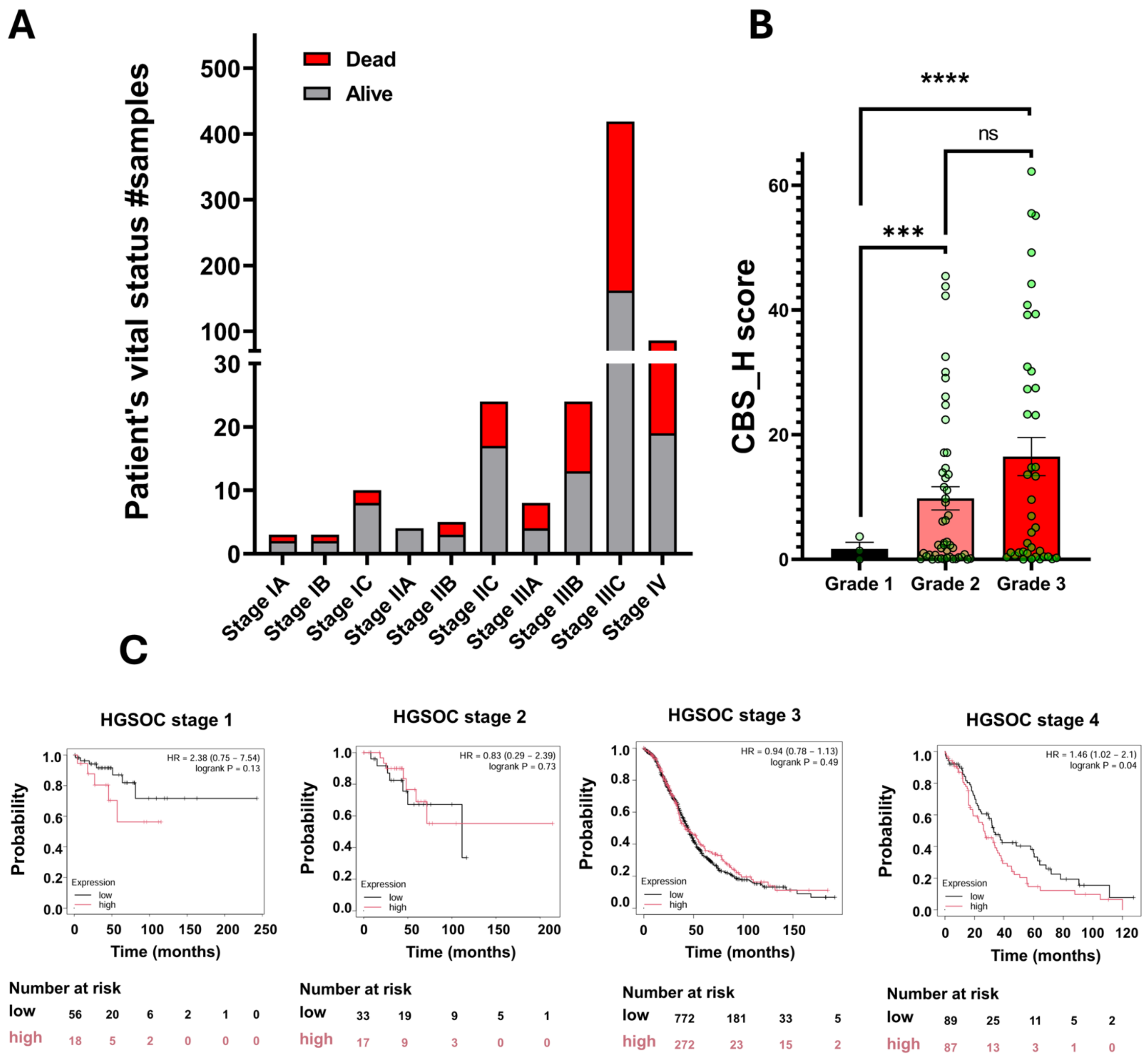


**Figure S1. Association of CBS Expression with Disease Stage and Histopathological Grade.** (A) Bar diagram showing change in the alive-to-dead patient ratio with advancing FIGO stage. (B) Plot showing distribution of CBS H scores across patients classified according to histopathological grades (Result from in house TMA analysis). (C) Kaplan–Meier survival analysis stratified by CBS expression across HGSOC stages (Stages 1–4) showing decline in Overall Survival (OS) probability in high CBS expressing cohort only in Stage 4.

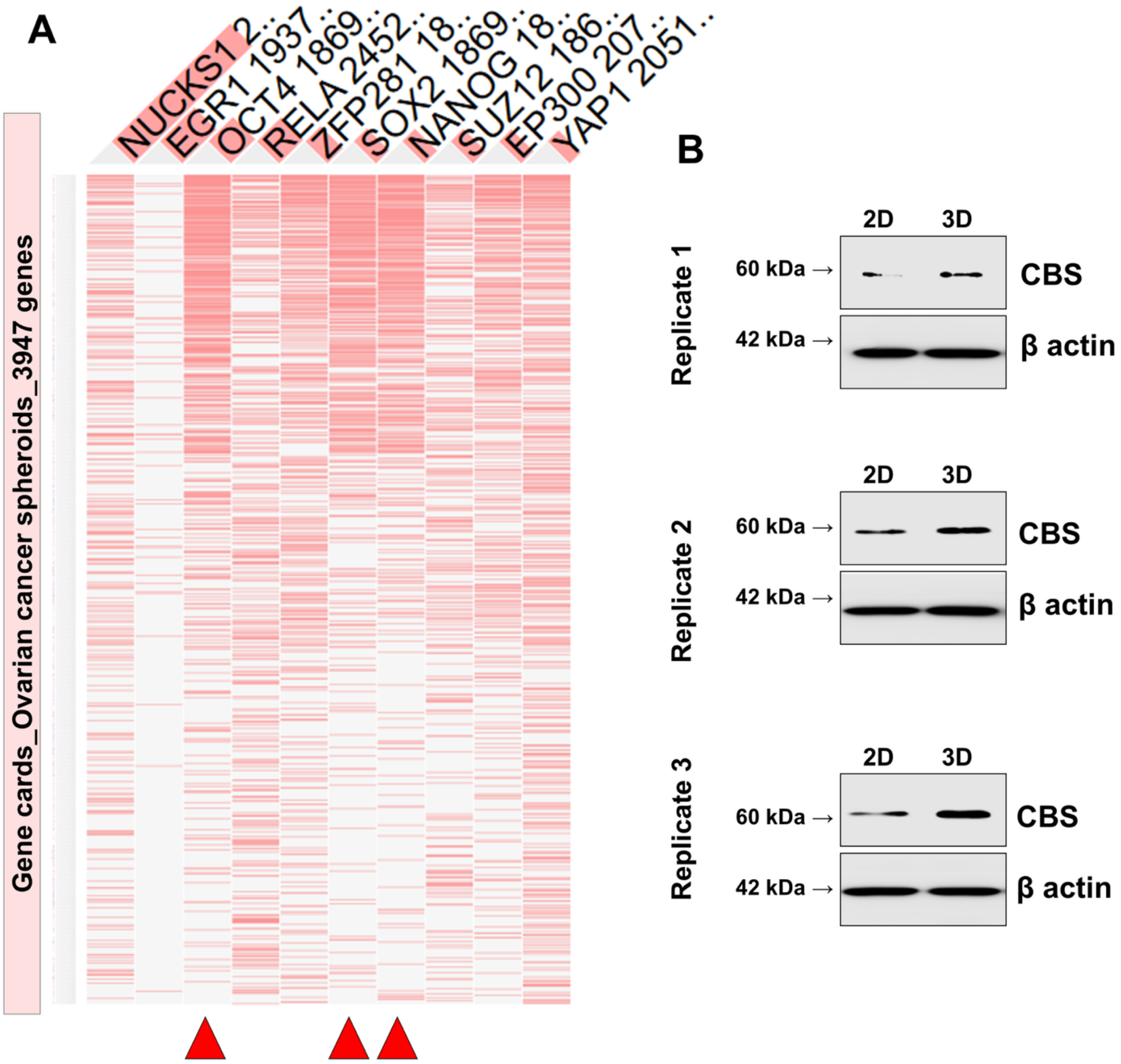


**Figure S2. Integrated identification of key spheroid-associated regulators in OvCa. (A)** OCT4, SOX2 and Nanog as most important pro-spheroidogenic transcription factors in OvCa. Clustergram showing top 10 enriched transcription factors generated through Transcription factor enrichment analysis of genes associated with the keyword "ovarian cancer spheroids". Red arrows indicate Oct4, Sox2 and Nanog. **(B)** Three replicates of immunoblot showing upregulation of CBS in COV318 spheroids.

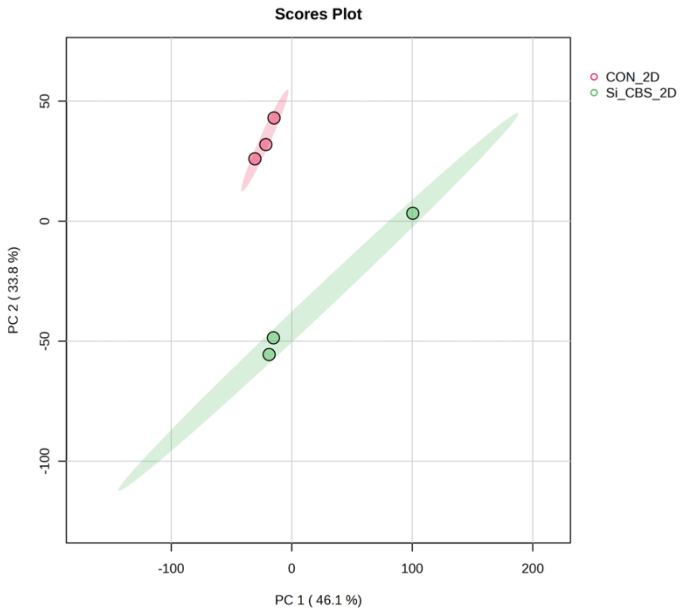


**Figure S3. CBS-expressing and CBS-silenced COV318 monolayers exhibit distinct proteomic profiles.** PCA plot (PLSDA) for 2D siCon vs 2D siCBS conditions (PC1 =46.1%, PC2=33.8%). Proteomic analysis was performed using n = 3 independent biological replicates per condition (siCon and siCBS).

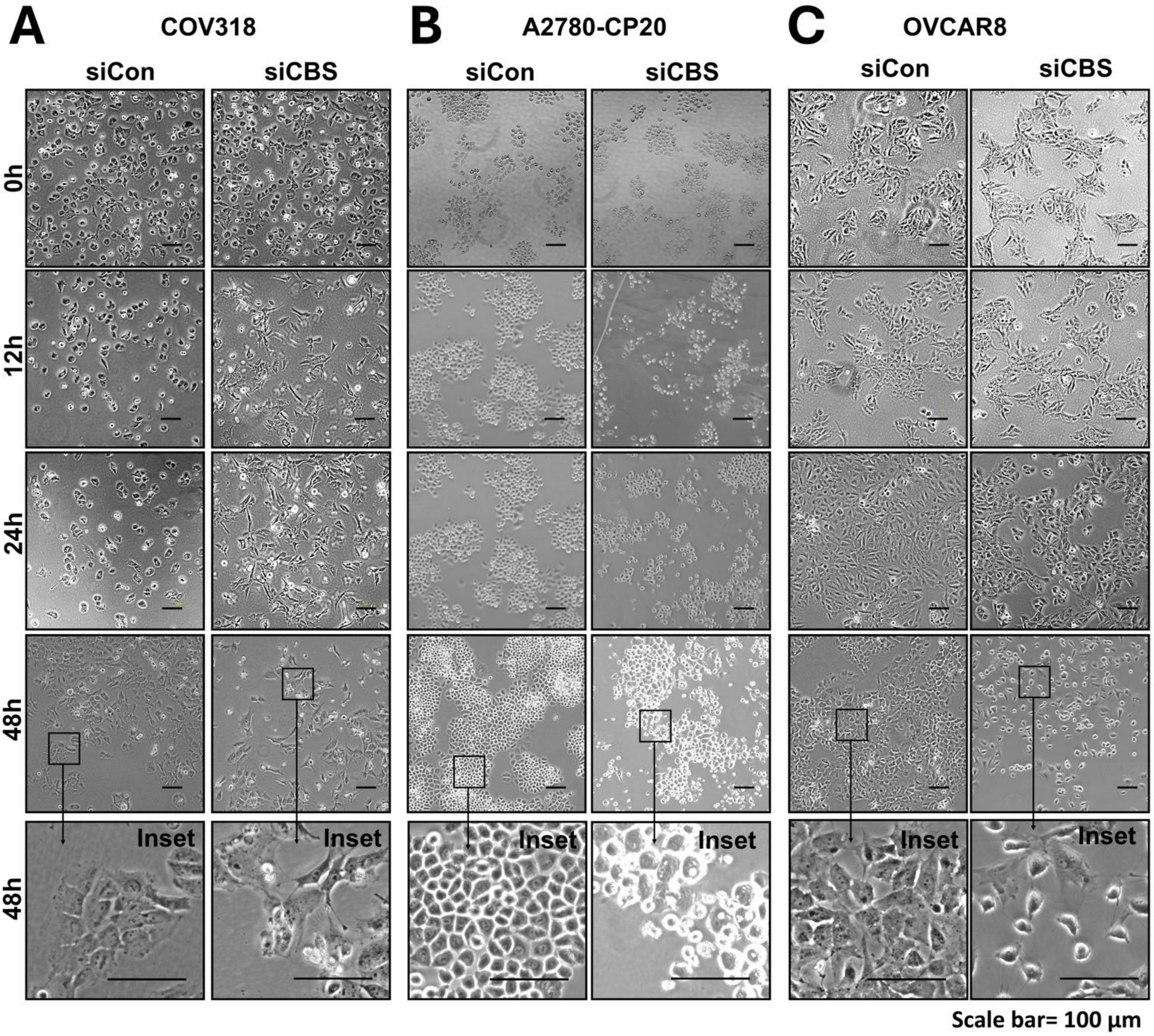


**Figure S4. Morphological alterations of COV318 (A), A2780-CP20 (B), and OVCAR 8 (C) monolayers upon CBS silencing.** Data recorded at different time points through 48 hours (Scale Bar = 100 µm). Inset images show significant morphological alterations upon CBS silencing.

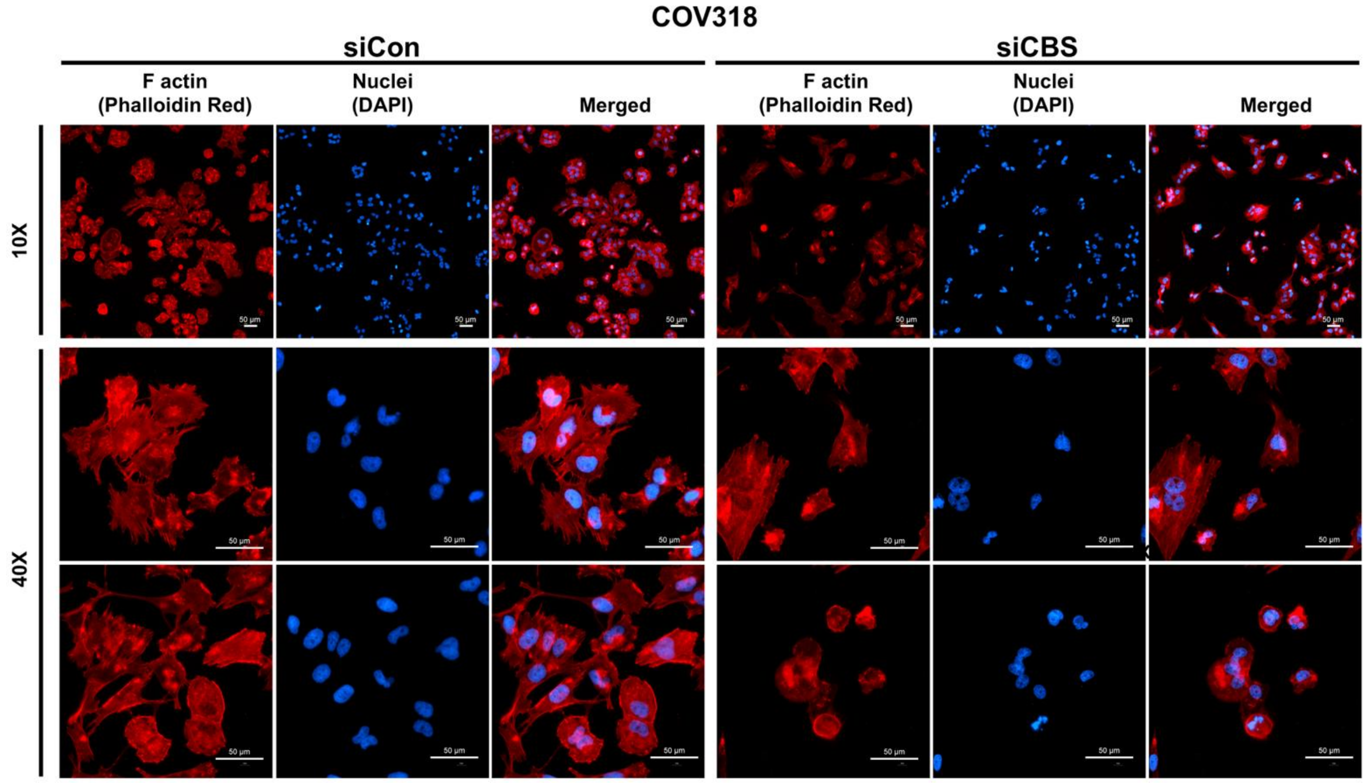


**Figure S5. Alterations in the F-actin cytoskeleton of COV318 monolayers upon CBS silencing.** CBS knockdown caused loss of cell density, shrinkage of cells, increase in the frequency of cells with stress fibers near the plasma membrane, membrane blebbing and shrunk nucleus.

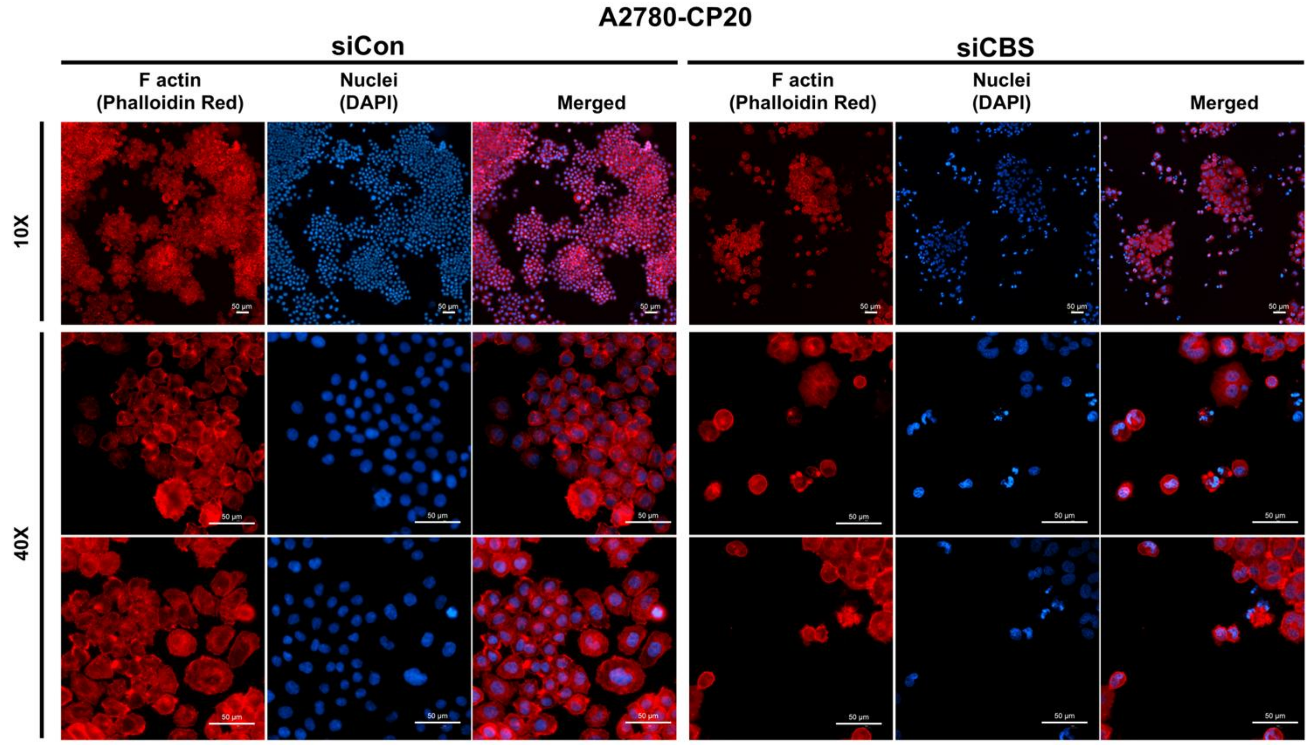


**Figure S6. Alterations in the F-actin cytoskeleton of A2780-CP20 monolayers upon CBS silencing.** CBS knockdown caused loss of cell density, shrinkage of cells, increase in the frequency of cells with membrane blebbing and shrunk nucleus.

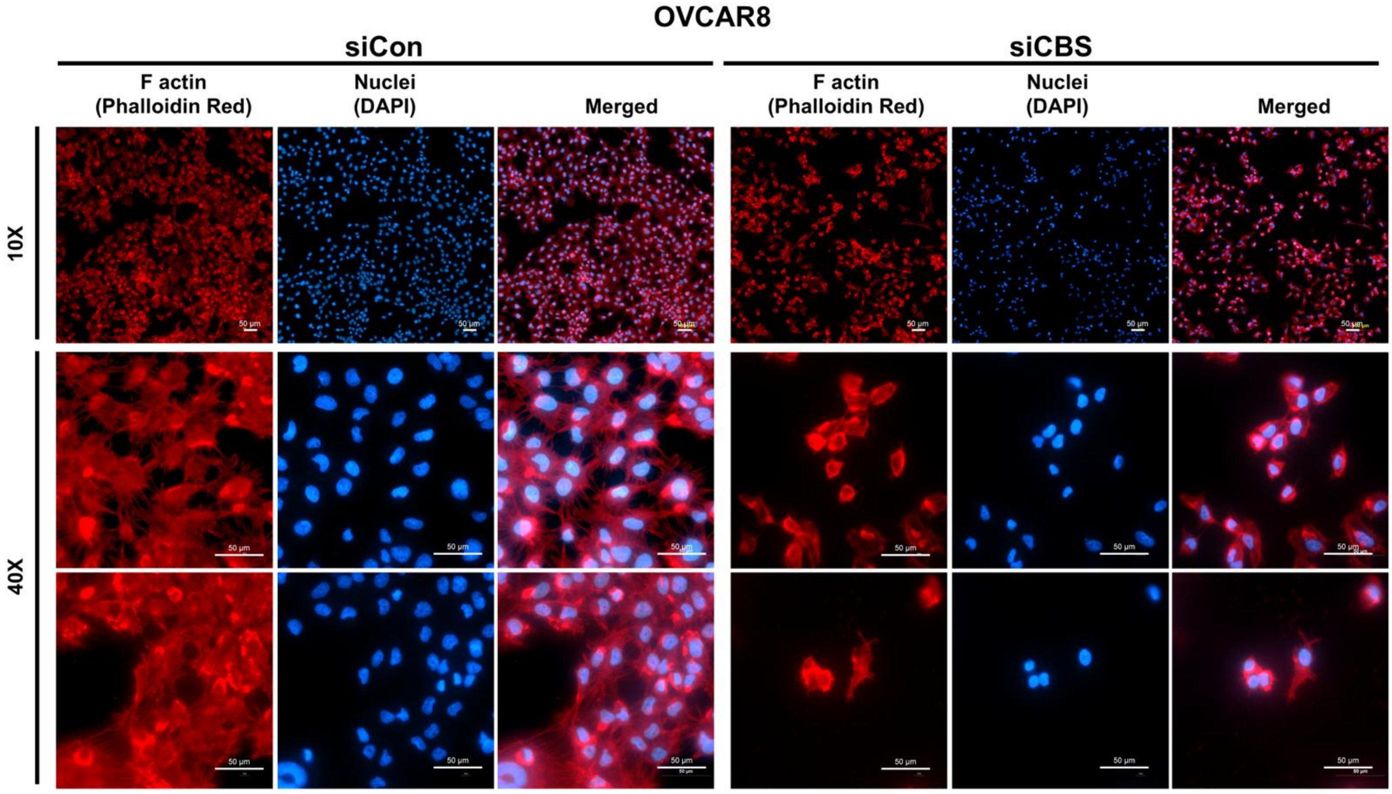


**Figure S7. Alterations in the F-actin cytoskeleton of OVCAR8 monolayers upon CBS silencing.** CBS knockdown caused loss of cell density, shrinkage of cell extensions, induction in the frequency of cells with stress fibers, membrane blebbing and shrunk nucleus.

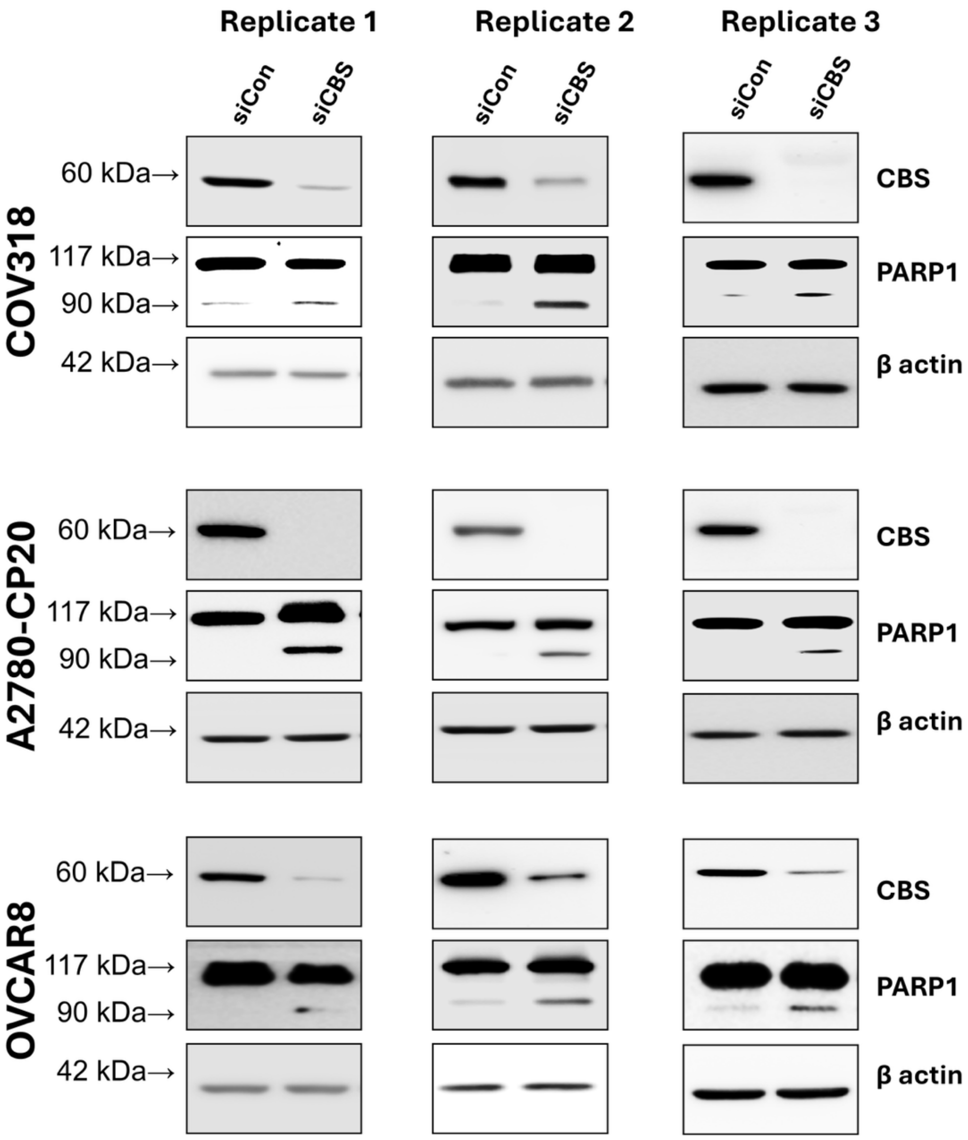


**Figure S8. Induction of apoptosis in CBS silenced OvCa cell monolayers (2D culture).** Western blots from 3 independent biological replicates (n=3) of CBS and PARP1 in CBS-expressing and CBS-silenced monolayers from COV318, A2780-CP20, and OVCAR8 cell lines.

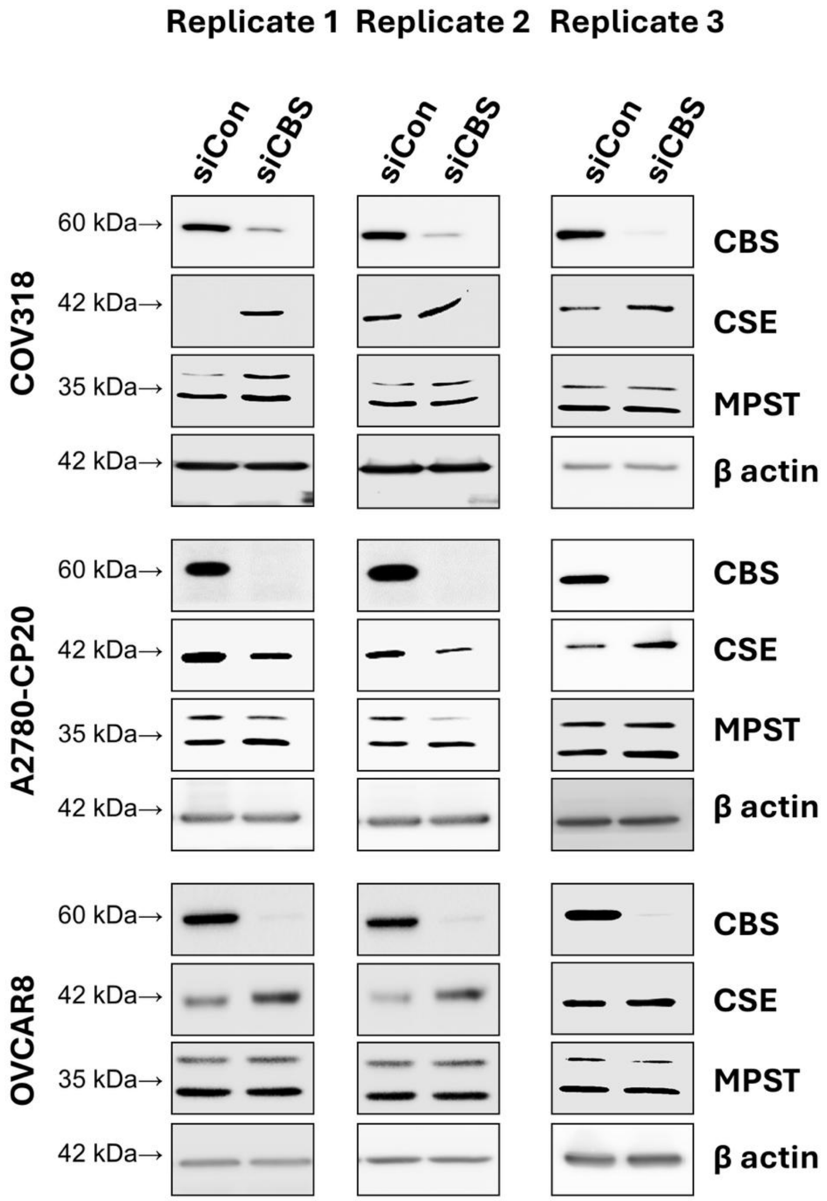


**Figure S9. Effect of CBS knockdown on expression of other sulfhydration protein enzymes in 3D spheroids.** Western blots showing three replicates of CBS, CSE and MPST levels in CBS-expressing and CBS-silenced spheroids (data from 3 independent biological replicates; n=3).

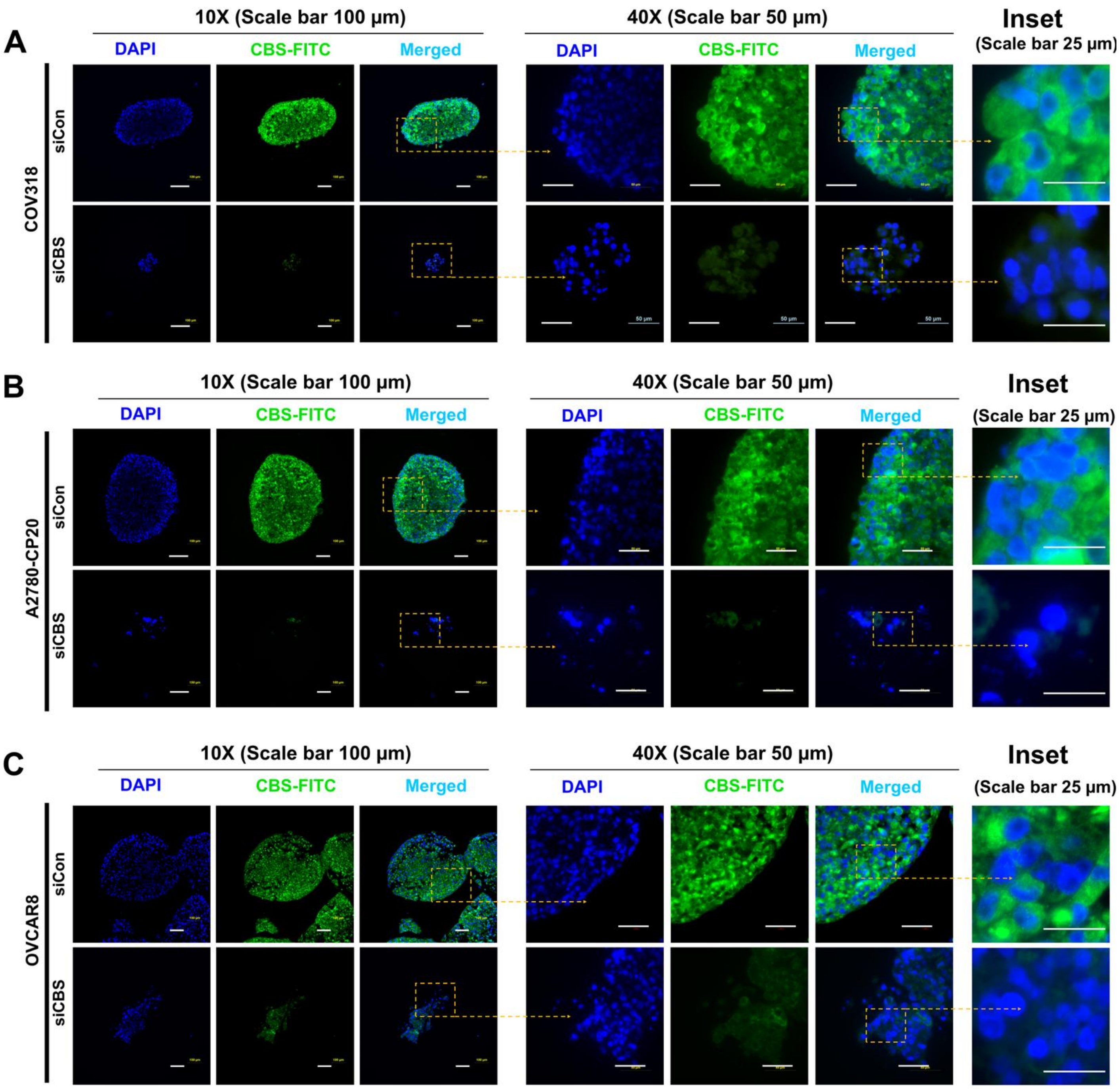


**Figure S10. CBS localization in 3D spheroids.** Immunofluorescence showing intracellular CBS localization in control spheroids and siCBS-treated spheroids of COV318 (A), A2780-CP20 (B) and OVCAR8 (C). Individual channels represent DAPI stained nuclei (Blue) and intracellular CBS (Green).

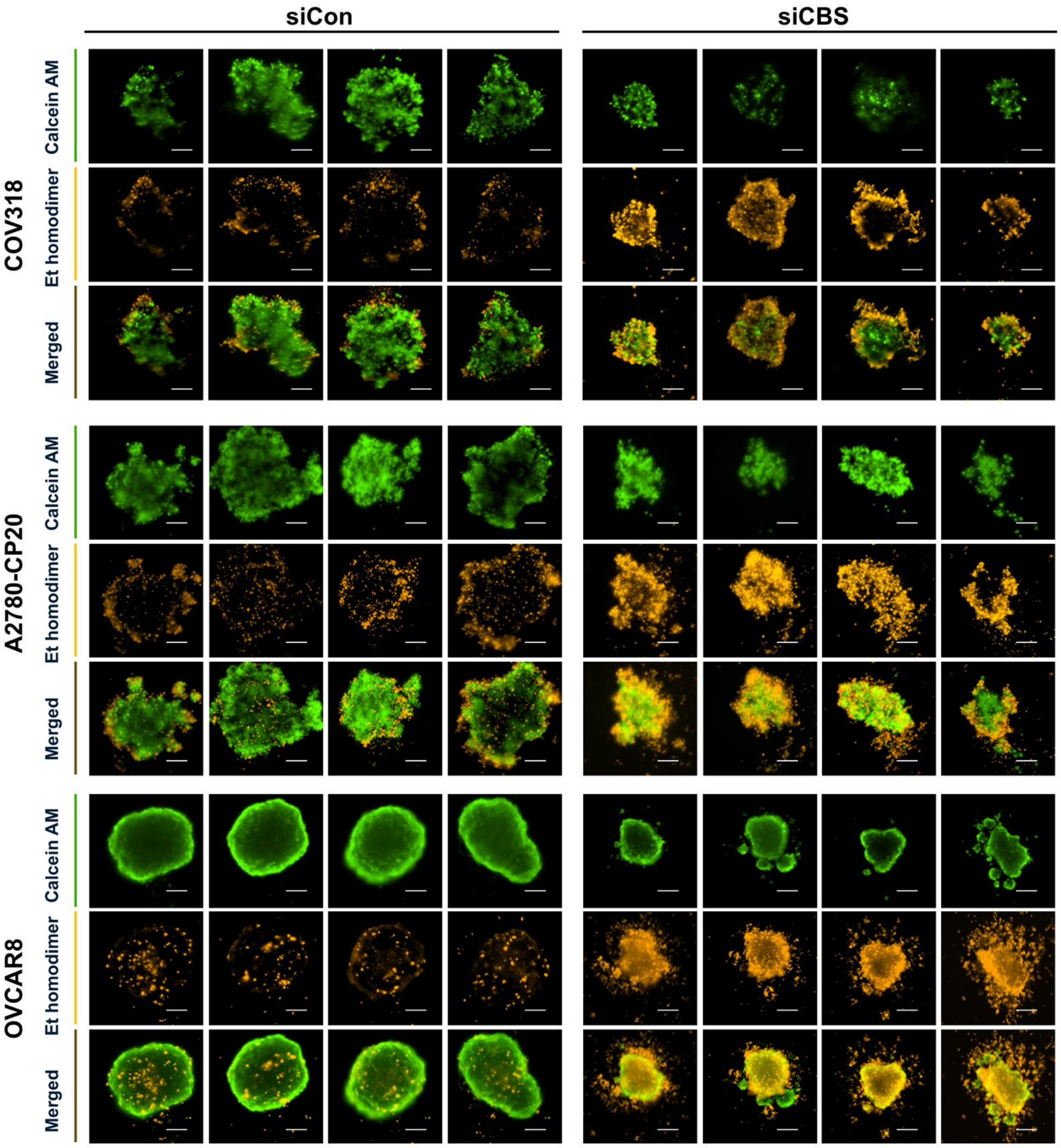


**Figure S11. Abrogation of anoikis resistance in CBS-silenced OvCa spheroids.** Calcein AM–Ethidium homodimer dual staining of CBS-expressing and CBS-silenced OvCa spheroids demonstrating loss of viability in CBS-silenced spheroids (Scale bar = 200 µm)

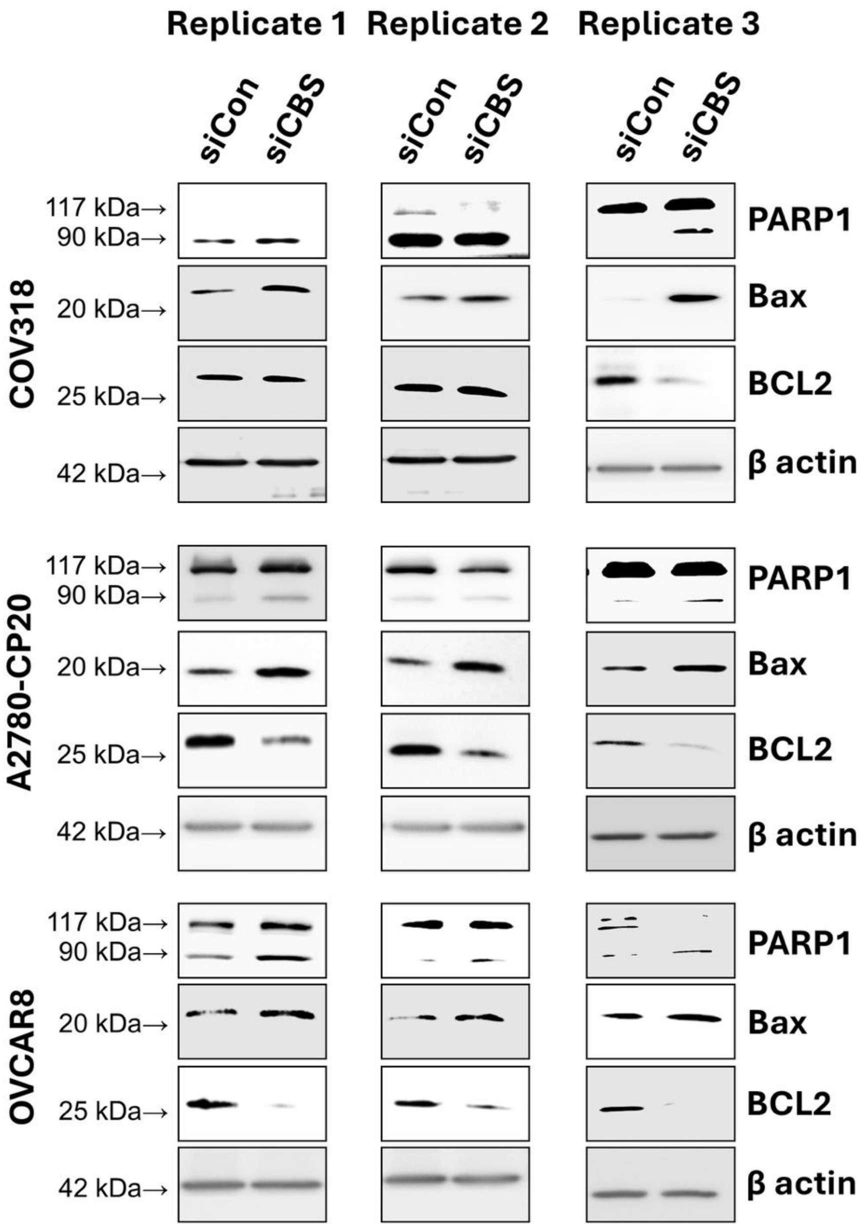


**Figure S12. Induction of apoptotic pathway in CBS silenced spheroids.** Western blots showing three replicates of anoikis markers in CBS-expressing and CBS-silenced spheroids from COV318, A2780-CP20, and OVCAR8 cell lines. Data from 3 independent biological replicates (n=3).

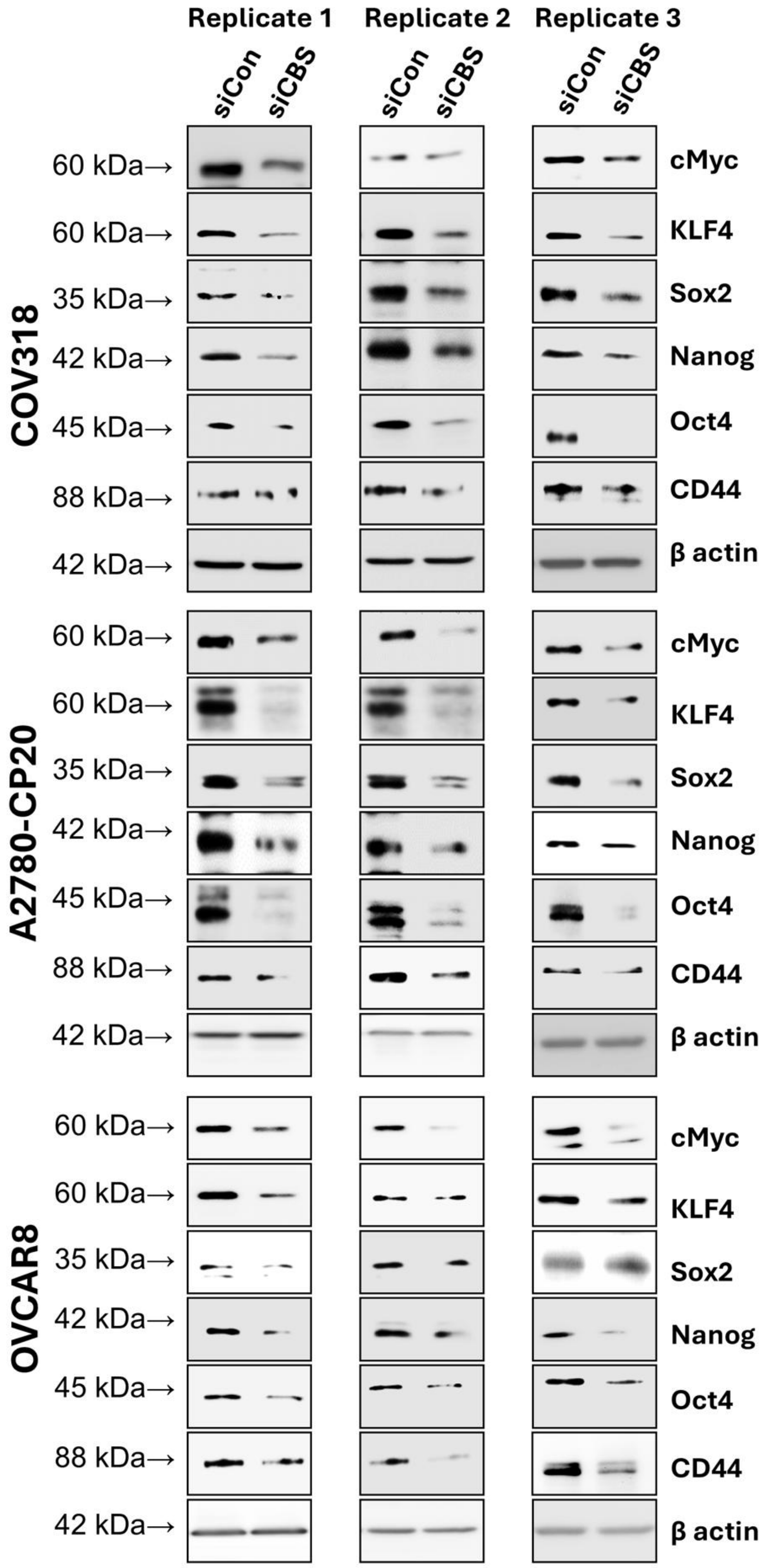


**Figure S13. Attenuation of stemness markers in CBS silenced OvCa spheroids.** Western blots showing three replicates of stemness markers and transcription factors in CBS-expressing and CBS-silenced spheroids from COV318, A2780-CP20, and OVCAR8 cell lines. Data from 3 independent biological replicates (n=3).

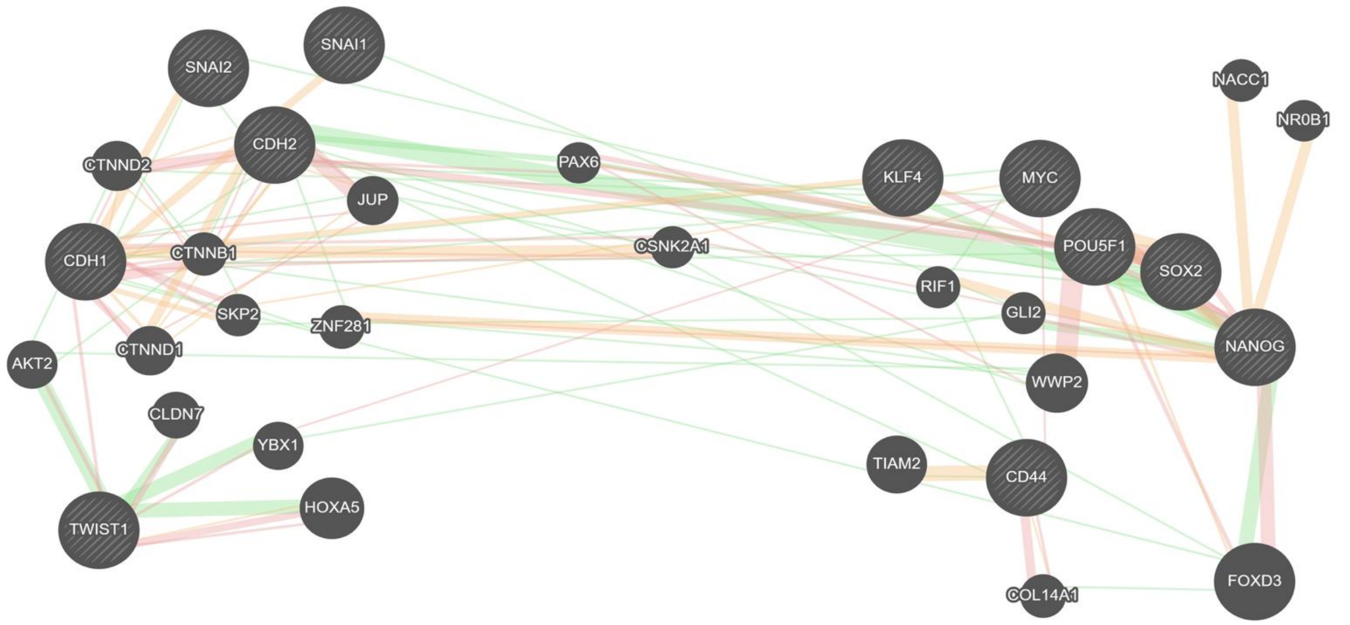


**Figure S14. Stemness contributes to EMT and subsequent OvCa metastasis.** PPI network analysis demonstrated significant associations between stemness-related proteins and EMT regulators implicated in metastatic initiation.

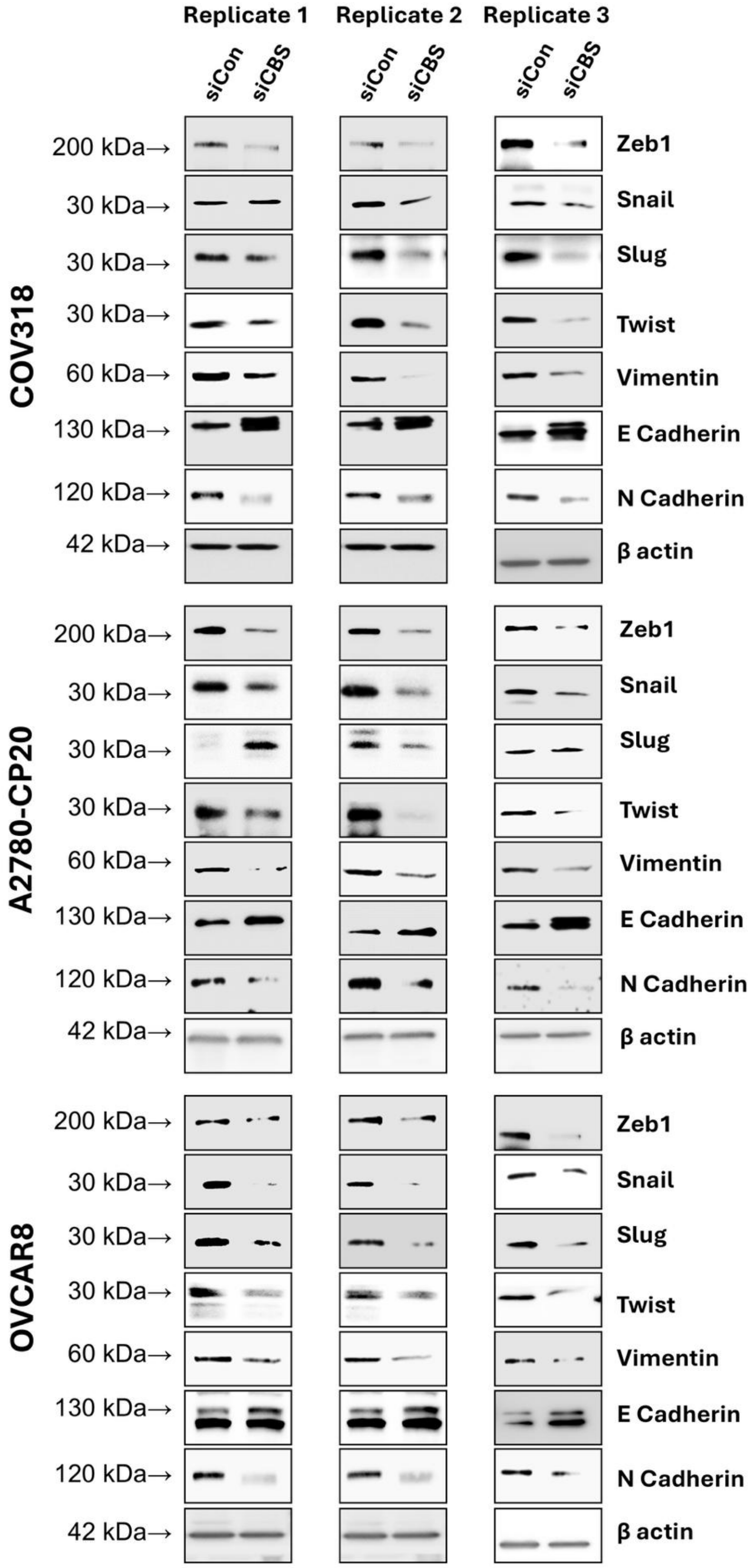


**Figure S15. Attenuation of mesenchymal phenotype in CBS silenced OvCa spheroids.** Western blots showing three replicates of EMT markers and transcription factors in CBS-expressing and CBS-silenced spheroids from COV318, A2780-CP20, and OVCAR8 cell lines. Data from 3 independent biological replicates (n=3).

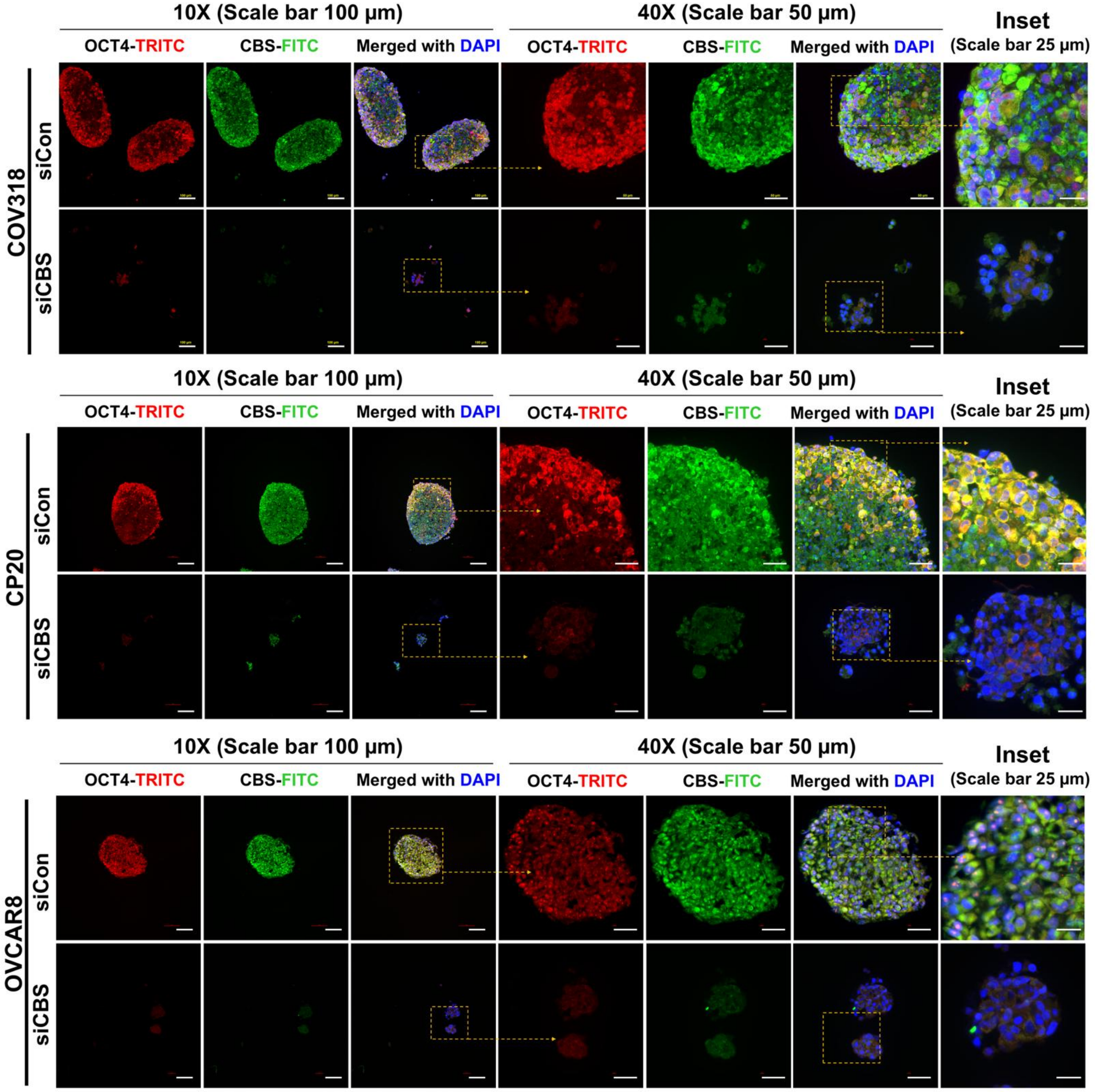


**Figure S16. Attenuation of stemness phenotype in CBS-silenced spheroids.** Oct4 localization and expression in OvCa (COV318, A2780-CP20 and OVCAR8) spheroids as a function of CBS expression. Individual channels represent Oct4 (Red), CBS (Green) and DAPI stained nuclei. In CBS expressing spheroids, cells with high CBS show nuclear localization of Oct4. CBS silencing resulted in spheroids with suppressed Oct4 expression.

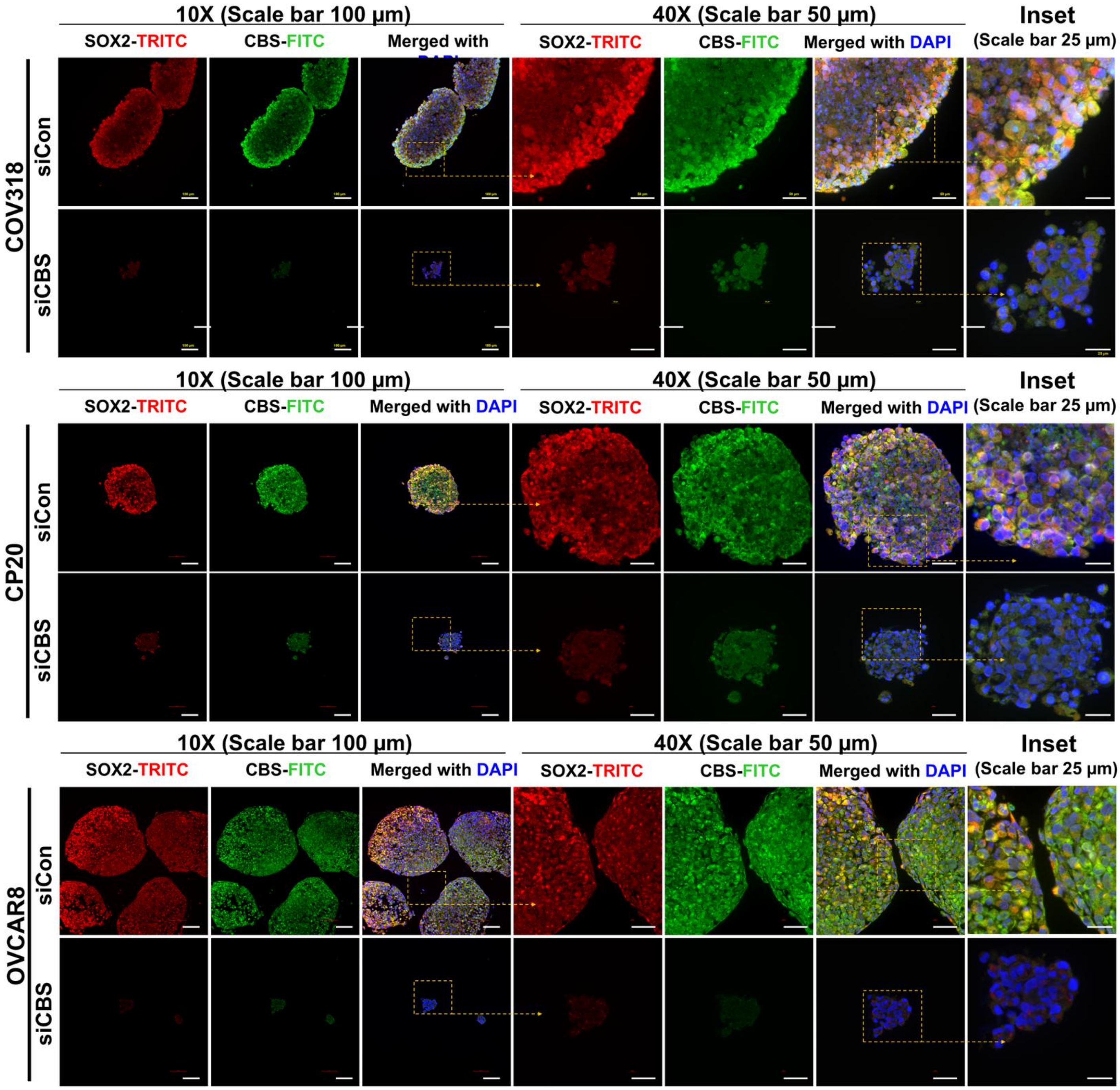


**Figure S17. Attenuation of stemness phenotype in CBS-silenced spheroids.** Sox2 localization and expression in COV318, A2780-CP20 and OVCAR8 spheroids as a function of CBS expression. Individual channels represent Sox2 (Red), CBS (Green) and DAPI stained nuclei.

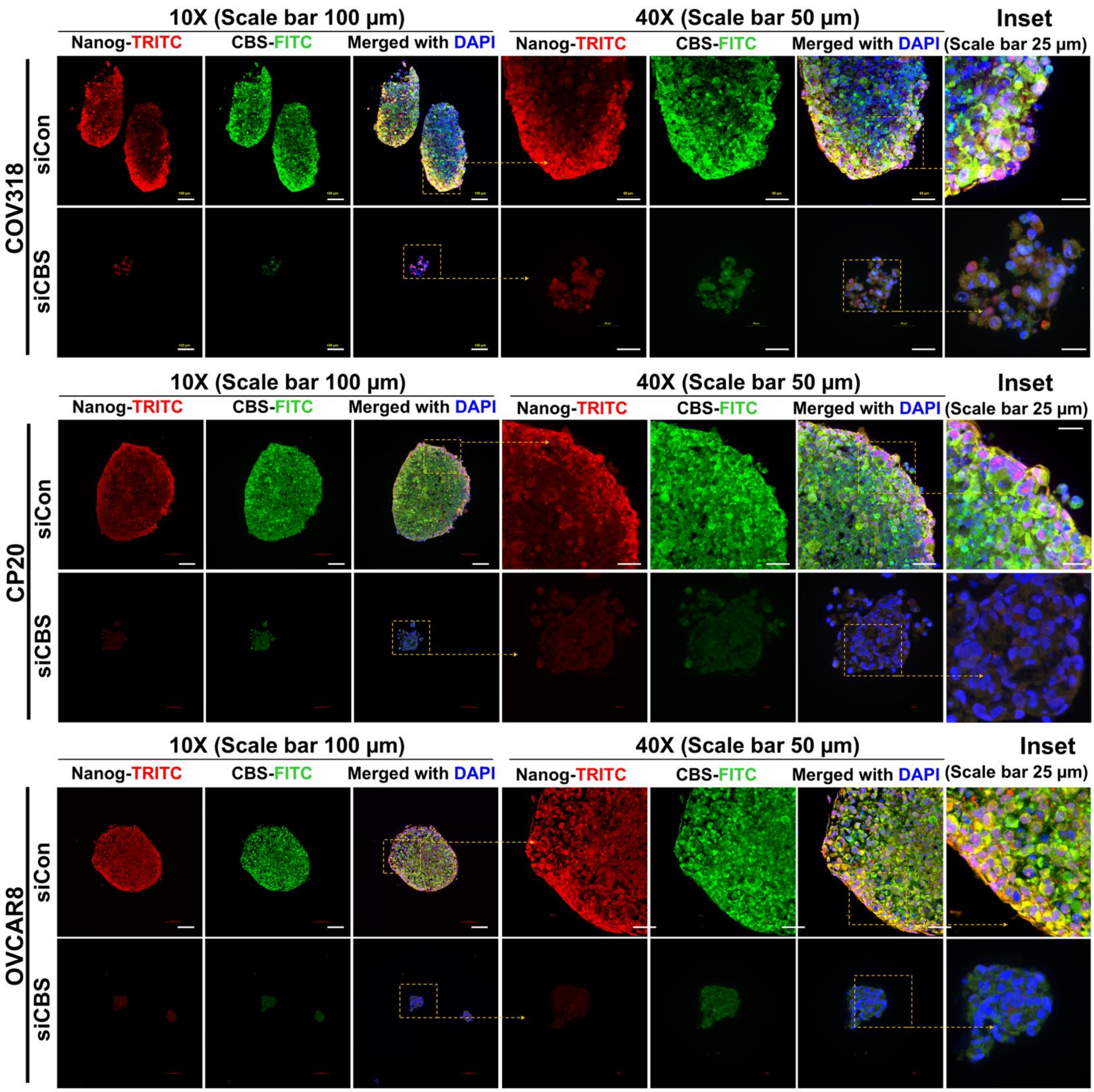


**Figure S18. Attenuation of stemness phenotype in CBS-silenced spheroids.** Localization of Nanog in OvCa (COV318, A2780-CP20 and OVCAR8) spheroids as a function of CBS expression. Individual channels are Nanog (Red), CBS (Green) and DAPI stained nuclei. In CBS expressing spheroids, cells with high CBS show nuclear localization of Nanog. CBS silencing resulted in spheroids with suppressed Nanog expression.

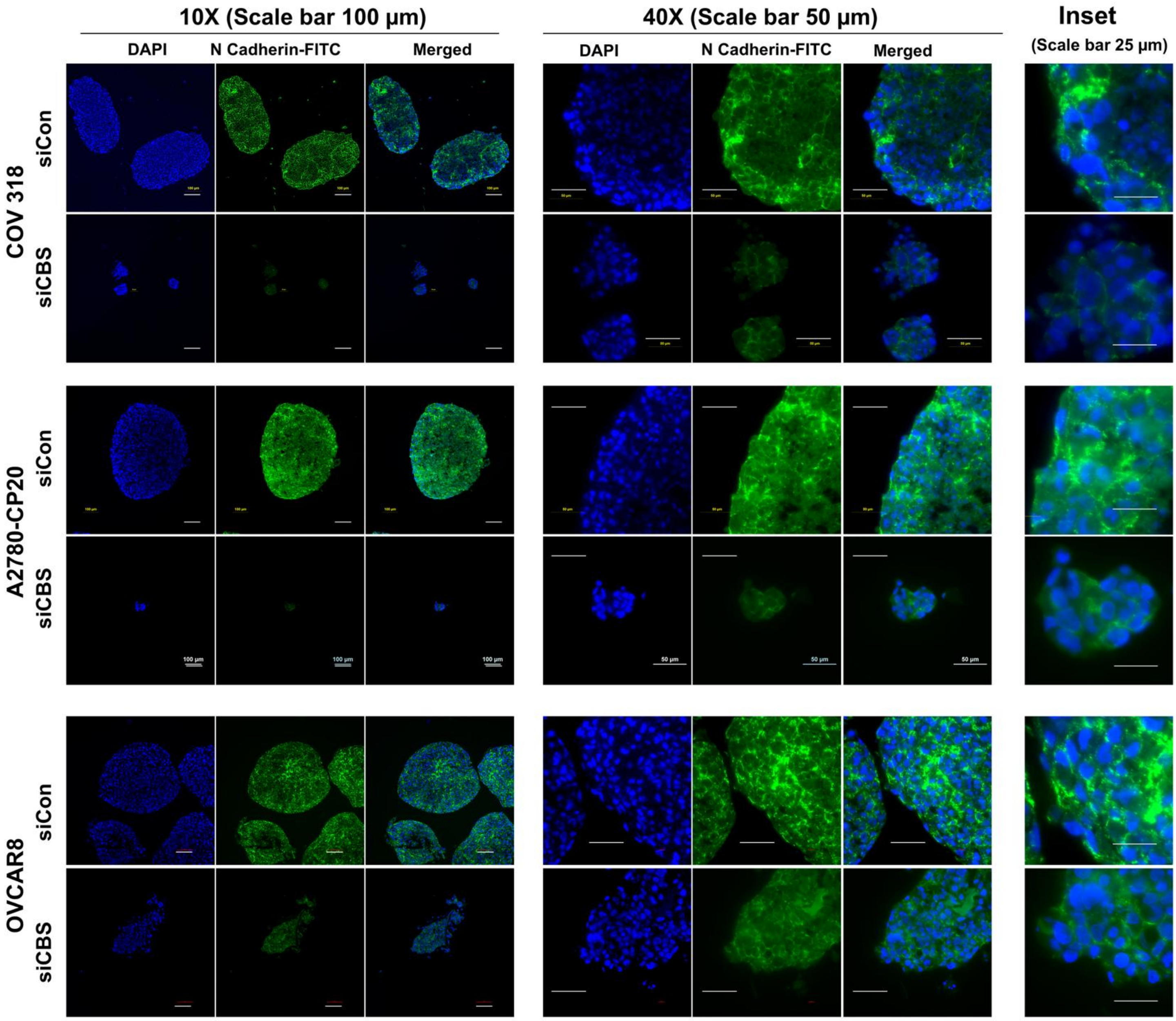


**Figure S19. Attenuation of mesenchymal phenotype in CBS-silenced spheroids.** N-cadherin localization and expression in OvCa (COV318, A2780-CP20 and OVCAR8) spheroids as a function of CBS expression. Individual channels represent DAPI stained nuclei (Blue) and N-Cadherin at the cellular junctions (Green). CBS knockdown resulted in spheroids with reduced mesenchymal attributes.

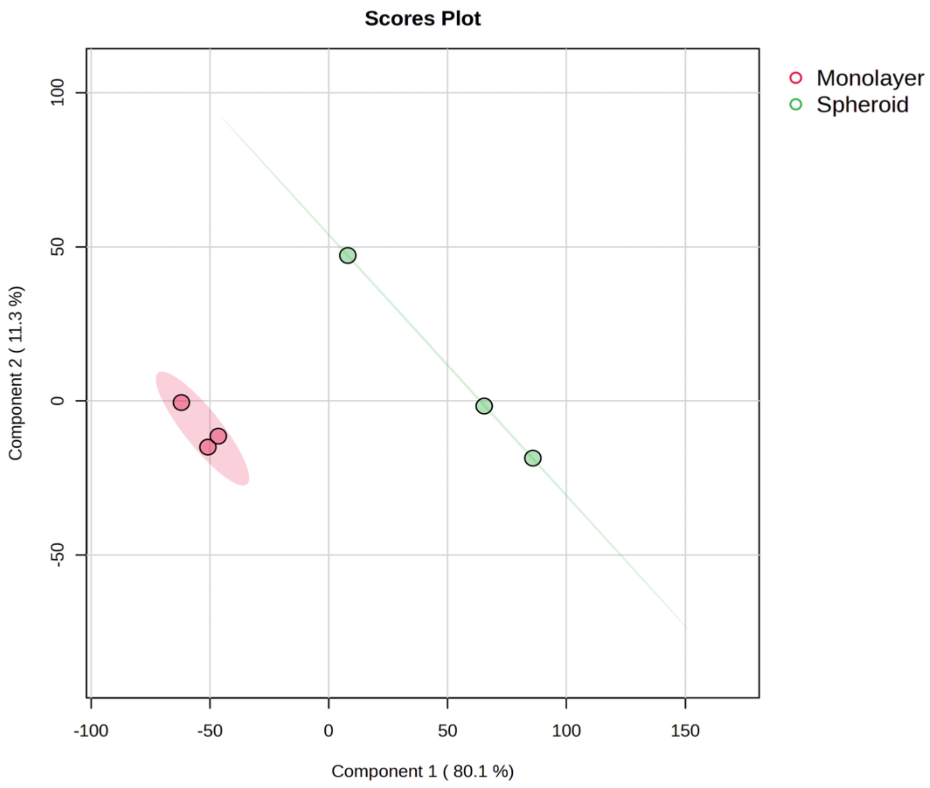


**Fig. S20. COV318 exhibited distinct proteomic profiles as monolayers and spheroids.** PCA plot (PLSDA) for 3D vs 2D conditions (PC1 =80.1%, PC2=11.3%). Proteomic analysis was performed using 3 independent biological replicates (n=3) for monolayers and spheroids each.

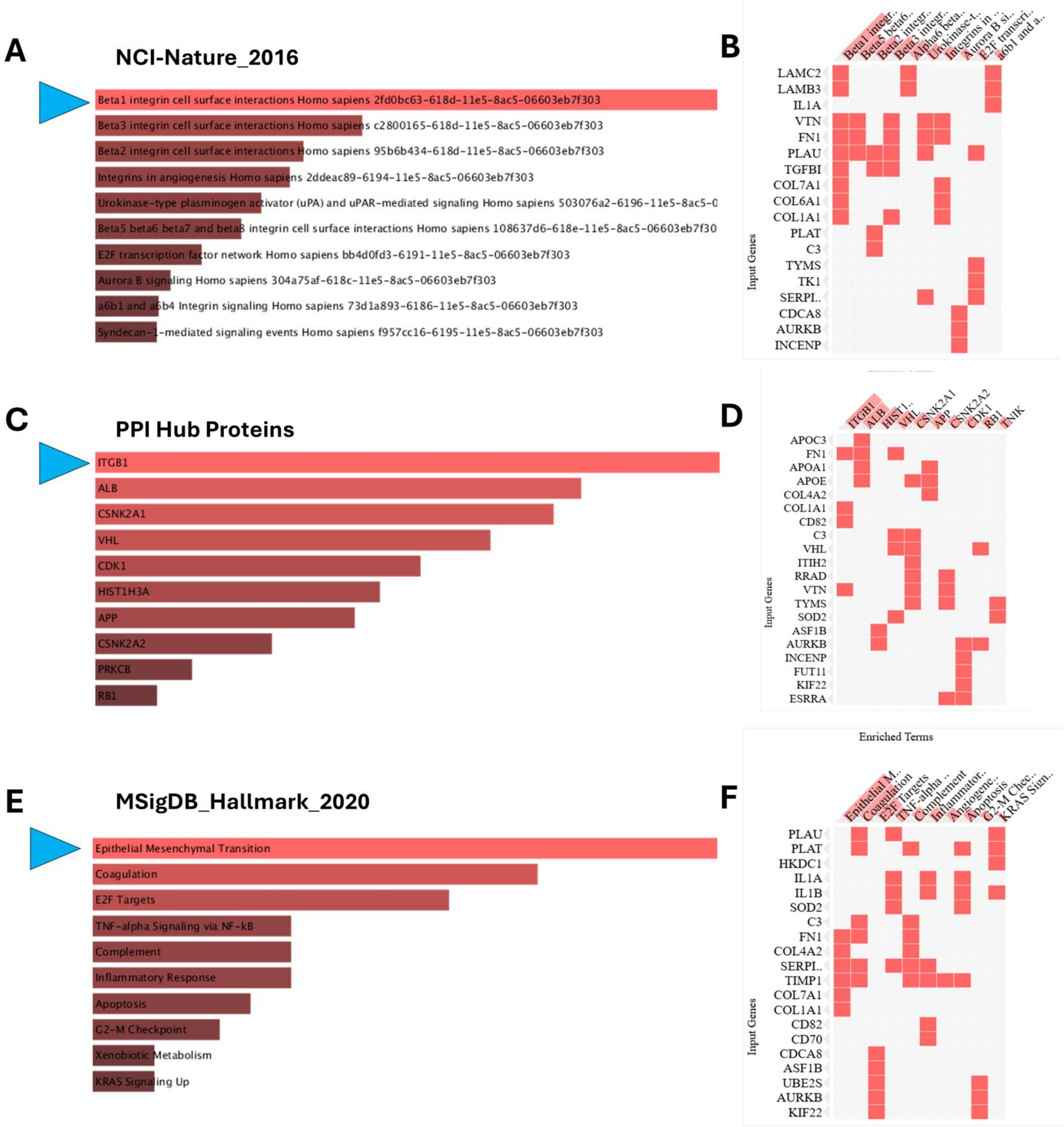


**Figure S21.** Enrichment analysis of upregulated DEGs using NCI-Nature_2016 (A and B) and PPI Hub proteins module (C and D) of Enrichr indicating involvement of ITGB1 in OvCa anchorage independent survival. Similar analysis with MSigDB-Hallmark_2020 module (E and F) reveals EMT to be crucial for 3D survival of anoikis-resistant spheroids.

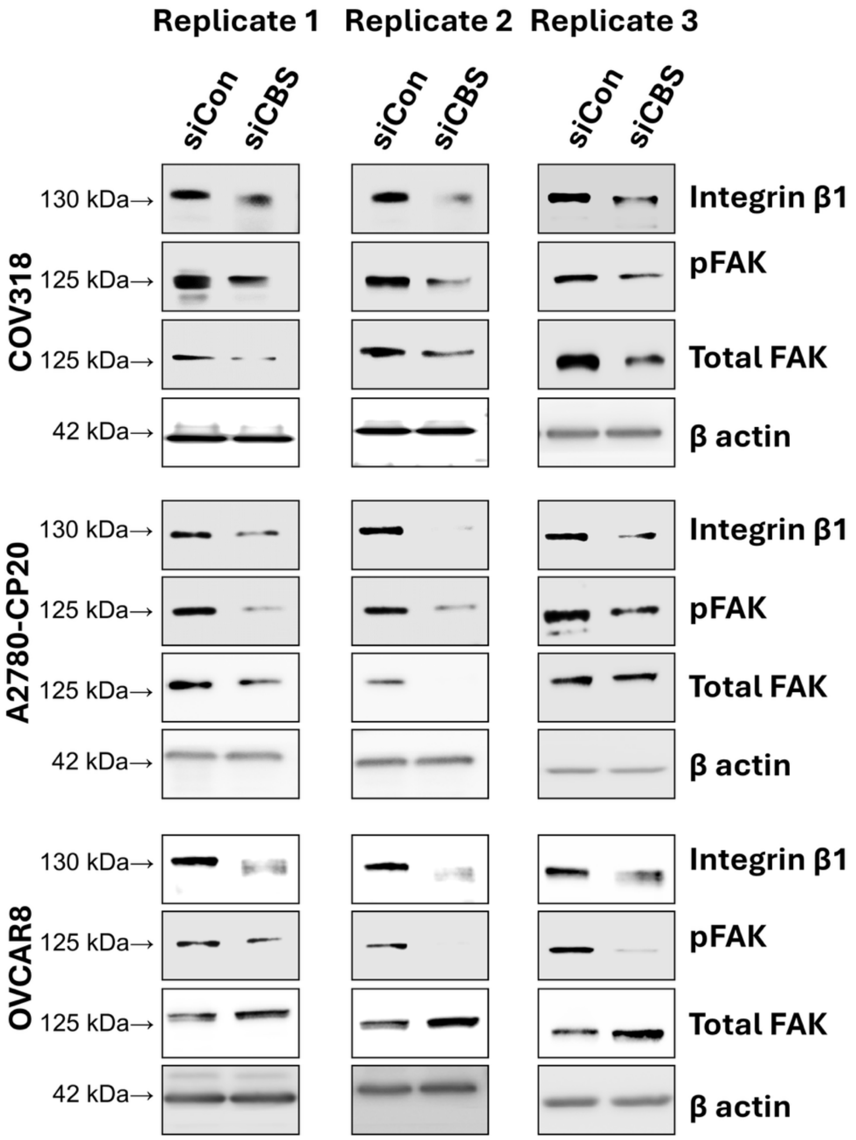


**Figure S22. CBS silencing causes repression of ITGB1 and its downstream effector.** Western blots showing three replicates of ITGB1, pFAK and FAK in CBS-expressing and-CBS silenced spheroids from COV318, A2780-CP20, and OVCAR8 cell lines. Data from 3 independent biological replicates (n=3).

.

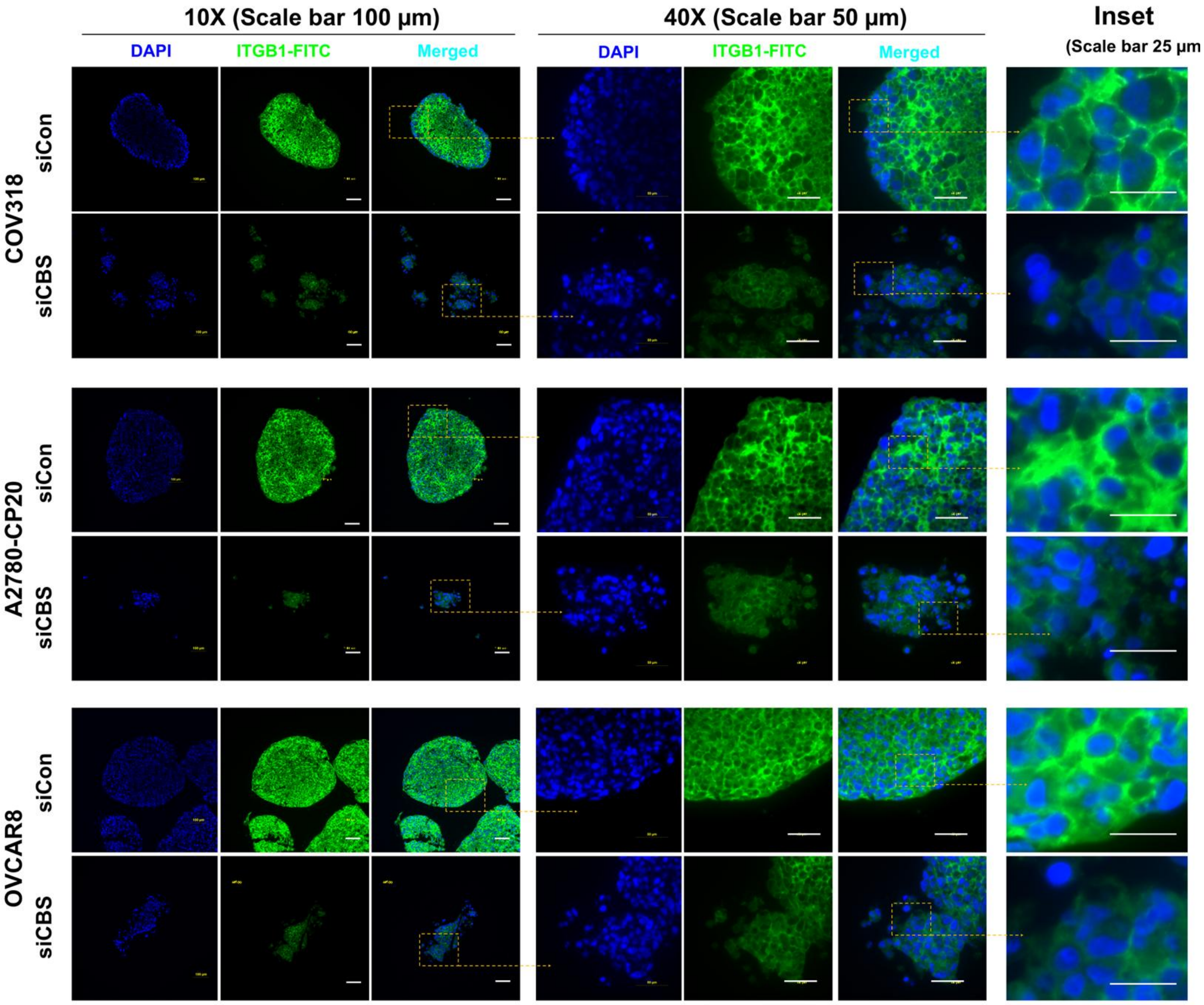


**Figure S23. CBS-silenced spheroids exhibit reduced ITGB1 at spheroidal cell junctions leading to loose spheroids.** ITGB1 localization and expression in OvCa (COV318, A2780-CP20 and OVCAR8) spheroids as a function of CBS expression. Individual channels represent DAPI stained nuclei (Blue) and ITGB1 at the cellular junctions (Green)**.** CBS knock down resulted in loose spheroids with reduced ITGB1 expression.

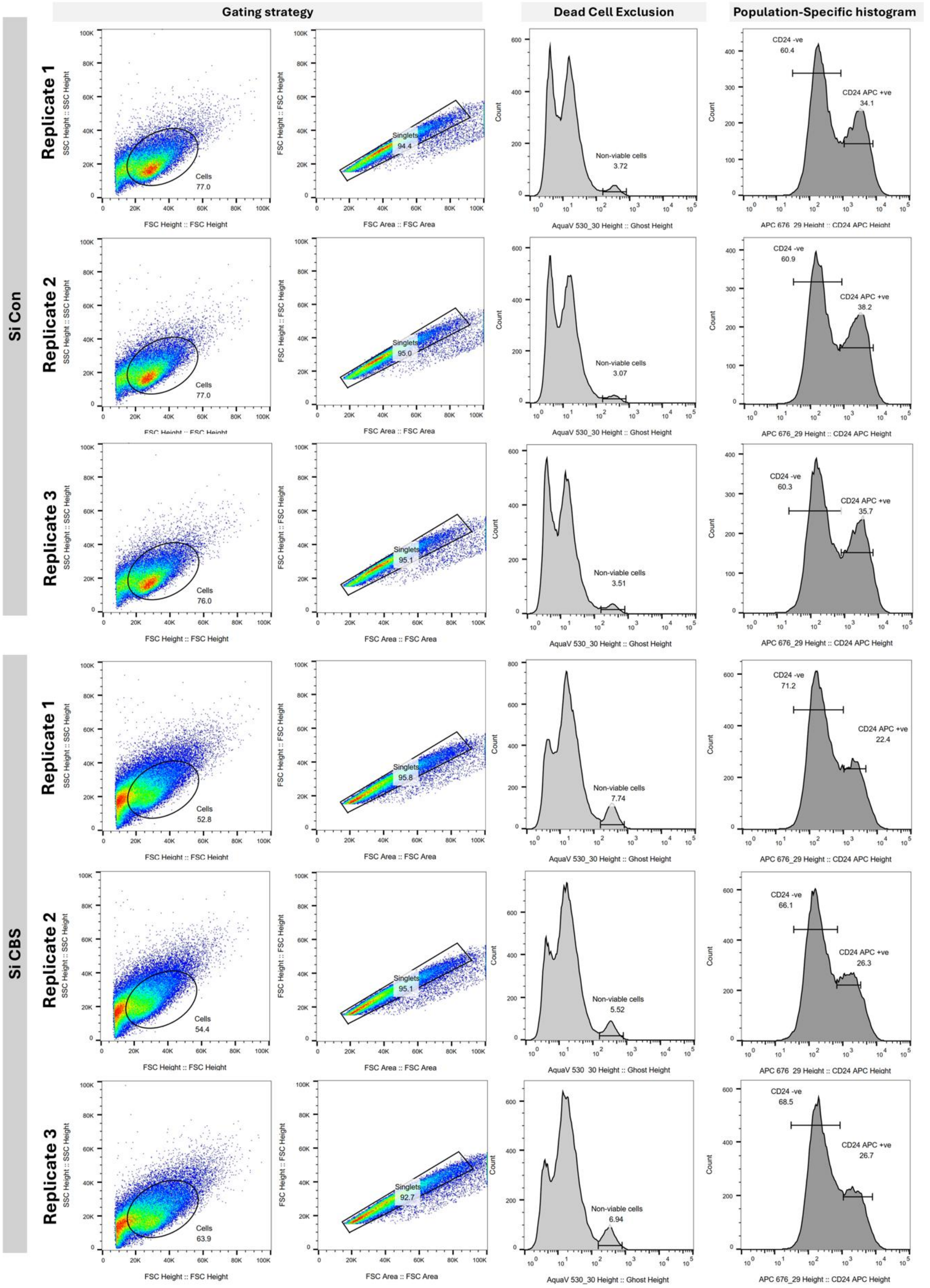


**Figure S24. CBS silencing in spheroids led to a decline in the CD24+ve subpopulation.** Flow cytometry data for CD24 staining in COV318 spheroidal cells also showing gating strategy and dead cell exclusion. Data represents experiments conducted in triplicate (n=3).

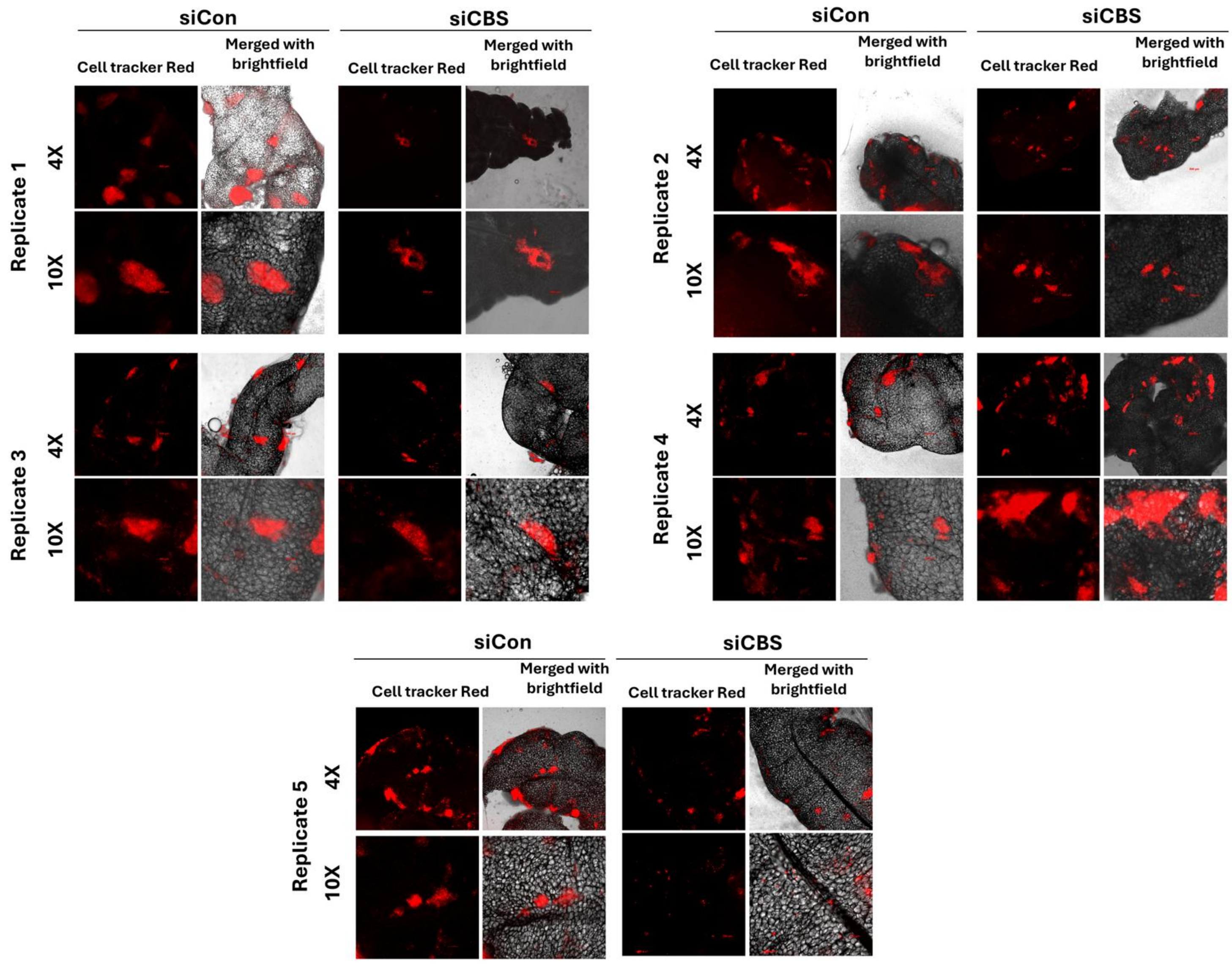


**Figure S25. Loss of metastatic competence in CBS-silenced spheroidal cells.** Photomicrographs of mice omentum (5 replicates; n=5 for each group) showing adhered fluorescently labelled anoikis-resistant COV318 spheroidal cells as a function of CBS knockdown.

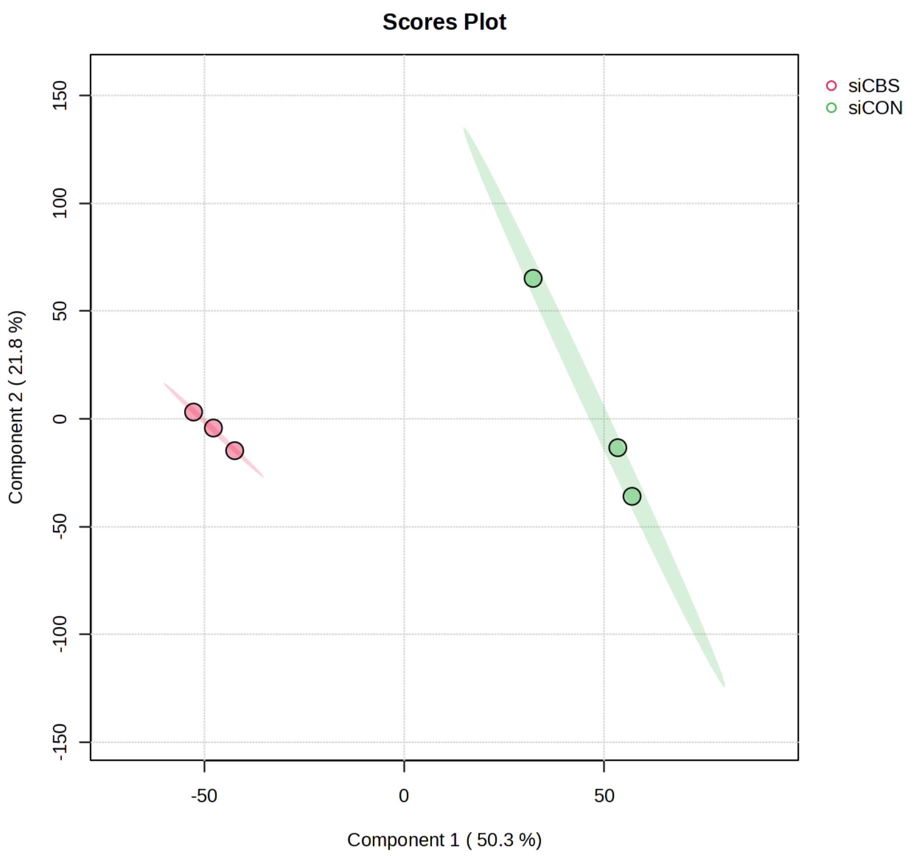


**Figure S26. CBS-expressing and CBS-silenced COV318 spheroids exhibit distinct proteomic profiles.** PCA plot (PLSDA) for 3D siCon vs 2D siCBS conditions (PC1 =50.3%, PC2=21.8%). Proteomic analysis was conducted using data from 3 independent biological replicates (n=3) of CBS silenced and CBS expressing spheroids each.

**A**

#### Sp1 : Sulfhydration

| ID | Position | Modification | FPR | Peptide |
|---|---|---|---|---|
| NP_003100.1 | 621 | S-sulfhydration | 0.79% | GKKKQHICHIQGCGK |
| NP_003100.1 | 626 | S-sulfhydration | 0.10% | HICHIQGCGKVYGKT |

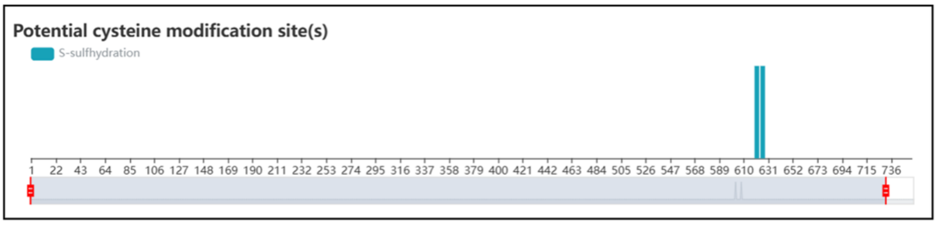


**B**

| ID | Position | Modification | FPR | Peptide |
|---|---|---|---|---|
| NP_001238754.1 | 580 | S-sulfhydration | 0.79% | GKKKQHICHIQGCGK |
| NP_001238754.1 | 585 | S-sulfhydration | 0.10% | HICHIQGCGKVYGKT |

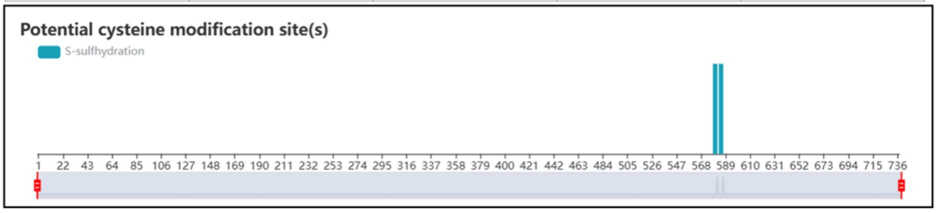


**C**

| ID | Position | Modification | FPR | Peptide |
|---|---|---|---|---|
| NP_612482.2 | 628 | S-sulfhydration | 0.79% | GKKKQHICHIQGCGK |
| NP_612482.2 | 633 | S-sulfhydration | 0.10% | HICHIQGCGKVYGKT |

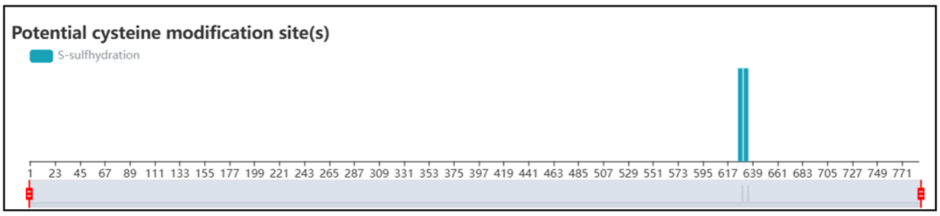


**Figure S27. SP1 isoforms show potential persulfidation sites.** Analysis of amino acid sequence of three SP1 isoforms through pCysMod demonstrating potential persulfidation sites.

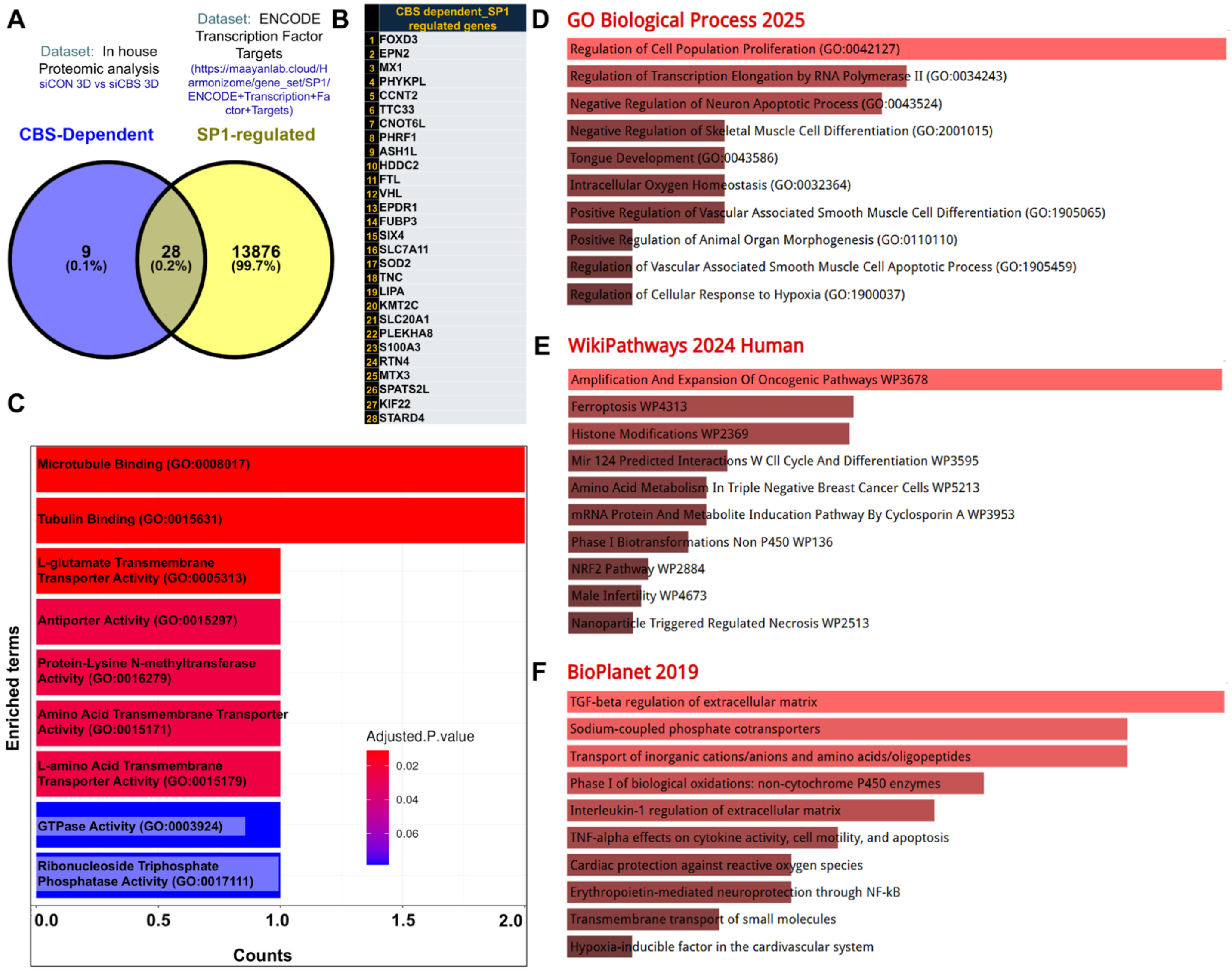


**Figure S28. Identification and functional characterization of CBS-dependent SP1-regulated genes in OvCa spheroids.** (A) Venn diagram showing overlap between CBS-dependent proteins identified in the 3D proteomic siCON vs siCBS dataset and SP1 transcriptional targets (from the Harmonizome ENCODE Transcription Factor Targets dataset). (B) List of overlapping genes representing CBS-dependent SP1-regulated candidates. (C) Gene Ontology (GO) enrichment analysis of overlapping genes performed using TNMplot (Biological Process category, ovarian cancer dataset), highlighting enrichment in cytoskeletal binding, transporter activity, and enzymatic functions. (D) GO Biological Process enrichment analysis using Enrichr, showing significant enrichment of pathways related to cell proliferation, transcriptional regulation, and apoptotic processes. (E) WikiPathways enrichment analysis indicating involvement in oncogenic signaling, ferroptosis, and metabolic regulation. (F) BioPlanet pathway analysis demonstrating enrichment in extracellular matrix regulation, transport processes, and stress response pathways. Colour intensity represents statistical significance (adjusted p-value), and bar length indicates gene counts.

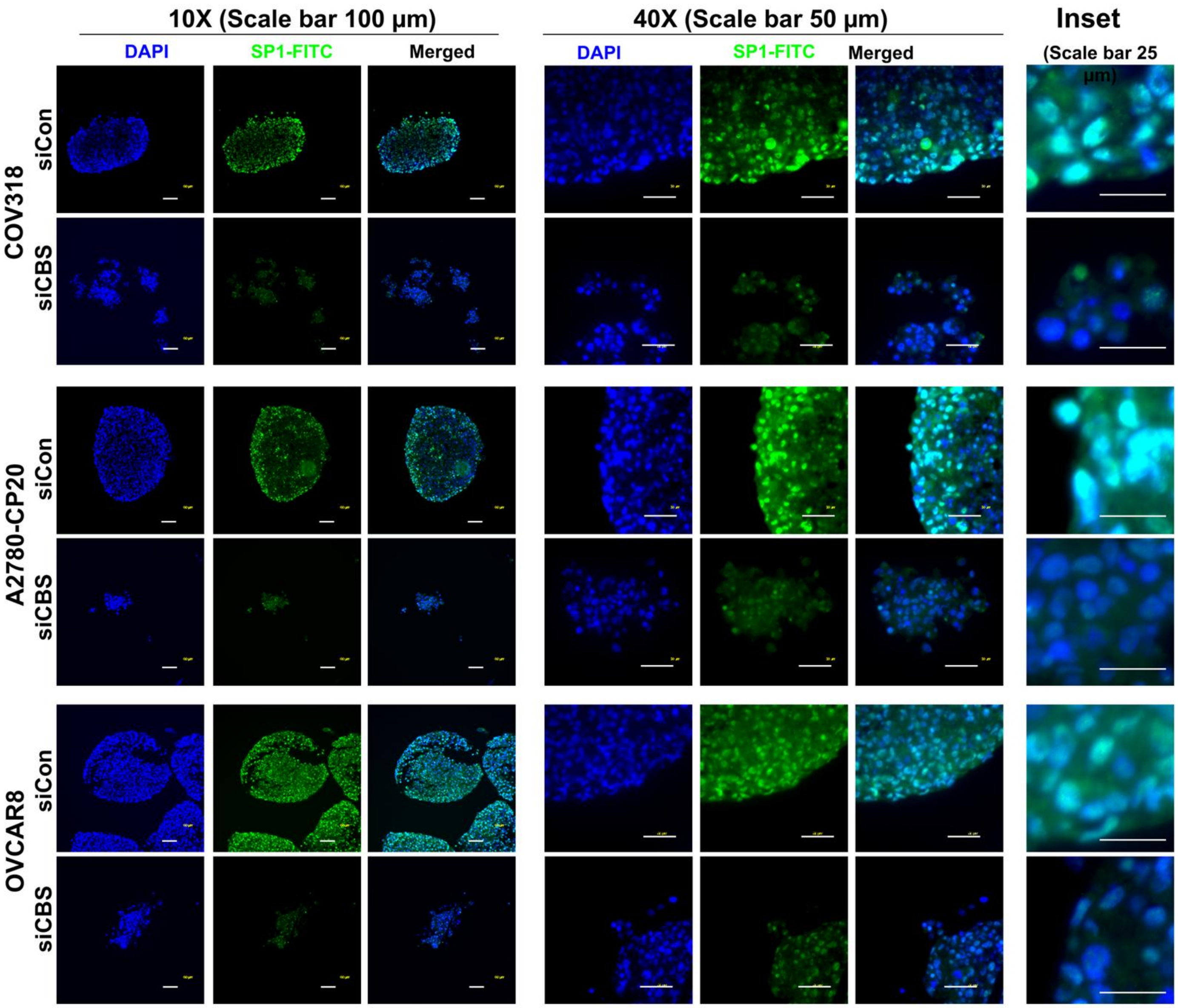


**Figure S29. CBS downregulation causes reduced nuclear presence of SP1 in spheroids.** SP1 localization and expression in OvCa (COV318, A2780-CP20 and OVCAR8) spheroids as a function of CBS expression. Individual channels represent DAPI stained nuclei (Blue) and SP1 at the cellular junctions (Green). CBS knock down resulted in spheroids with reduced mesenchymal attributes.

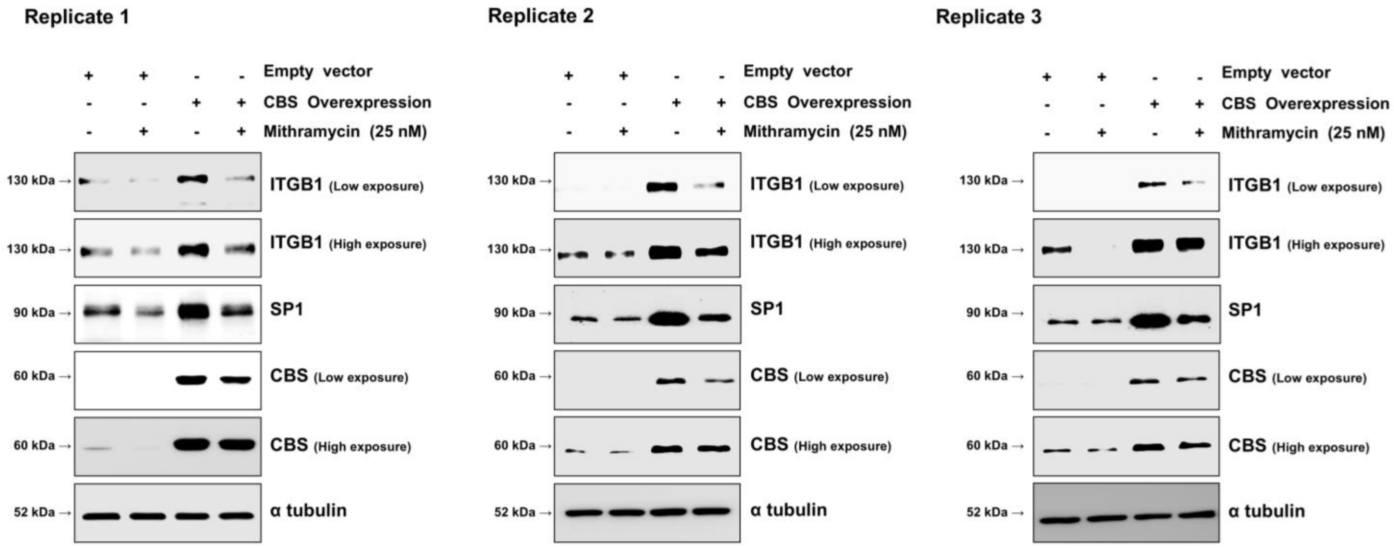


**Figure S30. CBS overexpression enhances SP1 and ITGB1 expression in a SP1-dependent manner in OvCa cells.** Representative immunoblots from three independent biological replicates (Replicates 1–3) showing the effect of CBS overexpression on SP1 and ITGB1 protein levels, and its modulation by SP1 inhibition in OVCAR8 cells. Cells were transfected with empty vector (control) or CBS expression plasmid construct for 24h and treated with the SP1 inhibitor mithramycin (25 μM) where indicated for another 24h. ITGB1 expression (top panels; Shown at both low and high exposure). CBS overexpression increases ITGB1 protein levels compared to control. This induction is attenuated upon mithramycin treatment (25 nM), indicating dependence on SP1 transcriptional activity. SP1 expression: CBS overexpression leads to increased SP1 protein levels, which are reduced upon mithramycin treatment, consistent with inhibition of SP1 function and/or stability. CBS expression: Immunoblot confirms successful overexpression of CBS (low and high exposure shown). α-tubulin: Serves as a loading control across all conditions. Collectively, these data demonstrate that CBS upregulates ITGB1 expression through an SP1-dependent mechanism, supporting a CBS–SP1–ITGB1 signaling axis in OvCa cells.

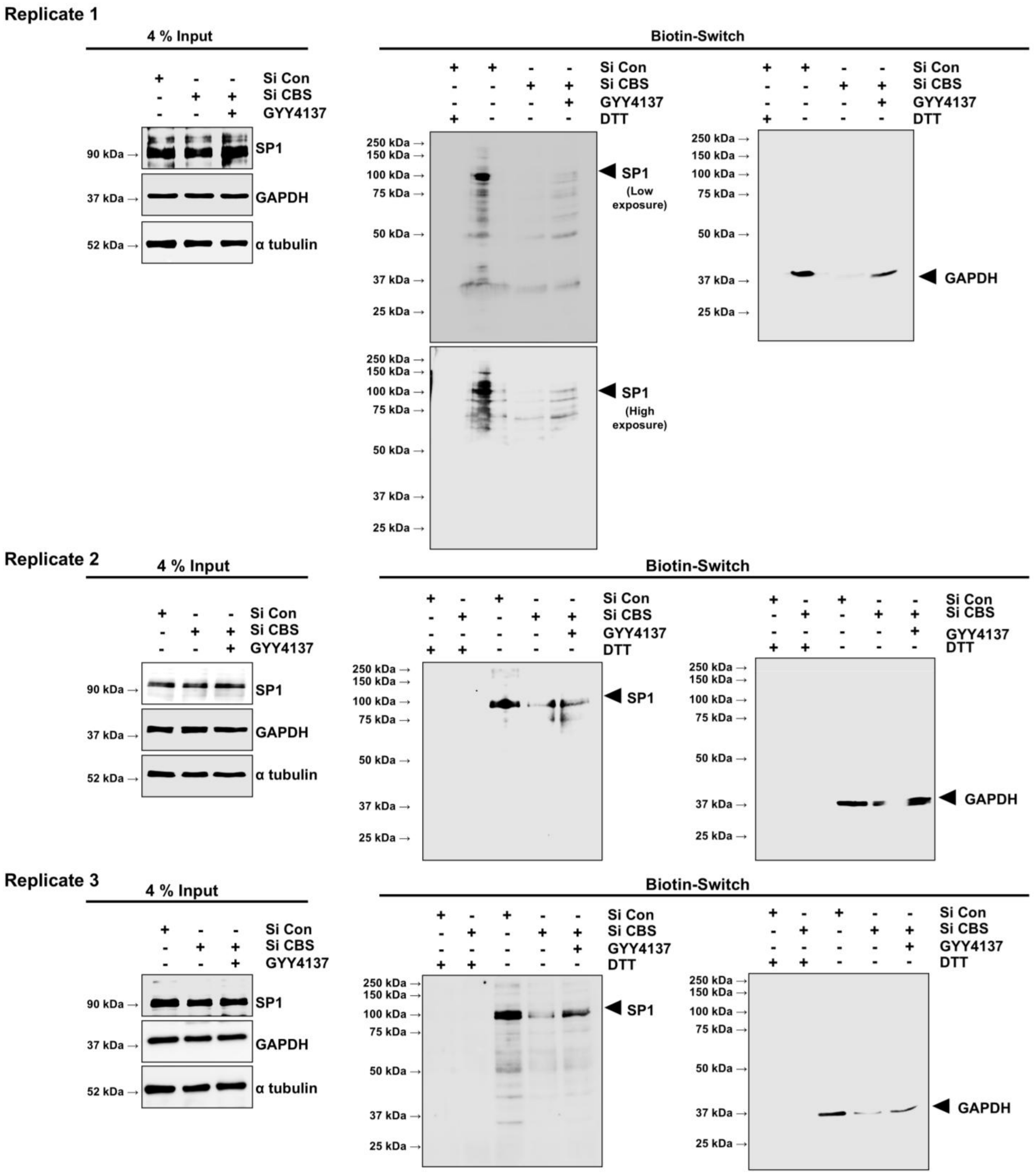


**Figure S31. CBS-derived $H_2S$ promotes SP1 persulfidation in OvCa cells as demonstrated by biotin-switch assay.** Replicates (n=3) of immunoblots showing SP1 persulfidation assessed using a modified biotin-switch assay. OVCAR8 cells were transfected with siCon or siCBS for 96 h. For rescue group, cells were treated with 1 mM GYY4137 for the final 24 h, i.e., 72 h after knockdown. Dithiothreitol (DTT) treatment was used as a negative control to reduce persulfide bonds. Left panels (4% input): Total protein lysates showing SP1 expression levels across conditions, confirming efficient CBS knockdown and comparable protein loading. GAPDH and α-tubulin serve as loading controls. Middle panels (Biotin-switch, SP1): Enrichment of persulfidated SP1 following biotin labeling and pull-down. Both low and high exposure images are shown (Replicate 1) to visualize signal dynamics. CBS silencing (siCBS) markedly reduces SP1 persulfidation compared to control cells, whereas treatment with GYY4137 restores or enhances SP1 persulfidation. DTT treatment abolishes the signal, confirming specificity for persulfide modifications. Right panels (Biotin-switch, GAPDH): Since GAPDH is a known persulfidation target of CBS, probing GAPDH in Biotin switch pull down serves as a positive experimental control.

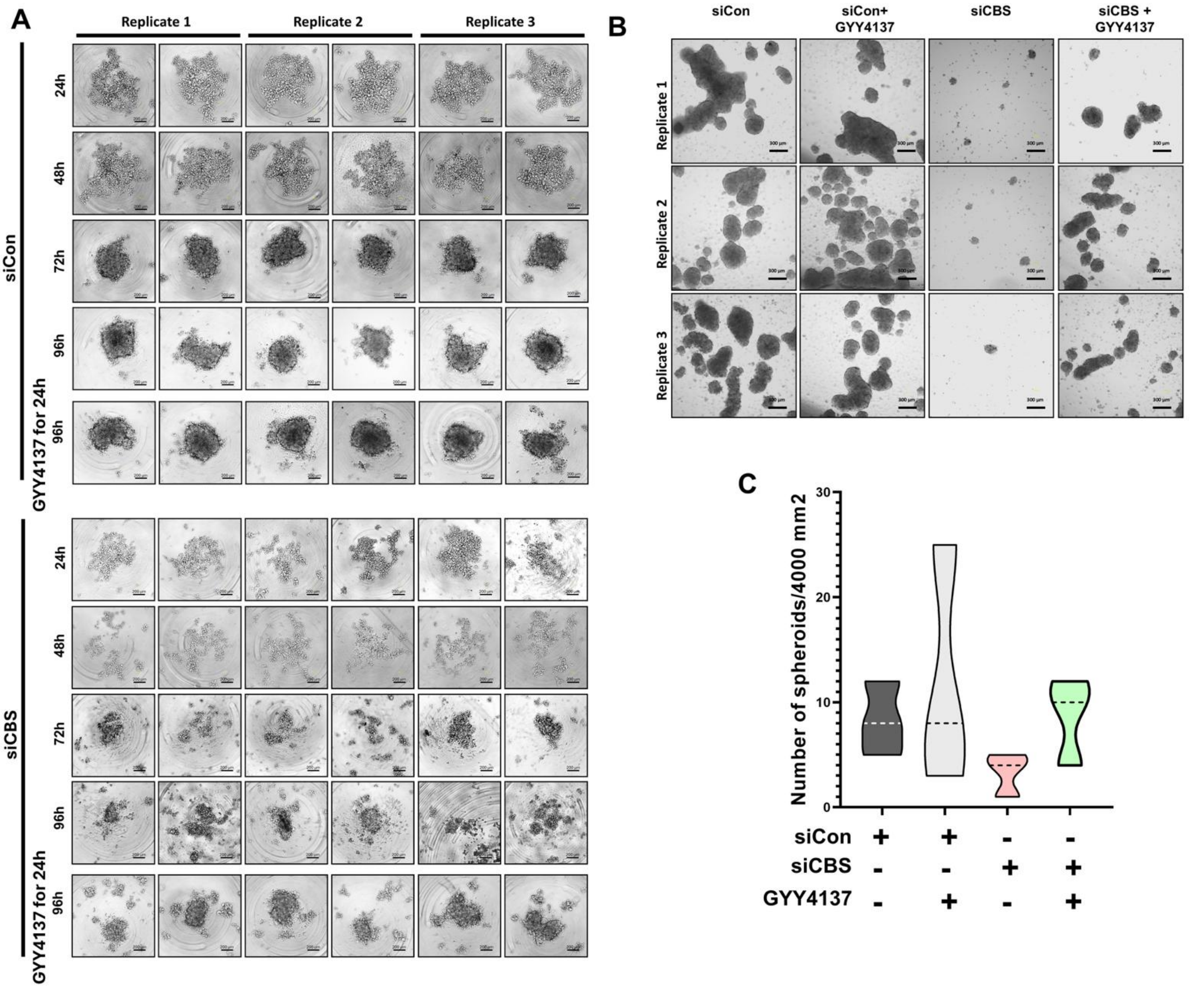


**Figure S32. Phase contrast photomicrographs of spheroids showing attenuation upon CBS silencing and restoration by GYY4137 ($H_2S$ donor) supplementation.** (A) Photomicrographs showing time dependent formation of individual spheroids in siCon, siCBS and siCBS+GYY4137-COV318. (B) Photomicrographs showing number of spheroids in the three treatment groups along with quantification (C). Data from 3 independent biological replicates (n=3)

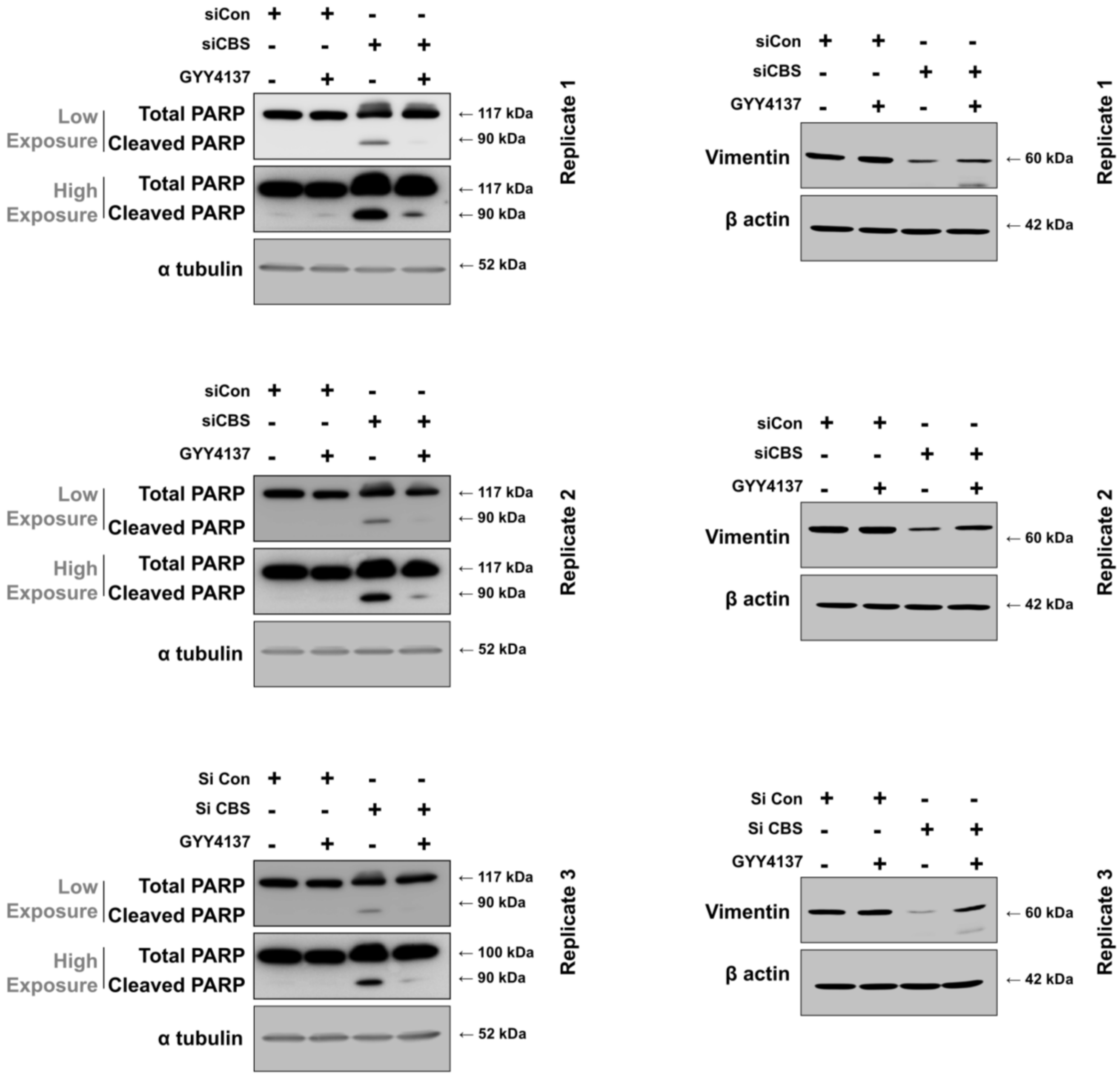


**Figure S33. Restoration of spheroidal viability and invasiveness in CBS silenced COV318 spheroids upon GYY4137 ($H_2S$ donor) supplementation.** Western blots showing three replicates of PARP1 and vimentin in CBS-expressing spheroids, CBS-expressing spheroids supplemented with GYY4137, CBS-silenced spheroids and CBS-silenced spheroids supplemented with GYY4137. GYY4137 ($H_2S$ donor) supplementation attenuated PARP cleavage and restored vimentin expression. Data from 3 independent biological replicates (n=3)

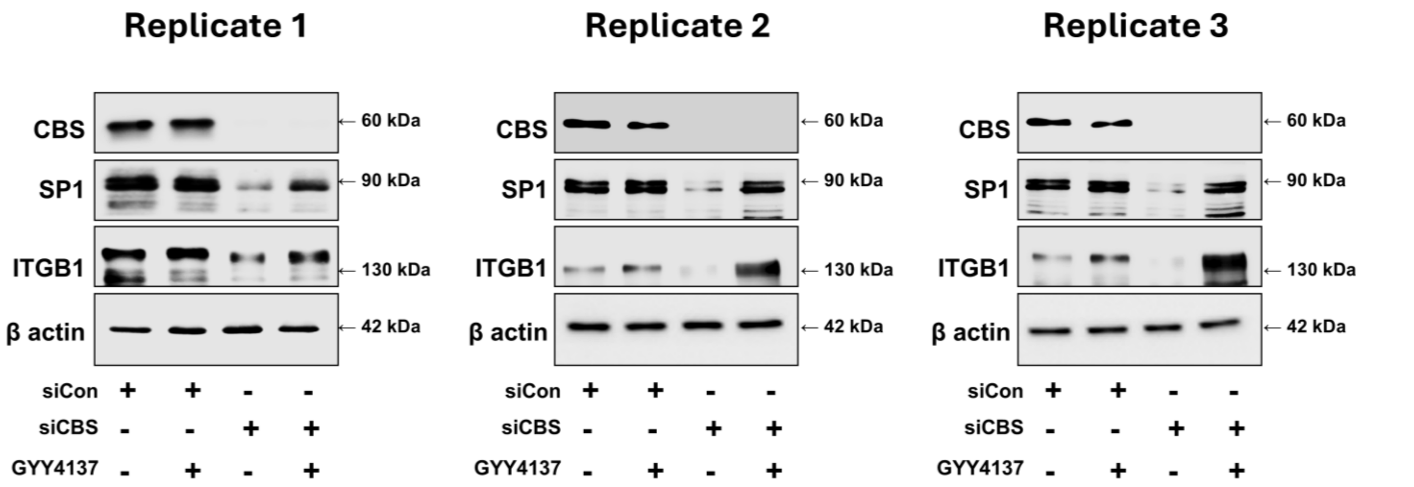


**Figure S34. Restoration of SP1 stability and subsequent ITGB1 expression in COV318 spheroids upon GYY4137 ($H_2S$ donor) supplementation.** Western blots showing three replicates of CBS, SP1 and ITGB1 in CBS-expressing spheroids, CBS-expressing spheroids supplemented with GYY4137, CBS-silenced spheroids and CBS-silenced spheroids supplemented with GYY4137. GYY4137 ($H_2S$ donor) supplementation restored ITGB1 expression through increasing the protein stability of SP1.

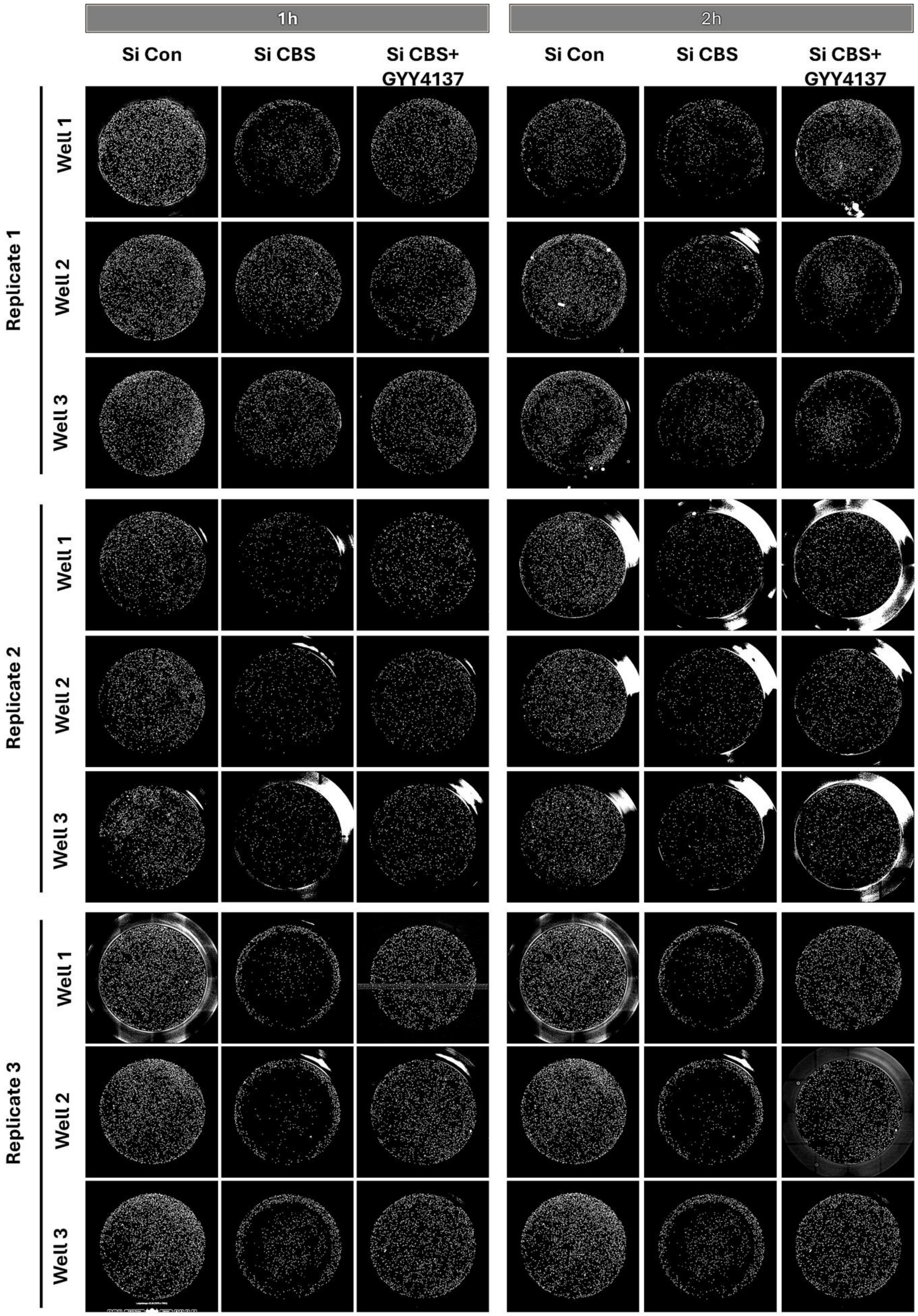


**Figure S35. CBS-mediated ITGB1 downregulation reduced spheroidal cell binding to fibronectin-coated surfaces, which was restored by GYY4137 ($H_2S$ donor) supplementation.** 16-bit stitched images of fibronectin-coated whole wells (96 well plate) plated with labelled CBS-expressing and CBS-silenced spheroidal cells. Data shows reduction in number of adherent cells upon CBS knockdown which was restored upon GYY4137 ($H_2S$ donor) supplementation. Data from 3 independent biological replicates (n=3)

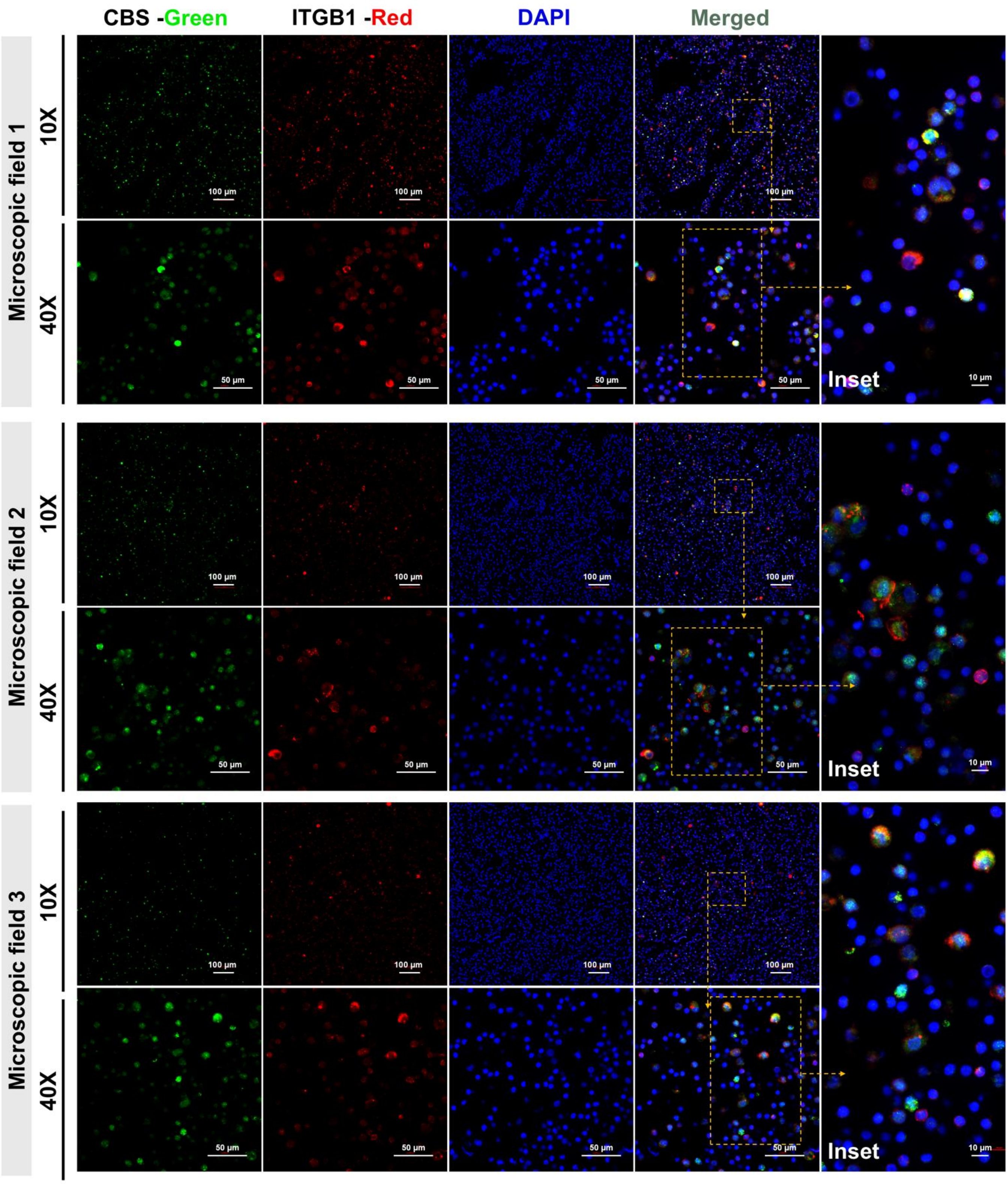


**Figure S36. Expression of CBS and ITGB1 in a low MCA ascites sample (Patient 1).** Immunofluorescence images from three separate microscopic fields (Field 1–3) of the same low MCA ascites sample, with CBS (green), ITGB1 (red), and nuclei (DAPI, blue) staining. The images are shown at 10× and 40× magnifications, with panels and insets.

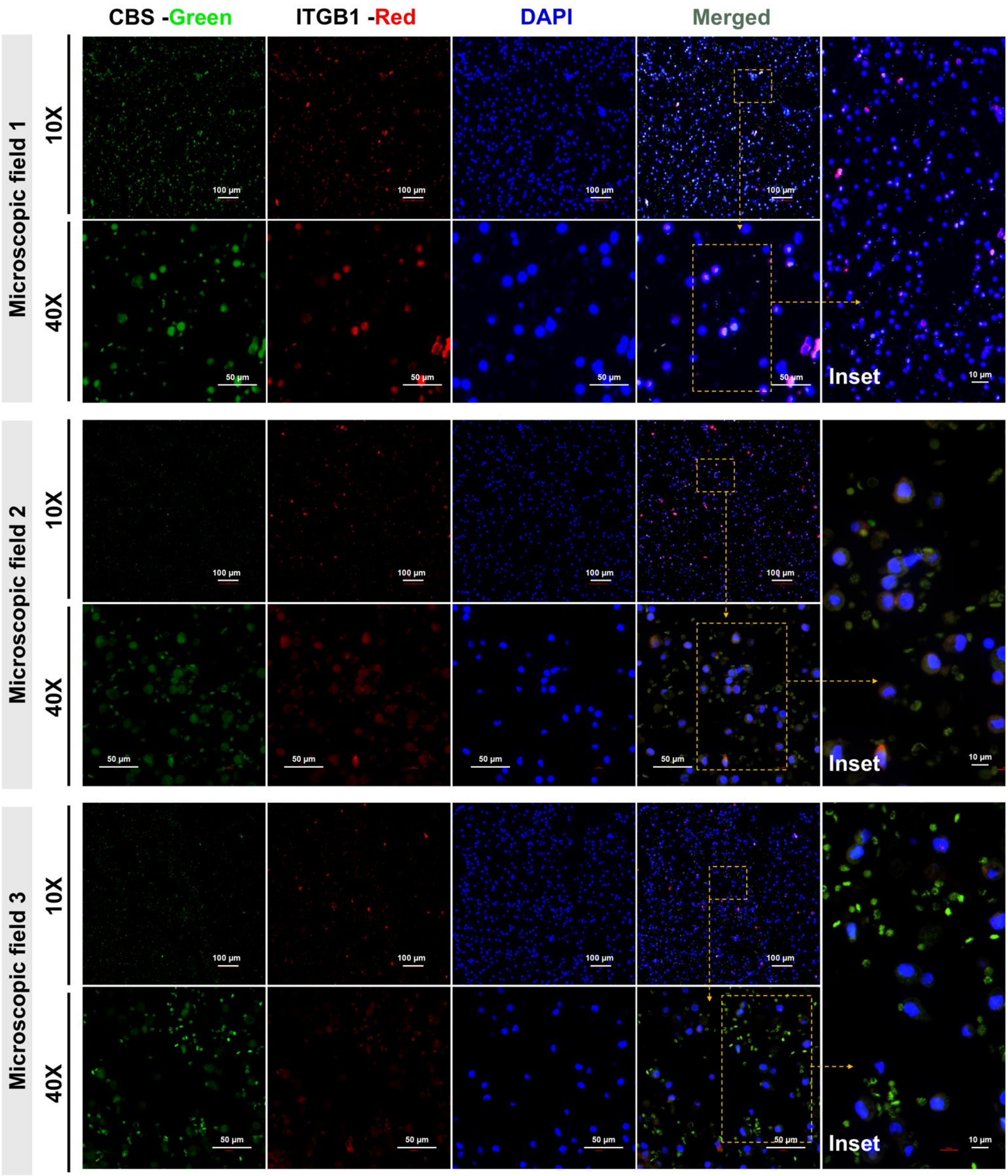


**Figure S37. Expression of CBS and ITGB1 in a low MCA ascites sample (Patient 2).** Immunofluorescence images from three separate microscopic fields (Field 1–3) of the same low MCA ascites sample, with CBS (green), ITGB1 (red), and nuclei (DAPI, blue) staining. The images are shown at 10× and 40× magnifications, with panels and insets.

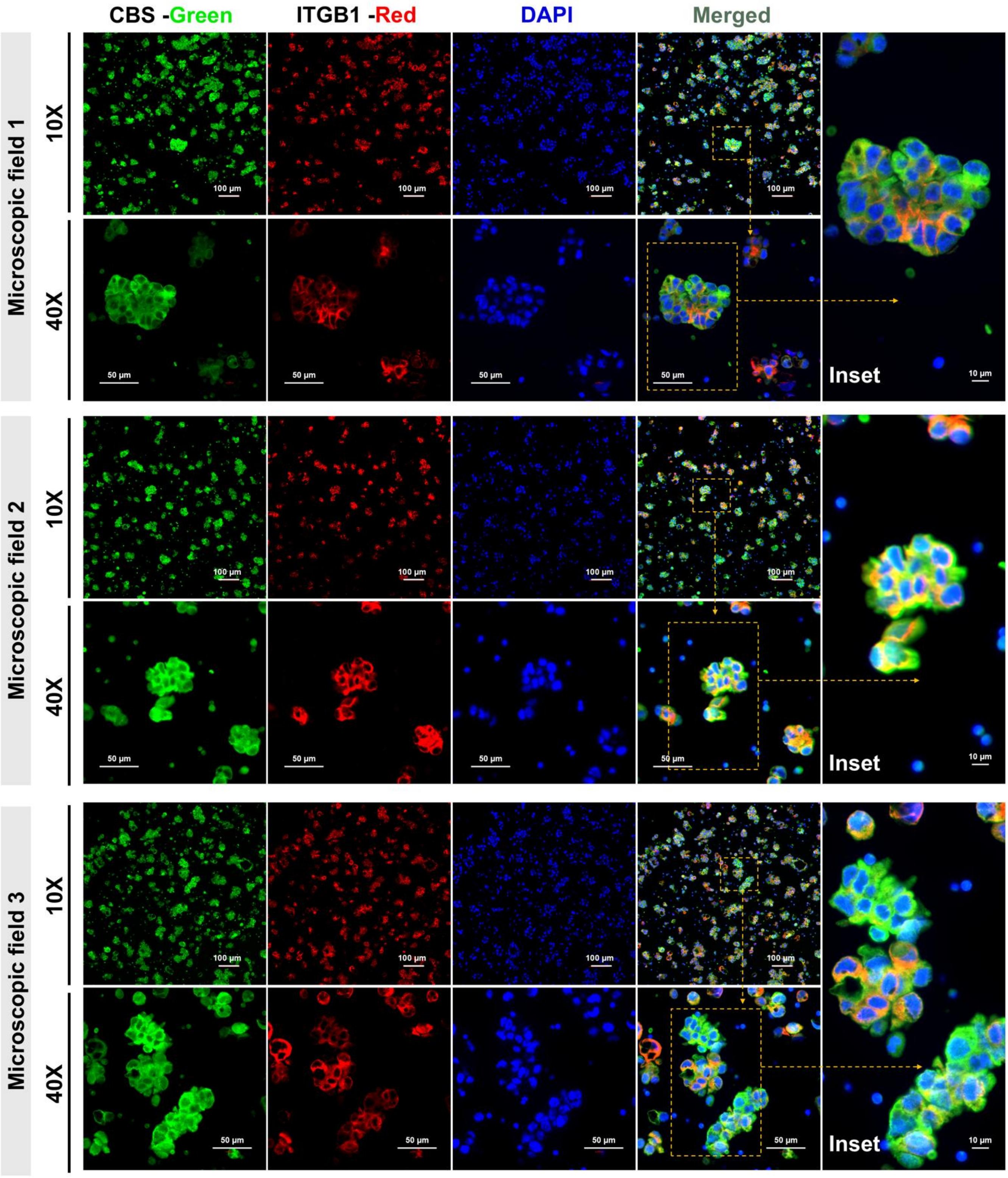


**Figure S38. CBS and ITGB1 expression in a high MCA ascites sample (Patient 3).** Representative immunofluorescence images from three distinct microscopic fields (Field 1–3) of the same patient-derived high MCA ascites sample. The cells were stained with DAPI (blue), CBS (green), and ITGB1 (red). The images are shown at 10× and 40× magnification with merged panels and insets that show spheroidal clusters. In all fields, multicellular aggregates show high CBS and ITGB1 expression.

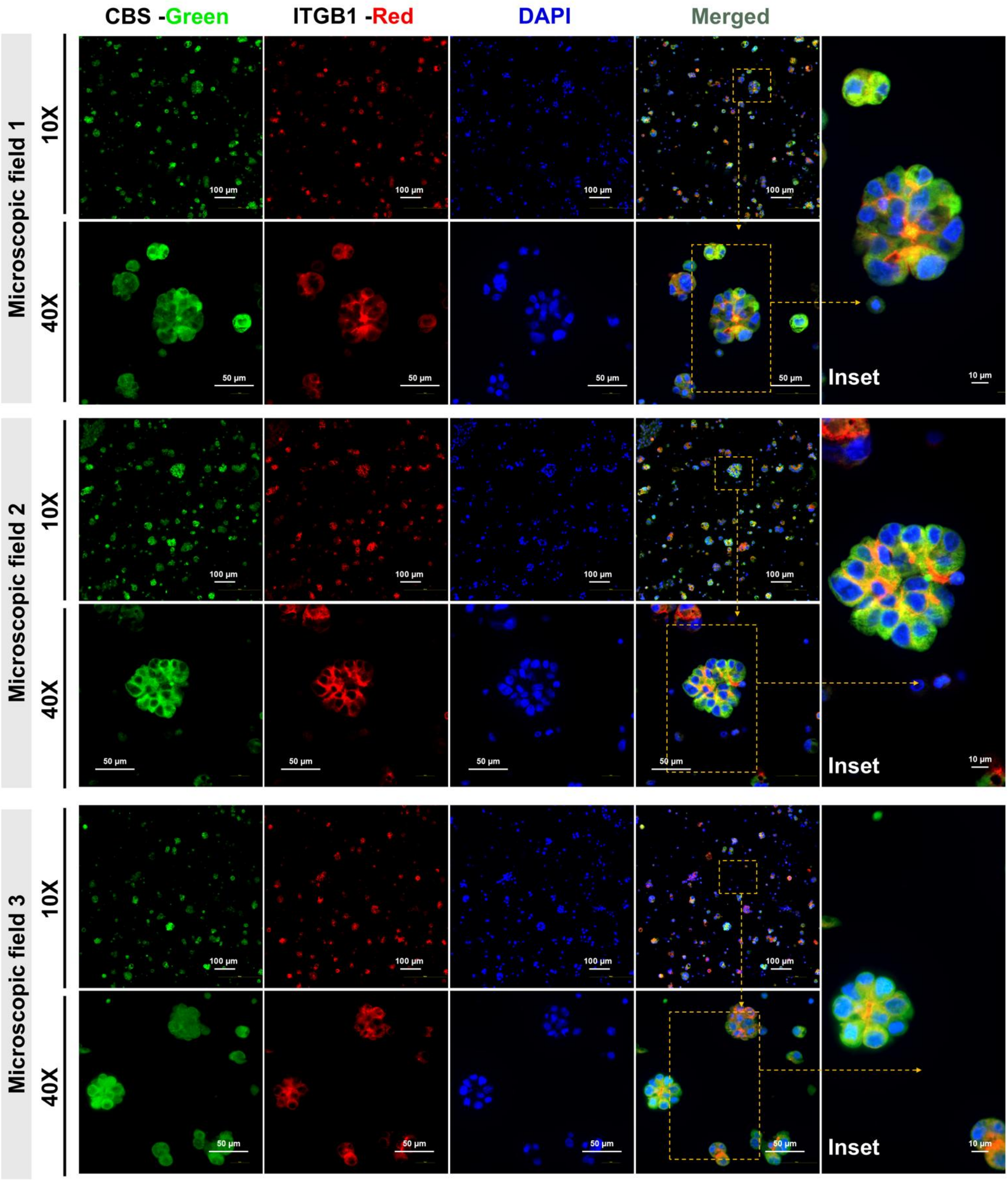


**Figure S39. CBS and ITGB1 expression in a high MCA ascites sample (Patient 4).** Representative immunofluorescence images from three distinct microscopic fields (Field 1–3) of the same patient-derived high MCA ascites sample. The cells were stained with DAPI (blue), CBS (green), and ITGB1 (red). The images are shown at 10× and 40× magnification with merged panels and insets that show spheroidal clusters. In all fields, multicellular aggregates show high CBS and ITGB1 expression.
